# Comprehensive Comparative Genomic revels: *Bacteroides fragilis* is a reservoir of antibiotic resistance genes in the gut microbiota

**DOI:** 10.1101/2022.05.30.494044

**Authors:** Eliane Evanovich, Patricia Jeanne de Souza Mendonça Mattos, João Farias Guerreiro

**Author notes:** Co-authors e-mail addresses. Laboratório de Genética Humana e Médica - Universidade Federal do Pará, Rua Augusto Corrêa, 1 - Guamá, Belém - PA, 66075-110.

## Abstract

*Bacteroides fragilis* are commensal bacteria of the gut microbiota of mammals and may cause severe infection in a susceptible host. Treatment can be cumbersome if multidrug resistant strains are present in the affected tissue. The principal aim of this study was to provide new insights into the genomic properties of *B. fragilis* through different approaches in comparative genomics. Results revealed that the pan-genome is opened, and an intense exchange of genetic material reinforces this inference. The Don complex, responsible for extraintestinal adaptation, is present in all strains, suggesting a crucial role for *B. fragilis* adaptation. CRISPR-Cas system is at 76% of the samples, but it apparently has low accuracy against prophage. Multidrug resistance genes are in 80% of strains. Conjugative transposons and integrative and conjugative elements (ICE) are the main spreaders of genes for antimicrobial resistance. We also reported evidence for horizontal gene transfer (HGT) of antimicrobial resistance genes among the *B. fragilis* strains and Bacteriodales. At least 398 genes are under positive selection, including genes for antimicrobial resistance and transport of toxins and nutrients.

## 1. Introduction

*Bacteroides fragilis* is an obligatory anaerobic Gram-negative bacillus. It is usually commensal in mammals and is critical for immune development and intestinal mucosal integrity [1, 2, 3]. It may become pathogenic in a vulnerable host, being able to invade intra- and extra-abdominal tissues, including blood [4–9]. Recent evidence suggests that enterotoxigenic *B. fragilis* (ETBF) is associated with colorectal cancer development [8–11].

ETBF can produce fragilysin (known as BFT), an endotoxin from the zinc-dependent metalloproteinase family that cleaves E-cadherin in colonocytes [12–16]. The interaction between BFT and intestinal epithelial cells triggers an inflammatory response that increases apoptosis and favors extraintestinal invasion [17]. The *bftP* gene encodes the BFT and has three alleles: *bftP-1, bftP-2*, and *bftP-3* [18]. It is on a pathogenic island termed BfPAI, along with genes for a hypothetical protein and metalloproteinase II *(mpII)* [13, 14, 18–21]. BfPAI of some ETBFs is in the CTn86 transposon [13–14].

Antimicrobial multidrug resistance (AMR) genes are widespread among the B. fragilis strains and hamper antimicrobial therapy [22]. Integrative and conjugative elements (ICE) disseminate AMR genes because they can mobilize plasmids, genomic islands, and other non-conjugative elements [23]. In addition, their genes encode proteins for excision, conjugation (type IV secretion system-T4SS), and transferring DNA sequences [24].

Other genes that improve *B. fragilis* adaptation to intestinal and extraintestinal environments belong to Type VI secretion system (T6SS), capsular lipopolysaccharide biosynthesis locus (LPS locus), and Don PUL complex. T6SS is a protein complex that acts during competition, inhibiting or killing other bacteria or eukaryotic cells [25–27]. The LPS locus has 19 orthologous genes (OG) that control the expression of three capsular polysaccharides (polysaccharides A, B, and C) and may trigger the formation of intra-abdominal abscesses [28–32]. Don PUL complex enables extraintestinal adaptation through uptake of N-linked glycans from fluids [33].

Despite the approaches described above, few studies have investigated the comparative genomics of *B. fragilis*. Thus, this study provides insights into the genomic properties of these bacteria. For this purpose, we analysed 183 strains by applying comparative genomics tools. Some insights obtained were 1) Most genes are related to metabolic processes, mainly carbohydrates and amino acids - a frequent pattern of bacterial gut microbiota; 2) The Don PUL is the only system common to all strains, suggesting that alternative resource capture is decisive for their survival in the gut microbiota; and 3) We also reported evidence of HGT within order Bacteroidales. Hence, this research extends our knowledge of the genomic evolution of B. fragilis.

## 2. Materials and Methods

### Sample

Samples used were available at the National Center for Biotechnology Information - NCBI (https://www.ncbi.nlm.nih.gov/genome/browse/#!/prokaryotes/414/). Genome lengths range from 4.87 to 7.60 megabases (complete, scaffold, and contig), and the minimal coverage was 15X [34] The strains used and their sources are in S1 Table.

### Pan-genome Analysis

Bacterial Pan Genome Analysis tool version 1.3 (BPGA) was used to identify core, accessory, and unique genomes [35]. Gene functions were based on Clusters of Orthologous Groups of proteins (COG) and the Kyoto Encyclopedia of Genes and Genomes (KEGG). Orthologous cluster analysis (default setting) was performed by Usearch9.2.64 [36] under a cut-off value of 95% to generate high-quality clusters [37]. Multiple sequence comparison by log-expectation (MUSCLE) was used to generate alignments and phylogenies [36]. Gnuplot 5.0 was used for plotting graphs [38].

### Prediction of CRISPR and Resistance genes

CRISPR-Cas sequences were estimated by CRISPRRCasFinder (https://crisprcas.i2bc.paris-saclay.fr/CrisprCasFinder/Index) [39]. Antimicrobial resistance genes were obtained from CARD (Comprehensive Antibiotic Resistance Database) (http://arpcard.mcmaster.ca/). CARD was performed under the Resistance Gene Identifier option - RGI (Perfect match equal to 100) [40]. Results with a ‘strict match’ between 95.0 and 99.9 were also analyzed under the Basic Local Alignment Search Tool-BLAST, as suggested by Jia et al. [40].

### Prediction of Mobilome

PhiSpy software was used to predict putative prophages, and ICEFinder was performed to identify ICE and IME (Integrative mobilizable elements) [41–42]. These analyses were restricted to 19 complete genomes because the prophage, ICE, and IME regions are usually long and would be cut into short sequences, such as contigs and scaffolds.

Candidate proteins were searched against the CARD (BLASTp) [43], Virulence factor database (VFDB) [http://www.mgc.ac.cn/VFs/search_VFs.htm; blastp/Protein sequences from VFDB full dataset (setB)] [44], and NCBI server (BLASTp) (https://blast.ncbi.nlm.nih.gov/) [45]. The score values significant were bit-score equal to or higher than 50; E-value ≤ 0.0, and an identity equal or higher than 30% [46]. Mauve under ProgressiveMauve (default setting) [47–48] was used to compare syntenic regions. MobileElementFinder (V1.0.3) (https://cge.cbs.dtu.dk/services/MobileElementFinder/) was performed to predict mobile genetic elements (MGEs) associated with antibiotic and virulence factors (Minimum alignment coverage = 95; minimum sequence identity-90%, maximum truncation-30 bp) [49–50].

### Analysis of Molecular Evolution

Genetic Algorithm Recombination Detection (GARD) was used to detect recombination. Random effect likelihood (FEL), Fast Unconstrained Bayesian Approximation (FUBAR), and Single-likelihood ancestor counting (SLAC) methods implemented by HyPhy 2.5 under the MG94xREV codon model were used to identify sites under positive selection [50–55]. Evidence of episodic positive selection was verified by the mixed-effects model of evolution (MEME) and Branch-site unrestricted statistical test for episodic diversification (BUSTED). The score values were significant whether it is less than or equal to 0.05 (BUSTED, SLAC, FEL, and MEME models), and for the posterior probability if it is ≥ 95% (FUBAR).

## 3. Results and Discussions

### 3.1 Pan-genome

Pan-genome is opened and has about 4232 protein-coding genes, 814 of which belong to the core genome, 3455 are in the set of accessory genomes, and 37 are in the unique genome, on average. Most of which are related to the metabolism of carbohydrates and amino acids (KEGG). Zou et al. [57] described the same pattern in other bacteria of the gut microbiota. According to COG distribution, most of the genes of the flexible gene pool (accessory and unique) are in the ‘Information storage and processing’ category [Translation, ribosomal structure and biogenesis (J); RNA processing and modification (A); Transcription (K); Replication, recombination and repair (L); Chromatin structure and dynamics (B)]. Analysis of USEARCH results indicated that many flexible genes encode truncated proteins. These genes might have been acquired through HGT because they are often are in mobile element regions.

Orthologous genes distributed in their COG and KEGG general categories are in Additional files, S1 Fig. COG distribution in specific categories is in S2 Fig.

KEGG analysis also displayed genes potentially related to human infections, mainly in the flexible genome. These genes often encode proteins with bacterial metabolic functions but may be related to virulence in specific bacteria. For example, the molecular chaperone DNAK protein, encoded by the *dnaK* gene, contributes to stress tolerance and cell growth, but it is also a virulence factor in *Mycobacterium tuberculosis* [58–61]. Fig 1 indicates the distribution of pan-genome across KEGG categories and a table that highlights genes associated with infectious diseases.

**Fig 1.**
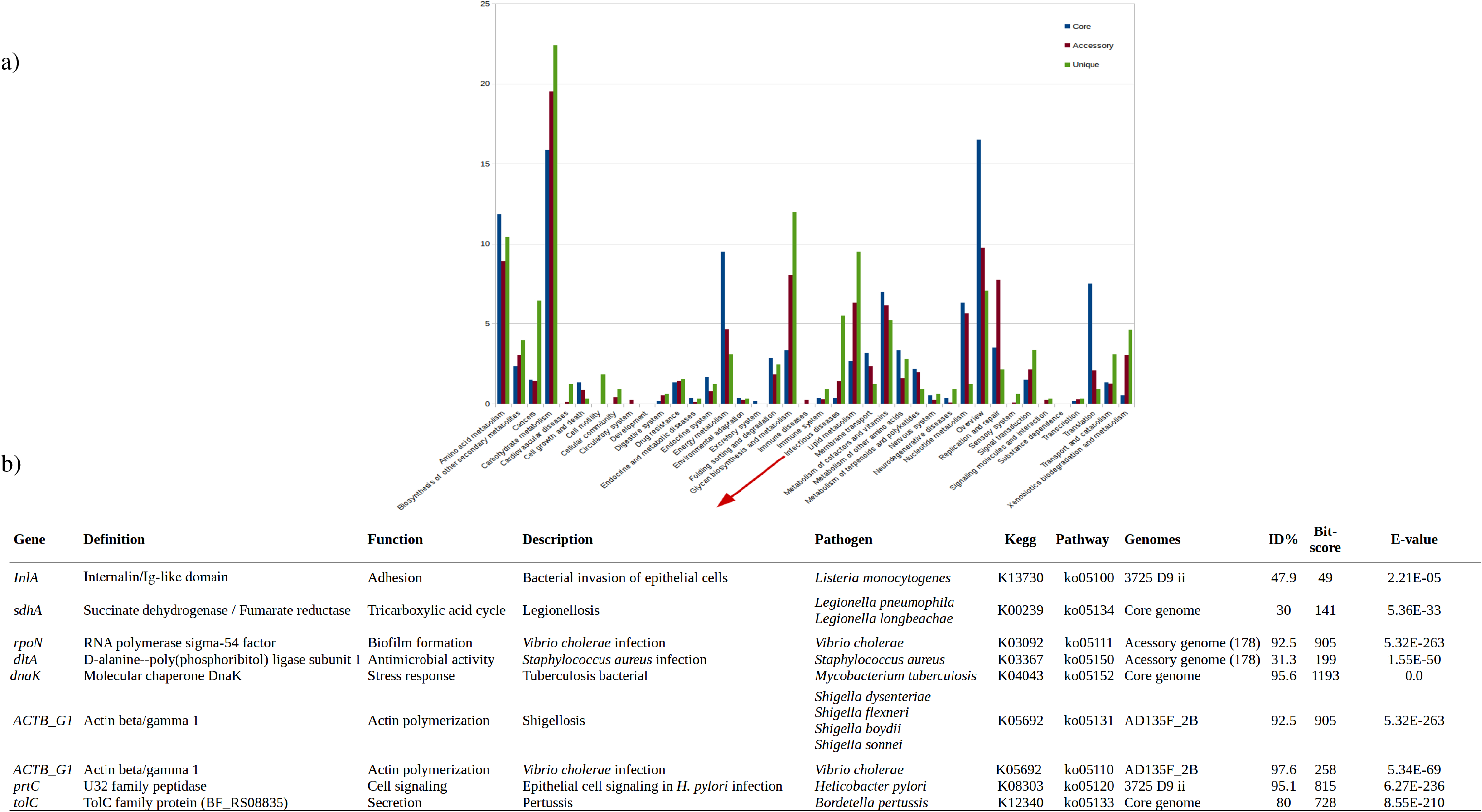
Distribution of orthologous genes into KEGG categories. (a) Frequency of core (in blue), accessory (in red), and unique (green) genes. The arrows point to the table corresponding to the graph bars. (c) Table showing the genes supposedly related to infectious diseases and their features.

Don PUL operon is in all strains, in this way belongs to the core genome. It encodes proteins that enable the deglycosylate N-linked glycans of the gut mucus layer, transferrin, and other glycoproteins [33]. Therefore, the operon is essential for *B. fragilis* adaptation to intra- and extra-intestinal environments. Other genes related to adaptation and virulence are in the accessory genome, such as *bftP*, perforin, LPS locus, and *T6SS* genes.

Forty strains have *bftP-1* allele while *bftP-2* is in eight strains (20793-3, 2-078382-3, 86-5443-2-2, BOB25, CL07T00C01, CL07T12C05, CM13 and DS71). No strain has *bftP-3*. BfPAI is absent in 47 strains described as ETBF [62], suggesting that it was not sequenced or annotated in these strains. Pan-genome prediction and the genomic location of BfPAI are in S2 Table.

The perforin gene is in 80 strains, both ETBFs and NTBFs (non-enterotoxigenic *B. fragilis*). The gene encodes a nonfunctional protein in five strains (1007-1-F #8, 3783N1-2, 3783N2-1, 3976T7, and S13 L11) because of the absence of 139 amino acids from the terminal portion of the protein, which includes the MACPF domain (YGTHVLTDITLGG)-critical for protein activity [61]. Perforin [membrane attack complex/perforin (MACPF) domain] is a bacteriocin that can rupture the plasma membrane of a cell host and might be a virulence factor [63]. It is found mainly in eukaryotes and some bacteria, including *Bacteroides spp*. [64].

LPS locus was described in 638R and NCTC9343 strains [31]. We also reported it in thirteen strains [DCMOUH0042B, 3719 A10, 3986 N3, 3986 T (B) 13, 3986 T (B) 9, AD126T_1B, AD126T_2B, ATCC 25285, BFR_KZ09, CL04T03C20, HMW 615, KLE1257, and KLET1758]. Most of them were isolated from extraintestinal infections (abscess, blood, or tissue). The capsular polysaccharide complex is one of the major virulence factors related to abscess formation [29, 31], and this might explain the source of the strains.

Fifteen complete genomes have GA3 T6SS genetic architecture [27], except the genomes of DCMOUH0067B, DCMSKEJBY0001B, FDAARGOS_763, Q1F2 strains, which belong to the carbapenem-resistant *B. fragilis* group [63]. This finding suggests T6SS is nonessential to *B. fragilis*.

### 3.2 Prediction of CRISPR-Cas system

Results revealed that 76,4% of the genomes have a CRISPR-Cas system. Fifty-eight strains have two CRISPR-Cas types in their genomes. Three types are only in six strains (AF26-6, BFR_KZ08, S14, S38L3, S38L5, and TM08-15). CRISPR-Cas system is absent in most NTBFs, whereas the IIIB type is the most common among ETBFs, corroborating Tajkarimi and Wexler [64]. Monophyletic groups often share the same CRISPR-Cas system. A phylogenetic tree displaying the CRISPR-Cas type of each strain is in S3 Fig.

### 3.3 Antimicrobial Resistance Genes

The most frequent antimicrobial resistance genes in *B. fragilis* are *cepA* [Class A beta-lactamase (EC:3.5.2.6)] and *tetQ* (Tetracycline-resistant ribosomal protection protein). Only 40 strains presented a ‘perfect match’ (100%) for *cepA*, but the results showed high identity values in 158 strains (range from 98 to 99%, ARO: 3003559). Thus, the functionality of *cepA* gene is probable. Thirty-six strains have the *tetQ* gene with a ‘perfect match’, and 125 strains presented a gene with a ‘strict match’ (ARO: 3000191), but likely functional, according to the comparison to the protein described by Lépine et al. [65]. *tetQ* is the most abundant gene for antimicrobials in the human gut microbiota, as reported in a metagenomics analysis [66]. Therefore, the spread of the gene seems advantageous.

*ermF* gene [23S rRNA (Adenine (2058)-N (6))-methyltransferase (EC: 2.1.1.-)] is in 43 strains with a ‘strict match’ (identity from 98 to 99.9%), but their binding sites - essential for protein functionality [67] are unchanged. This gene confers multidrug resistance (macrolide, lincosamide, and streptogramin B) [68] and has a clinical impact on treatment. Antimicrobial resistance genes present in the strains are in S3 Table.

*aadS* [Aminoglycoside 6-nucleotidyltransferase (EC: 2.7.1.95)]*, aphA* [Aminoglycoside 3’-phosphotransferase (EC: 2.7.1.95)], *ccrA* [Subclass B1 beta-lactamase (EC 3.5.2.6)], *OXA-347* [Beta-lactamase class D OXA-209 (EC: 3.5.2.6)], *SUL2* [Dihydropteroate synthase type-2 (EC 2.5.1.15)], and *TEM-1* [Beta-lactamase class A TEM (EC: 3.5.2.6)] genes had ‘perfect matches’, but they were restricted to a few strains.

### 3.4 Mobilome

#### a) Analysis of Prophages

Results indicated the complete genomes have two (BOB25 strain) to fifteen (DCMOUH0067B and FDAASGOS_763 strains) prophage regions. Most virulence-related proteins in these regions had low scores, except for EF-Tu, which had higher values [VFDB database = VFG046475 (F7308_0636) Tu translation elongation factor (EF-Tu) *(Francisella sp*. TX077308) ID = 64%; bit-score = 540, and E-value = e-154]. EF-Tu protein performs various functions in bacteria, including catalyzing the binding of aminoacyl-tRNA to the ribosome, forming a biofilm, and decreasing the host immune response [69]. It is unclear whether it plays a role related to virulence in *B. fragilis*. We found no antimicrobial resistance genes in the prophage region (S4 Table).

#### b) ICEFinder Analysis

The size of ICEs varies from 24.6 kilobase pairs (kb) to 193 kb, while EMIs are smaller (5 kb to 76.8 kb). Genes for antimicrobial resistance, mainly *tetQ*, were identified in ICEs and IMEs. Complete genomes have at least one putative ICE with T4SS, except those from BE1 and Q1F2 with only putative IMEs. S5 Table indicates the genomic location of the ICEs, IMEs and the antibiotic resistance genes found in them.

ICEFinder, Mauve, and NCBI BLAST search (Bit-score = 70088; e-value = 1.020e + 05; ID = 98%) analyses supported the hypothesis of an HGT (horizontal gene transfer) event between the genomic regions of *B. fragilis DCMOUH0085B* (position 2993358 to 3044318) and one from *Butyricimonas faecalis* H184 (Accession no. CP032819.1). The two regions also share the genes for antimicrobial resistance, *AADE* (aminoglycoside 6-adenylyltransferase gene), *ermB* [rRNA adenine N-6-methyltransferase gene], (*APH (3’) - IIIA* {ANT-like pseudogene (9) ANT-like aminoglycoside (9) aminoglycoside nucleotidyltransferase family gene]. HGT among Bacteroidales species was also reported by Coyne et al. [70] and García-Bayona et al. [71].

The results of the Mauve and ICEFinder analyses are in Fig 2 and Table 1.

**Fig 2. Insertion elements found in the strains.** Frequency of insertion elements inferred by MobileElementFinder. The green bar corresponds to the insertion elements with an antimicrobial resistance gene, the red bar denotes insertion elements without antimicrobial resistance gene, and the gray bar shows the untransferred plasmid (repUS2).

**Table 1.** Features of the insertion elements.

| Strain | Accession Number | IS Type | IS Family | Position in Contigs | ID (%) |
| --- | --- | --- | --- | --- | --- |
| **2_1_16** | GG705215.1 | ISBthe1- *nimB* | IS4 | 39206-40521 | 98.9 |
| **320_BFRA** | JVLR01000059.1 | ISBf8 (Tn4555)- *cfxA* | IS4 | 1-1663 | 99.7 |
| **885_BFRA** | JUPJ01000031.1 | IS4351 (Tn4351)- *tetX* | IS30 | 2258-3424 | 99.9 |
| **894_BFRA** | JUOZ01000029.1 | IS4351 (Tn4351)- *tetX* | IS30 | 2258-3424 | 99.9 |
| **915_BFRA** | JUOA01000729.1 | IS4351 (Tn4351)- *tetX* | IS30 | 3147-3947 | 100 |
| **BF8 Consensus_4** | LGTH01000004.1 | IS4351 (Tn4351)- *tetX* | IS30 | 924279-925445 | 99.9 |
|  | LGTH01000004.1 | IS4351 (Tn4351)- *ermF* | IS30 | 922085-923239 | 100 |
| **BF8 Consensus_1** | LGTH01000001.1 | IS1168- *nimB* |  | 612426-612920 | 100 |
| **BF8 Consensus_2** | LGTH01000002.1 | IS942- *cfiA* | IS4 | 1220392-1221141 | 99.1 |
| **CL07T00C01** | AGXM01000001.1 | IS4351 (Tn4351)- *ermF* | IS30 | 582395-583195 | 100 |

A genomic region from the CL03T12C07 strain (position 3350878 to 3378618) is a putative ICE with T4SS that includes a CtnDOT but its *ermF* gene is unfunctional. CtnDOT is a transposon widespread among *Bacteroides spp*. and contains the genes for antimicrobial resistance, *ermF, tetQ, tetX* (flavin-dependent monooxygenase gene) and *tetX1* (flavin-dependent monooxygenase gene) [72–74].

We also identified in DCMSKEJBY0001B strain a putative IME without identified DR that harbours CTn341. This conjugative transposon is typical of *B. fragilis* and carries the *tetQ* gene. CTn341 (or CTn341-like) may also be found in other Bacteroidates, such as *Bacteroides sp*. PHL 2737, *B. xylanisolvens XB1A*, and *Phocaeicola vulgatus* VIC01is. Thus, CTn341 is apparently the major disseminator of the *tetQ* gene within Bacteroidales (S4 Fig).

#### c) Conjugative Transposon

Among the conjugative transposons previously described in *Bacteroides spp*., we reported CTnDOT (aforementioned), CTn86, and CTn341.

We have identified a CTn86-like (Accession no. AY372755.1) in thirteen ETBF strains (2-F-2#5, 3397 N3, 078320–1, 1001285H_161024_D4, AM31-13AC, BFR_KZ09, CL07T00C01, CL07T12C05, CL05T12C13, HMW 615, HMW 616, TL139C_1B, and TL139C_1B2). The *PER* gene, which encodes an ABC permease transporter protein, was excluded from these strains, possibly because of transposon insertion, as described by Franco [13]. CTn86 was also identified in 20793–3, 86-5443-2-2, and BOB25 strains [13, 75–76].

CTn341-like is in 20656-2-1, AF14-14AC, and AF14-26 strains. It does not have all CTn341 genes and attachment sites (*attL* and *attR*) [77]. It has genes for transfer (*traG*, *traI*, *traJ*, and *traM* genes), integration (*int* gene), mobilization (*mobA*, *mobB*, and *mobC*), and excision (*exc* gene) [78–80], besides *tetQ* (S5 Fig).

#### c) Insertion Elements (IS)

ISBf5 (IS1182 family) is the insertion element most common in *B. fragilis* genomes. It has 1830 bp and carries one transposase gene [81].

ISBf1 (IS21 family) [82] is in some strains, but its *cepA* gene is incomplete in eight strains (885_BFR, AD135F_2B, AF32-10, Am47-7, COR2-248-WT-1, HCK-B3, HAP130N_2B, TM08 −15) and absent in AD135F_1B, AD135F_3B, ATCC 25285, and Ds-233 strains. However, they have in their genomes a *cepA* functional with the insertion of IS1224 in their promoter region that increases the *cepA* expression [82–83].

Functional resistance genes were identified in ISBthe1 (*nimB* gene, metronidazole), ISBf8 (*cfxA* gene), and IS4351 (*tetX* gene). ISBf8 and IS4351 are within the non-conjugative transposons (Tn4555 and Tn4351, respectively), and for that reason, they cannot spread their genes.

Therefore, the spread of genes for antibiotic resistance in *B. fragilis* depends on conjugative transposons, as suggested by Whittle et al. [84], Yan, Hall and Jiang [85], and Johnson and Grossman [23], and of ICE with T4SS, as we observed in our results. IS frequency and its genes for antimicrobial resistance are in Fig 3 and the genomic location of the insertion elements is shown in S6 Table.

**Fig 3. Comparison between two putative ICEs with T4SS from *Bacteroides fragilis* DCMOUH0085B and *Butyricimonas faecalis* H184 genomes.** In (a) shown the conserved synteny between these genomic regions; and b) indicates their features in DCMOUH0085B and H184. Antimicrobial resistance genes and their localization in the genomes are indicated in the figure.

### 3.5 Analysis of Molecular Evolution

#### a) Molecular Evolution of Antibiotic Resistance Genes

Results indicated evidence of positive selection in the *cepA* (beta-lactamase gene) and *tetQ* genes. Codon 88 of the *cepA* gene is under positive selection (FUBAR method; Posterior posterior = 0.9878). The substitutions found do not contain any putative ribosome-binding sites indicated by Rogers et al. [86].

*tetQ* alignment displayed 116 variable sites. FUBAR method inferred three codon sites under positive selection: 35 (posterior probability = 0.9676), 237 (posterior probability = 0.9669), and 426 (posterior probability = 0.9623) while MEME suggested episodic positive selection at 354 codon (p-value = 0.003), and 550 (p-value = 0.0464). Comparative analysis with the *tetQ* protein indicated that the four GDP-GTP binding sites are intact, as described by Lépine et al. [65].

#### b) Molecular Evolution in Pan-genome

At least 396 genes are under positive selection (episodic or pervasive). Most of them encode membrane proteins, such as the adenosine triphosphate (atp)-binding cassette (abc) superfamily (*macB* genes), resistance nodulation-division (*rnD*) family (*macA, mdtA, mdtE*, and *mexA* genes), and tonb-dependent receptor (*susC* and *susD* genes).

ATP-binding and RND proteins are efflux pumps that facilitate the elimination of toxins and are related to antimicrobial resistance phenotypes [87–90].

TonB-dependent receptors contribute to the transport and nutrient absorption [90]. Recently, they have been the subject of research for developing vaccines against Gram-negative bacteria because of their cell multiple functions [92–93].

Seven paralogous genes of the Colicin I receptor (*cirA* genes) also had several sites under positive selection. These proteins belong to the TonB-dependent copper receptor family (IPR010100) and perform the transport of nutrients across the periplasmic space [94–95]. Colicins in *Escherichia coli* are toxins that act in intraspecific competition [96–97], but their role in *B. fragilis* is unclear. Yoshizaki et al. [90] compared the genomes of 638R, NCTC9343, and YCH46 and found 52 genes under positive selection. The greater diversity used in our work might explain the difference between the results. The results of Hyphy are in S7 Table.

## 4. Conclusions

HGT is an event of extreme evolutionary relevance for *B. fragilis*. The *tetQ* gene spread was mainly by ICE with T4SS and conjugative transposons, whereas the *cepA* gene was an ancient spread through a mobile element that lost its function. We can describe HGT events not yet reported in previous research. Nonetheless, greater availability of complete genomes is required for the verification and validation of these results. Besides antimicrobial resistance genes, other genes related to efflux pumps facilitating the exit of substances from the cells; transport and absorption of nutrients; and alternative carbon acquisition (DON PUL) might provide adaptive advantages for *B. fragilis*. There is no evidence that *EF-Tu*, *dnaK*, or colicin genes are associated with virulence and need further research to clarify their functions in *B. fragilis*.

## Abbreviation

AMR: Antimicrobial multidrug resistance
BFT: Fragilysin
CRISPR: Clustered regularly interspaced short palindromic repeat
ETBF: Enterotoxigenic *Bacteroides fragilis*
HGT: Horizontal gene transfer
ICE: Integrative and conjugative elements
IME: Integrative mobilizable elements
NTBF: Non-enterotoxigenic *B. fragilis*

## Supporting information

**S1 Table. *Bacteroides fragilis* strains used in this study.**

**S2 Table. Pan-genome prediction of 183 *B. fragilis* strains.**

**S3 Table. Antibiotic resistance genes identified by CARD.**

**S4 Table. Results obtained by PhiSpy compared to the VFDB database.**

**S5 Table. Results obtained by ICEFinder compared to NCBI BLAST and CARD databases.**

**S6 Table. Results obtained by MobileElementFinder.**

**S7 Table. Results obtained by Hyphy software.**

**S1 Fig. Pan-genome prediction.**

**S2 Fig. COG distribution of genes found in the core, accessory and unique genomes.**

**S3 Fig. Pan-genome tree indicating the features of each strain.**

**S4 Fig. Comparison among the CTn341 from DCMSKEJBY0001B, *Phocaeicola vulgatus* VIC01, *Bacteroides sp*. PHL 2737, and *B. xylanisolvens* XB1A.**

**S5 Fig. Comparison among the CTn341 from AF14-14AC, AF14-26 and 20656-2-1 strains.**

## Notes

### Competing Interest Statement

The authors have declared no competing interest.

