## Supplementary material for "Comprehensive Comparative Genomic revels: *Bacteroides fragilis* is a reservoir of antibiotic resistance genes in the gut microbiota": S1

**Table S1.** ***Bacteroides fragilis*** **strains used in this study.**

| Strain | Source | Status | Country | Collection date | Accession |
|---|---|---|---|---|---|
| 143 | Blood | Scaffold | Ecuador | Jan-2017 | NZ_SRKB00000000.1 |
| 1284 | Gut microbiota | Scaffold | USA | 2015 | NZ_PDCX00000000.1 |
| 12905 | Gut microbiota | Scaffold | USA | 2015 | NZ_PDCW00000000.1 |
| 078320-1 | Faeces | Scaffold | USA | Jan-2015 | NZ_PDCV00000000.1 |
| 1001175B_160314_C3 | Faeces | Scaffold | USA | Mar-2016 | NZ_JADMXK000000000.1 |
| 1001175st1_C3 | Faeces | Scaffold | USA | Mar-2016 | NZ_SPGY00000000.1 |
| 1001285H_161024_D4 | Faeces | Scaffold | USA | Oct-2016 | NZ_JADNGU000000000.1 |
| 1007-1-F #10[a] | Faeces | Contig | USA | - | NZ_JGEA00000000.1 |
| 1007-1-F #3[a] | Tissue | Contig | USA | Dec-2007 | NZ_JGEB00000000.1 |
| 1007-1-F #4[a] | Tissue | Contig | USA | Dec-2007 | NZ_JGDK00000000.1 |
| 1007-1-F #5[a] | Tissue | Contig | USA | Dec-2007 | NZ_JGVD00000000.1 |
| 1007-1-F #6[a] | Tissue | Contig | USA | Dec-2007 | NZ_JGVF00000000.1 |
| 1007-1-F #7[a] | Faeces | Contig | USA | - | NZ_JGDL00000000.1 |
| 1007-1-F #8[a] | Faeces | Contig | USA | - | NZ_JGCM00000000.1 |
| 1007-1-F #9[a] | Faeces | Contig | USA | - | NZ_JGEC00000000.1 |
| 1009-4-F #10[a] | Faeces | Contig | USA | - | NZ_JGED00000000.1 |
| 1009-4-F #7[a] | Faeces | Contig | USA | - | NZ_JGEE00000000.1 |
| 14-106904-1 | Appendix | Scaffold | Sweden | 2014 | NZ_MKXL00000000.2 |
| 2_1_16 | Gastrointestinal tract | Scaffold | USA | - | NZ_ACPP00000000.1 |
| 2_1_56FAA | Gastrointestinal tract | Scaffold | USA | - | NZ_ACWI00000000.1 |
| 2-078382-3[a] | Faeces | Contig | USA | 1989 | NZ_LIDV00000000.1 |
| 2-F-2# 4[a] | Tissue | Contig | USA | Dec-2011 | NZ_JGDM00000000.1 |
| 2-F-2# 5[a] | Tissue | Contig | USA | Dec-2011 | NZ_JGCN00000000.1 |
| 2-F-2# 7[a] | Tissue | Contig | USA | Dec-2011 | NZ_JGCO00000000.1 |
| 20656-2-1[a] | Faeces | Contig | USA | 1987 | NZ_LIDU00000000.1 |
| 20793-3[a] | Faeces | Contig | USA | 1987 | NZ_JGEF00000000.1 |
| 3-F-2 #6 | Faeces | Contig | USA | Dec-2011 | NZ_JGDT00000000.1 |
| 320_BFRA | Fluid | Contig | USA | - | NZ_JVLR00000000.1 |
| 321_BFRA | Fluid | Scaffold | USA | - | NZ_JVLQ00000000.1 |
| 322_BFRA | Fluid | Contig | USA | - | NZ_JVLP00000000.1 |
| 3397 N2[a] | Tissue | Contig | USA | 2010 | NZ_JGDN00000000.1 |
| 3397 N3[a] | Tissue | Contig | USA | - | NZ_JGEG00000000.1 |
| 3397 T10[a] | Tissue | Contig | USA | - | JGDN00000000 |
| 3397 T14[a] | Tissue | Contig | USA | - | NZ_JGDO00000000.1 |
| 34-F-2 #13[a] | Tissue | Contig | USA | Dec-2012 | NZ_JGCQ00000000.1 |
| 3719 A10[a] | Tissue | Contig | USA | Jul-2010 | NZ_JGDP00000000.1 |

| | | | | | |
|---|---|---|---|---|---|
| 3719 T6[a] | Tissue | Contig | USA | Jul-2010 | NZ_JGEH00000000.1 |
| 3725 D9 ii[a] | Tissue | Contig | USA | Aug-2010 | JGEI00000000 |
| 3725 D9(v)[a] | Tissue | Contig | USA | Aug-2010 | NZ_JGDQ00000000.1 |
| 3774 T13[a] | Tissue | Contig | USA | Mai-2011 | NZ_JGCR00000000.1 |
| 3783N1-2[a] | Tissue | Contig | USA | Jul-2011 | NZ_JGCS00000000.1 |
| 3783N1-6[a] | Tissue | Contig | USA | Jul-2011 | NZ_JGEU00000000.1 |
| 3783N1-8[a] | R colon | Contig | USA | Jul-2011 | NZ_JGDR00000000.1 |
| 3783N2-1[a] | Tissue | Contig | USA | Jul-2011 | NZ_JGCT00000000.1 |
| 3976T7[a] | Tissue | Contig | USA | Nov-2011 | NZ_JGCU00000000.1 |
| 3976T8[a] | Tissue | Contig | USA | Nov-2011 | NZ_JGDS00000000.1 |
| 3986 N(B)19[a] | Tissue | Contig | USA | Feb-2012 | NZ_JGCW00000000.1 |
| 3986 N(B)22[a] | Tissue | Contig | USA | Feb-2012 | NZ_JGEI00000000.1 |
| 3986 N3[a] | Tissue | Contig | USA | Feb-2012 | NZ_JGVG00000000.1 |
| 3986 T(B)13[a] | Tissue | Contig | USA | Feb-2012 | NZ_JGEJ00000000.1 |
| 3986 T(B)9[a] | Tissue | Contig | USA | Feb-2012 | NZ_JGCX00000000.1 |
| 3986T(B)10[a] | Tissue | Contig | USA | Feb-2012 | NZ_JGCV00000000.1 |
| 3988 T1[a] | Tissue | Contig | USA | Mar-2012 | NZ_JGCZ00000000.1 |
| 3988T(B)14[a] | Tissue | Contig | USA | Mar-2012 | NZ_JGCY00000000.1 |
| 3996 N(B) 6[a] | Tissue | Contig | USA | May-2012 | NZ_JGDA00000000.1 |
| 3998 T(B) 4[a] | Tissue | Contig | USA | May-2012 | NZ_JGDC00000000.1 |
| 3998T(B)3[a] | Tissue | Contig | USA | 2012 | NZ_JGDB00000000.1 |
| 638R | Abdominal abscess | Complete | UK | - | NC_016776.1 |
| 86-5443-2-2[a] | Faeces | Contig | USA | 1997 | NZ_LIDS00000000.1 |
| 885_BFRA | ICU[b] | Contig | USA | - | NZ_JUPJ00000000.1 |
| 894_BFRA | ICU[b] | Contig | USA | - | NZ_JUOZ00000000.1 |
| 8E3_BL_hyb[a] | Blood | Contig | USA | - | NZ_CAEUHN000000000.1 |
| 915_BFRA | Faeces | Contig | USA | - | NZ_JUOA00000000.1 |
| A7 (UDC12-2)[a] | Tissue | Contig | Thailand | - | NZ_JGEK00000000.1 |
| AD126T_1B | Transverse colon | Contig | USA | 2018 | NZ_VOHZ00000000.1 |
| AD126T_2B | Purulent sample | Contig | USA | 2018 | NZ_VOIJ00000000.1 |
| AD135F_1B | Purulent sample | Contig | USA | 2018 | NZ_VOHV00000000.1 |
| AD135F_2B | Faeces | Contig | USA | 2018 | NZ_WCHZ00000000.1 |
| AD135F_3B | Purulent sample | Contig | USA | 2018 | NZ_VOHT00000000.1 |
| AF14-14AC | Purulent sample | Scaffold | China | Oct-2013 | NZ_QRZO00000000.1 |
| AF14-26 | Purulent sample | Scaffold | China | Oct-2013 | NZ_QRZH00000000.1 |
| AF15-10LB | Faeces | Scaffold | China | Nov-2013 | NZ_QRYS00000000.1 |
| AF26-6 | Purulent sample | Scaffold | China | Dec-2013 | NZ_QRTT00000000.1 |
| AF27-10 | Purulent sample | Scaffold | China | Dec-2013 | NZ_QRTQ00000000.1 |
| AF32-10 | Purulent sample | Scaffold | China | Jan-2014 | NZ_QRQH00000000.1 |

| | | | | | |
|---|---|---|---|---|---|
| am_0171 | Purulent sample | Contig | USA | Jan-2015 | NZ_RCXN00000000.1 |
| AM13-18 | Faeces of an elephant | Scaffold | China | Aug-2013 | NZ_QRLH00000000.1 |
| AM15-16 | Clinical isolate | Scaffold | China | Sep-2013 | NZ_QRKP00000000.1 |
| AM16-16A | Faeces | Scaffold | China | Sep-2013 | NZ_QRKL00000000.1 |
| AM17-19 | Faeces | Scaffold | China | Sep-2013 | NZ_QRJX00000000.1 |
| AM18-6 | Faeces | Scaffold | China | Sep-2013 | NZ_QRJE00000000.1 |
| AM26-13LB | Faeces | Scaffold | China | Nov-2013 | NZ_QSLE00000000.1 |
| AM31-13AC | Faeces | Scaffold | China | Dec-2013 | NZ_QSJF00000000.1 |
| AM40-4AC | Faeces | Scaffold | China | Jan-2014 | NZ_QSGL00000000.1 |
| AM47-7 | Faeces | Scaffold | China | Apr-2014 | U30316 |
| ATCC 25285 | Apendix abscess | Scaffold | USA | - | NZ_MTGH00000000.1 |
| B1 (UDC16-1)[a] | Tissue | Contig | USA | - | PDCU01000000 |
| BE1 | Blood | Complete | UK | - | NZ_LN877293.1 |
| BF8 | Infection site | Contig | Hungary | 1985 | NZ_LGTH00000000.1 |
| BFG-1 | Blood | Complete | USA | 1985 | NZ_CP081922.1 |
| BFR_KZ01 | Purulent sample | Contig | Kazakhstan | Dec-2018 | JGDW00000000 |
| BFR_KZ02 | Wound | Contig | Kazakhstan | Nov-2018 | PDCW01000000 |
| BFR_KZ03 | Faeces | Contig | Kazakhstan | Nov-2018 | QSRG01000000 |
| BFR_KZ05 | Faeces | Scaffold | Kazakhstan | Mai-2019 | NZ_JACENG000000000.1 |
| BFR_KZ06 | Faeces | Contig | Kazakhstan | Mai-2019 | NZ_JACFSS000000000.1 |
| BFR_KZ07 | Faeces | Scaffold | Kazakhstan | Jul-2019 | NZ_JACFST000000000.1 |
| BFR_KZ08 | Faeces | Scaffold | Kazakhstan | Aug-2019 | NZ_JACFSU000000000.1 |
| BFR_KZ09 | Faeces | Contig | Kazakhstan | Oct-2019 | NZ_JACFSV000000000.1 |
| BFR_KZ10 | Faeces | Scaffold | Kazakhstan | Oct-2019 | NZ_JACFSW000000000.1 |
| BFR_KZ11 | Faeces | Contig | Kazakhstan | Nov-2019 | NZ_JACFSX000000000.1 |
| BFR_KZ12 | Purulent sample | Contig | Kazakhstan | Nov-2019 | NZ_JACFSY000000000.1 |
| BOB25[a] | Faeces | Complete | Russia | Mar-2013 | NZ_CP011073.1 |
| CCUG4856T | Faeces | Complete | UK | 1955 | NZ_CP036555.1 |
| Plasmid pBF9343 | | | | | NZ_CP036556.1 |
| CFPLTA004_1B | Faeces | Contig | USA | 2018 | NZ_VOHY00000000.1 |
| CL03T00C08 | Faeces | Scaffold | USA | - | NZ_AGXK00000000.1 |
| CL03T12C07 | Faeces | Complete | USA | Mar-2009 | CP072257.1 |
| CL04T03C20 | Faeces | Scaffold | USA | Mar-2009 | NZ_PDCU00000000.1 |
| CL05T00C42 | Faeces | Contig | USA | - | NZ_AKBY00000000.1 |
| CL05T12C13 | Faeces | Scaffold | USA | - | NZ_AGXP00000000.1 |
| CL07T00C01 | Faeces | Scaffold | USA | - | NZ_AGXM00000000.1 |
| CL07T12C05 | Faeces | Scaffold | USA | - | NZ_AGXN00000000.1 |
| CM13 | Faeces | Scaffold | USA | 2015 | NZ_PDCT00000000.1 |

| | | | | | |
|---|---|---|---|---|---|
| COR2-248-WT-1 | Faeces | Contig | Germany | Mar-2018 | NZ_JABAGK000000000.1 |
| DCMOUH0017B | Faeces | Complete | Denmark | Jun-2006 | NZ_CP036539.1 |
| Plasmid pBFO17_1 | | | | | NZ_CP036540.1 |
| Plasmid pBFO17_2 | | | | | NZ_CP036541.1 |
| DCMOUH0018B | Blood | Complete | Denmark | 2011/10/10 | NZ_CP036542.1 |
| plasmid pBFO18_1 | | | | | |
| DCMOUH0042B | Faeces | Complete | Denmark | 2008 | NZ_CP036550.1 |
| Plasmid pBFO42_1 | | | | | NZ_CP036551.1 |
| Plasmid pBFO42_2 | | | | | NZ_CP036552.1 |
| DCMOUH0067B | Faeces | Complete | Denmark | 2009 | NZ_CP036553.1 |
| Plasmid pBFO67_1 | | | | | NZ_CP036543.1 |
| Plasmid pBFO18_2 | | | | | NZ_CP036544.1 |
| plasmid pBFO18_3 | | | | | NZ_CP036545.1 |
| DCMOUH0085B | Faeces | Complete | Denmark | 2010 | NZ_CP037440.1 |
| DCMSKEJBY0001B | Faeces | Complete | Denmark | Aug-2013 | NZ_CP036546.1 |
| Plasmid pBFS01_1 | | | | | NZ_CP036547.1 |
| Plasmid pBFS01_2 | | | | | NZ_CP036548.1 |
| Plasmid pBFS01_3 | | | | | NZ_CP036549.1 |
| DK1985 | Blood | Complete | Denmark | 1985 | NZ_CP044428.1 |
| DS-166[a] | Tissue | Contig | USA | - | NZ_JGDD00000000.1 |
| DS-208[a] | Tissue | Contig | USA | - | NZ_JGDE00000000.1 |
| Ds-233[a] | Tissue | Contig | USA | - | NZ_JGDG00000000.1 |
| DS-71[a] | Tissue | Contig | USA | - | NZ_JGDF00000000.1 |
| FDAARGOS_763 | Faeces | Complete | USA | - | NZ_CP054003 |
| Plasmid unnamed1 | | | | | NZ_CP054001.1 |
| Plasmid unnamed2 | | | | | NZ_CP054002.1 |
| Plasmid unnamed3 | | | | | NZ_CP054004.1 |
| GUT04 | Faeces | Complete | Korea | Aug-2019 | NZ_CP043610.1 |
| Plasmid unnamed1 | | | | | NZ_CP043609.1 |
| HAP130N_1B | Faeces | Contig | USA | 2018 | NZ_WCIJ00000000.1 |
| HAP130N_2B | Faeces | Contig | USA | 2018 | NZ_VOIS00000000.1 |
| HAP130N_3B | Faeces | Contig | USA | 2018 | NZ_WCIE00000000.1 |
| HCK-B3 | Faeces | Contig | China | Apr-2017 | NZ_QRBX00000000.1 |
| HMW 610 | Faeces | Scaffold | USA | - | NZ_AGXQ00000000.1 |
| HMW 615 | Faeces | Scaffold | USA | - | NZ_AGXR00000000.1 |
| HMW 616 | Faeces | Scaffold | USA | - | NZ_ALOB00000000.1 |
| I1345[a] | L colon sample | Contig | USA | Dec-2012 | NZ_JGEW00000000.1 |
| J1101437_171009_F3 | Faeces | Scaffold | USA | Oct-2017 | NZ_JADNMB010000001.1 |
| J-143-4[a] | Tissue | Contig | USA | - | NZ_JGDH00000000.1 |
| J38-1[a] | Tissue | Contig | USA | - | NZ_JGDV00000000.1 |
| JIM10 | Faeces | Contig | Russia | Mar-2013 | NZ_MBRB00000000.1 |

| | | | | | |
|---|---|---|---|---|---|
| KLE1257 | Faeces | Scaffold | USA | - | NZ_LTZL00000000.1 |
| KLE1758 | Blood | Scaffold | USA | - | NZ_LTZK00000000.1 |
| Korea 419[a] | Tissue | Contig | USA | - | JMZX02000000 |
| MC1 | Faeces | Scaffold | USA | - | NZ_CAAKNW000000000.1 |
| MGYG-HGUT-00236 | Blood | Scaffold | UK | - | NZ_CABJEQ000000000.1 |
| MGYG-HGUT-01337 | Blood | Scaffold | UK | - | NZ_CABKOU000000000.1 |
| NCTC9343 | Diarrhea;Abscesses | Contig | UK | - | NZ_UFTH00000000.1 |
| NCTC 9343 | Appendix abscess | Complete | UK | 2017 | NC_003228.3 |
| O:21 | Blood | Scaffold | Sweden | 1995 | NZ_LMAC00000000.1 |
| OF01-1 | Faeces | Scaffold | China | Dec-2013 | NZ_QSDG00000000.1 |
| OF03-7 | Faeces | Scaffold | China | Dec-2013 | SFHX01000000 |
| OF05-11AC | Faeces | Scaffold | China | Jan-2014 | NZ_QSWE00000000.1 |
| OF05-13AC | Faeces | Scaffold | China | Jan-2014 | NZ_QSWB00000000.1 |
| OM02-11 | Faeces | Scaffold | China | Nov-2013 | NZ_QSVT00000000.1 |
| OM02-9 | Faeces | Scaffold | China | Nov-2013 | NZ_QSVD00000000.1 |
| OM04-9BH | Faeces | Scaffold | China | Dec-2013 | NZ_QSUS00000000.1 |
| OM06-1 | Faeces | Scaffold | China | Dec-2013 | NZ_QSUD00000000.1 |
| OM06-30AC | Faeces | Scaffold | China | Dec-2013 | NZ_QSTV00000000.1 |
| Q1F2 | Faeces | Complete | Hong Kong | Feb-2013 | NZ_CP018937.1 |
| Plasmid Q1F2-p1 | | | | | NZ_CP018938.1 |
| Plasmid Q1F2-p2 | | | | | NZ_CP018939.1 |
| S13 L11[a] | Tissue | Contig | USA | Nov-2011 | NZ_JGDI00000000.1 |
| S14[a] | Infection | Complete | Hungary | 2014 | NZ_CP012706.1 |
| S23 R14[a] | Tissue | Contig | USA | Feb-2012 | NZ_JGDX00000000.1 |
| S23L17[a] | Tissue | Contig | USA | Feb-2012 | NZ_JHEF00000000.1 |
| S23L24[a] | Tissue | Contig | USA | Feb-2012 | NZ_JGEL00000000.1 |
| S24L15[a] | Tissue | Contig | USA | Feb-2012 | NZ_JGEM00000000.1 |
| S24L26[a] | Tissue | Contig | USA | Feb-2012 | NZ_JGEN00000000.1 |
| S24L34[a] | Tissue | Contig | USA | Feb-2012 | NZ_JGEO00000000.1 |
| S36L11[a] | Patient sample with septicemia | Contig | USA | Jul-2012 | NZ_JGDJ00000000.1 |
| S36L12[a] | Patient sample with septicemia | Contig | USA | Jul-2012 | NZ_JGEP00000000.1 |
| S36L5[a] | Tissue | Contig | USA | Jul-2012 | NZ_JGEQ00000000.1 |
| S38L3[a] | Healthy biopsy Tissue | Contig | USA | Aug-2012 | NZ_JGEV00000000.1 |
| S38L5[a] | Appendix | Contig | USA | Aug-2012 | NZ_JGER00000000.1 |
| S6L3[a] | Faeces | Contig | USA | Aug-2011 | NZ_JGDY00000000.1 |
| S6L5[a] | Faeces day 7 after weanning | Contig | USA | Aug-2011 | RCXN01000000 |
| S6L8[a] | Purulent sample | Contig | USA | Aug-2011 | NZ_JGES00000000.1 |

| | | | | | |
|---|---|---|---|---|---|
| S6R6[a] | Purulent sample | Contig | USA | Aug-2011 | NZ_JGDZ00000000.1 |
| S6R8[a] | Appendix abscess | Contig | USA | Aug-2011 | NZ_JGVE00000000.1 |
| TF05-31 | Infection site | Scaffold | China | Aug-2013 | NZ_QSSJ00000000.1 |
| TF09-1 | Faeces | Scaffold | China | Feb-2014 | NZ_QSRG00000000.1 |
| TL139C_1B | Faeces | Contig | USA | 2018 | NZ_VOHX00000000.1 |
| TL139C_2B | Faeces | Contig | USA | 2018 | NZ_WCII00000000.1 |
| TM08-15 | Faeces | Scaffold | China | Feb-2014 | NZ_QSOR00000000.1 |
| US326 | Faeces (Healthy Adult) | Scaffold | USA | 2015 | NZ_PDCS00000000.1 |
| YCH46 | Faeces (Healthy Adult) | Complete | Japan | - | NC_006347.1 |
| YCH46 strain 82A12 | Faeces (Healthy Adult) | Contig | Japan | - | NZ_UYXF00000000.1 |

[a] Enterotoxigenic Bacteroides fragilis (ETBF) (Science, 2013).
