## Supplementary material for "Comprehensive Comparative Genomic revels: *Bacteroides fragilis* is a reservoir of antibiotic resistance genes in the gut microbiota": S2

**Table S2. Pan-genome prediction of 183 *Bacteroides fragilis* strains.** The number of genes found in the core genome was 814. The pangenome is open. The interrogation (?) corresponds to genes (*btfP* and *mpII*) or genomic location not detected in strains previously assumed as ETBF. The black filled rectangles correspond to the presence of genes (*btfp, mpII,* and perforin gene) or genomic regions (bfpai and *lps* locus).

| | **Pangenome prediction** | | **BfPAI** | | **Location of BfPAI** | **Perforin** | **LPS locus** |
|---|---|---|---|---|---|---|---|
| **Strain** | **Accessory genes** | **Unique genes** | ***BTFP*** | ***MPII*** | **Accession Number** | | |
| 143 | 3353 | 4 | | | | ■ | |
| 1284 | 3165 | 10 | | | | | |
| 12905 | 3351 | 44 | | | | | |
| 078320-1 | 3013 | 21 | ■ | ■ | NZ_PDCV01000002.1 | ■ | |
| 1001175B_160314_C3 | 3548 | 7 | | | | ■ | |
| 1001175st1_C3 | 3834 | 1 | | | | ■ | |
| 1001285H_161024_D4 | 3316 | 8 | ■ | ■ | NZ_JADNGU010000002.1 | | |
| 1007-1-F #10 | 3933 | 4 | | | | ■ | |
| 1007-1-F #3 | 3930 | 6 | ? | ? | ? | ■ | |
| 1007-1-F #4 | 3798 | 3 | ? | ? | ? | ■ | |
| 1007-1-F #5 | 3931 | 5 | ? | ? | ? | ■ | |
| 1007-1-F #6 | 3950 | 4 | ? | ? | ? | ■ | |
| 1007-1-F #7 | 3977 | 2 | ? | ? | ? | ■ | |
| 1007-1-F #8 | 3990 | 17 | ? | ? | ? | ■ | |
| 1007-1-F #9 | 3959 | 9 | ? | ? | ? | ■ | |
| 1009-4-F #10 | 3368 | 16 | ? | ? | NZ_JGED01000029.1 | ■ | |
| 1009-4-F #7 | 3329 | 10 | ? | ? | NZ_JGEE01000045.1 | ■ | |
| 14-106904-1 | 3520 | 31 | | | | ■ | |
| 2_1_16 | 3584 | 49 | | | | | |
| 2_1_56FAA | 3614 | 53 | | | | ■ | |
| 2-078382-3 | 3425 | 21 | ■ | ■ | NZ_LIDV01000087.1 | ■ | |
| 2-F-2# 4 | 3868 | 27 | ■ | ■ | JGDM01000028.1 | ■ | |
| 2-F-2# 5 | 3671 | 21 | ■ | ■ | NZ_JGCN01000039.1 | ■ | |
| 2-F-2# 7 | 3811 | 24 | ■ | ■ | JGCO01000270.1 | ■ | |
| 20656-2-1 | 3142 | 37 | ■ | ■ | NZ_LIDT01000031.1 | | |
| 20793-3 | 3558 | 34 | ■ | ■ | NZ_LIDU01000032.1 | | |
| 3-F-2 #6 | 3633 | 33 | ? | ? | ? | ■ | |
| 320_BFRA | 3314 | 0 | | | | ■ | |
| 321_BFRA | 3331 | 0 | | | | ■ | |
| 322_BFRA | 3308 | 0 | | | | ■ | |
| 3397 N2 | 3387 | 9 | ■ | ■ | JGDN01000086.1 | | |
| 3397 N3 | 3469 | 4 | ■ | ■ | NZ_JGEG01000073.1 | | |
| 3397 T10 | 4207 | 619 | ■ | ■ | JGCP01002420.1 JGCP01002421.1 | | |
| 3397 T14 | 3550 | 3 | ■ | ■ | NZ_JGDO01000036.1 | | |

| | | | | | |
|---|---|---|---|---|---|
| 34-F-2 #13 | 3580 | 26 | | | JGCQ01000253.1 |
| 3719 A10 | 3484 | 30 | | | NZ_JGDP01000071.1 |
| 3719 T6 | 3371 | 23 | ? | ? | ? |
| 3725 D9 ii | 2768 | 2101 | ? | ? | ? |
| 3725 D9(v) | 3940 | 41 | ? | ? | ? |
| 3774 T13 | 3759 | 30 | | | JGCR01000197.1 |
| 3783N1-2 | 3715 | 15 | ? | ? | ? |
| 3783N1-6 | 3845 | 9 | ? | ? | ? |
| 3783N1-8 | 3800 | 13 | ? | ? | ? |
| 3783N2-1 | 3632 | 6 | ? | ? | ? |
| 3976T7 | 3627 | 6 | ? | ? | ? |
| 3976T8 | 3569 | 157 | ? | ? | ? |
| 3986 N(B)19 | 3398 | 65 | ? | ? | ? |
| 3986 N(B)22 | 3392 | 5 | ? | ? | ? |
| 3986 N3 | 3400 | 0 | ? | ? | ? |
| 3986 T(B)13 | 3410 | 1 | | | JGCR01000040.1 |
| 3986 T(B)9 | 3185 | 12 | ? | ? | ? |
| 3986T(B)10 | 3373 | 9 | ? | ? | ? |
| 3988 T1 | 3528 | 60 | ? | ? | ? |
| 3988T(B)14 | 3451 | 28 | ? | ? | ? |
| 3996 N(B)6 | 3638 | 85 | ? | ? | ? |
| 3998 T(B)4 | 3841 | 26 | ? | ? | ? |
| 3998T(B)3 | 3888 | 49 | ? | ? | ? |
| 638R | 3246 | 56 | | | |
| 86-5443-2-2 | 3633 | 25 | | | NZ_LIDS01000027.1 |
| 885_BFRA | 3345 | 1 | | | NZ_JUPJ01000235.1 |
| 894_BFRA | 3532 | 0 | | | NZ_JUOZ01000165.1 |
| 8E3_BL_hyb | 3096 | 37 | | | |
| 915_BFRA | 3483 | 5 | | | NZ_JUOA01001180.1 |
| A7 (UDC12-2) | 3446 | 11 | | | NZ_JGEK01000083.1 |
| AD126T_1B | 3454 | 0 | | | |
| AD126T_2B | 3455 | 1 | | | |
| AD135F_1B | 3757 | 0 | | | |
| AD135F_2B | 3764 | 5 | | | |
| AD135F_3B | 3753 | 0 | | | |
| AF14-14AC | 3422 | 2 | | | |
| AF14-26 | 3431 | 4 | | | |
| AF15-10LB | 3381 | 11 | | | |
| AF26-6 | 3511 | 11 | | | |
| AF27-10 | 3549 | 19 | | | |
| AF32-10 | 3506 | 3 | | | |
| am_0171 | 3467 | 36 | | | |
| AM13-18 | 3626 | 41 | | | |
| AM15-16 | 3523 | 1 | | | |
| AM16-16A | 3521 | 0 | | | |
| AM17-19 | 3418 | 8 | | | |
| AM18-6 | 3445 | 242 | | | |

| | | | | | | |
|---|---|---|---|---|---|---|
| AM26-13LB | 3355 | 29 | | | ■ | |
| AM31-13AC | 3205 | 49 | ■ | NZ_QSJF01000009.1 | | |
| AM40-4AC | 3394 | 19 | | | | |
| AM47-7 | 2748 | 19 | | | ■ | |
| ATCC 25285 | 3424 | 5 | | | | ■ |
| B1 (UDC16-1) | 3242 | 62 | ■ | JGDU01000228.1 | ■ | |
| BE1 | 3243 | 20 | | | ■ | |
| BF8 | 3164 | 4 | | | ■ | |
| BFG-1 | 3086 | 5 | | | ■ | |
| BFR_KZ01 | 3156 | 37 | | | ■ | |
| BFR_KZ02 | 3133 | 0 | | | ■ | |
| BFR_KZ03 | 3417 | 32 | | | ■ | |
| BFR_KZ05 | 3139 | 44 | | | | |
| BFR_KZ06 | 3205 | 28 | | | ■ | |
| BFR_KZ07 | 3155 | 25 | | | | |
| BFR_KZ08 | 3140 | 18 | | | | |
| BFR_KZ09 | 3000 | 5 | ■ | NZ_JACFSV010000004.1 | | ■ |
| BFR_KZ10 | 3170 | 6 | | | | |
| BFR_KZ11 | 3189 | 14 | | | | |
| BFR_KZ12 | 3278 | 0 | | | | |
| BOB25 | 3084 | 13 | ■ | NZ_CP011073.1 | ■ | |
| CCUG4856T | 3447 | 1 | | | | |
| CFPLTA004_1B | 3856 | 102 | | | ■ | |
| CL03T00C08 | 3354 | 4 | | | | |
| CL03T12C07 | 3352 | 7 | | | | |
| CL04T03C20 | 3257 | 2 | | | | ■ |
| CL05T00C42 | 3380 | 3 | ■ | NZ_JH724188.1 | | |
| CL05T12C13 | 3367 | 2 | ■ | NZ_JH724193.1 | | |
| CL07T00C01 | 3553 | 2 | ■ | NZ_JH724206.1 | ■ | |
| CL07T12C05 | 3562 | 2 | ■ | NZ_JH724218.1 | ■ | |
| CM13 | 3228 | 43 | ■ | NZ_PDCT01000007.1 | | |
| COR2-248-WT-1 | 3283 | 2 | | | ■ | |
| DCMOUH0017B | 3410 | 48 | | | | |
| DCMOUH0018B | 3368 | 25 | | | ■ | |
| DCMOUH0042B | 3265 | 19 | | | | |
| DCMOUH0067B | 3220 | 87 | | | | |
| DCMOUH0085B | 3378 | 53 | | | | |
| DCMSKEJBY0001B | 3312 | 29 | | | | |
| DK1985 | 3311 | 32 | | | | |
| DS-166 | 3509 | 10 | ■ | NZ_JGDD01000077.1 NZ_JGDD01000078.1 | | |
| DS-208 | 3328 | 116 | ■ | NZ_JGDE01000059.1 NZ_JGDE01000060.1 | | |
| Ds-233 | 3311 | 67 | ■ | NZ_JGDG01000407.1 | | |
| DS-71 | 3285 | 39 | ■ | NZ_JGDF01000051.1 | | |
| FDAARGOS_763 | 3429 | 56 | | | ■ | |

| | | | | | |
|---|---|---|---|---|---|
| GUT04 | 3577 | 3 | | | |
| HAP130N_1B | 3438 | 10 | | | |
| HAP130N_2B | 3424 | 0 | | | |
| HAP130N_3B | 3433 | 2 | | | |
| HCK-B3 | 3484 | 39 | | | |
| HMW 610 | 3512 | 89 | | | |
| HMW 615 | 3355 | 43 | | | NZ_JH815492.1 |
| HMW 616 | 3458 | 103 | | | NZ_JH815524.1 |
| I1345 | 3585 | 8 | ? | ? | NZ_JGEW01000032.1 |
| J-143-4 | 3648 | 65 | | | NZ_JGDH01000029.1<br>NZ_JGDH01000030.1 |
| J38-1 | 3381 | 35 | | | NZ_JGDV01000110.1<br>NZ_JGDV01000028.1 |
| JIM10 | 3135 | 19 | | | |
| J1101437_171009_F3 | 3353 | 137 | | | |
| KLE1257 | 3822 | 0 | | | |
| KLE1758 | 3822 | 0 | | | |
| Korea 419 | 3314 | 43 | | | |
| MC1 | 3245 | 2 | | | |
| MGYG-HGUT-00236 | 3397 | 1 | | | |
| MGYG-HGUT-01337 | 3323 | 2 | | | |
| NCTC 9343 | 3365 | 13 | | | |
| NCTC9343 | 3357 | 15 | | | |
| O:21 | 3097 | 1 | | | |
| OF01-1 | 3334 | 32 | | | |
| OF03-7 | 2755 | 11 | | | |
| OF05-11AC | 3640 | 1 | | | |
| OF05-13AC | 3580 | 1 | | | |
| OM02-11 | 3240 | 25 | | | |
| OM02-9 | 3236 | 3 | | | |
| OM04-9BH | 3506 | 4 | | | |
| OM06-1 | 3353 | 0 | | | |
| OM06-30AC | 3349 | 0 | | | |
| Q1F2 | 3209 | 38 | | | |
| S13 L11 | 3198 | 56 | ? | ? | ? |
| S14 | 2875 | 6 | | | ? |
| S23R14 | 3411 | 17 | ? | ? | ? |
| S23L17 | 3635 | 16 | ? | ? | ? |
| S23L24 | 3630 | 19 | ? | ? | ? |
| S24L15 | 3552 | 8 | ? | ? | ? |
| S24L26 | 3562 | 3 | ? | ? | ? |
| S24L34 | 3552 | 13 | ? | ? | ? |
| S36L11 | 4090 | 12 | ? | ? | ? |
| S36L12 | 4073 | 11 | ? | ? | ? |
| S36L5 | 4087 | 22 | ? | ? | ? |
| S38L3 | 3267 | 12 | ? | ? | ? |
| S38L5 | 3293 | 7 | ? | ? | ? |
| S6L3 | 3465 | 1 | ? | ? | ? |

| | | | | | | |
|---|---|---|---|---|---|---|
| S6L5 | 3569 | 5 | ? | ? | ? | |
| S6L8 | 3594 | 10 | ? | ? | ? | |
| S6R6 | 3599 | 12 | ? | ? | ? | |
| S6R8 | 3579 | 5 | ? | ? | NZ_JGVE01000096.1 | |
| TF05-31 | 3272 | 17 | | | | |
| TF09-1 | 3486 | 11 | [redacted] | | NZ_QSRG01000006.1 | [redacted] |
| TL139C_1B | 3551 | 0 | [redacted] | | NZ_VOHX01000001.1 | |
| TL139C_2B | 3555 | 5 | [redacted] | | NZ_WCII01000001.1 | |
| TM08-15 | 3397 | 5 | | | | |
| US326 | 3255 | 35 | | | | |
| YCH46 | 3572 | 53 | | | | [redacted] |
| YCH46 strain 82A12 | 3289 | 52 | | | | |
