## Supplementary material for "Comprehensive Comparative Genomic revels: *Bacteroides fragilis* is a reservoir of antibiotic resistance genes in the gut microbiota": S3

**Table S3. Antibiotic resistance genes identified by CARD.** 100% % Identity of matching region is in black, and 96-99%% Identities are in grey. The asterisk indicates that *cfiA* is a pseudogene.

| **Strain** | ***aadS*** | ***aphA*** | ***cepA*** | ***cfiA*** | ***cfxa2*** | ***cfxa3*** | ***ermB*** | ***ermF*** | ***mef (EN2)*** | ***OXA-347*** | ***sul2*** | ***TEM-1*** | ***tetQ*** | ***tetX*** |
|---|---|---|---|---|---|---|---|---|---|---|---|---|---|---|
| 143 | | | grey | | | | | grey | | | | | black | |
| 1284 | | | grey | | | | | | | | | | grey | |
| 12905 | | | grey | | | | | | | | | | black | |
| 078320-1 | | | grey | | | | | | | | | | | |
| 1001175B_160314_C3 | | | black | | black | | | grey | | | | grey | grey | |
| 1001175st1_C3 | | | grey | | black | | | grey | | | | grey | grey | |
| 1001285H_161024_D4 | | | grey | | | | | | | | | | black | |
| 1007-1-F #10 | | | black | | | | | | | | | | black | |
| 1007-1-F #3 | | | black | | | | | | | | | | black | |
| 1007-1-F #4 | | | black | | | | | | | | | | black | |
| 1007-1-F #5 | | | black | | | | | | | | | | black | |
| 1007-1-F #6 | | | black | | | | | | | | | | black | |
| 1007-1-F #7 | | | black | | | | | | | | | | black | |
| 1007-1-F #8 | | | black | | | | | | | | | | black | |
| 1007-1-F #9 | | | black | | | | | | | | | | grey | |
| 1009-4-F #10 | | | grey | | | | | | | | | | grey | |
| 1009-4-F #7 | | | grey | | | | | | | | | | grey | |
| 14-106904-1 | | | grey | | | black | | | | | black | | grey | |
| 2_1_16 | | | black | | | | | | | | | | grey | |
| 2_1_56FAA | | | grey | | | | | grey | | | | | grey | |
| 2-078382-3 | | | grey | | | | | | | | | | | |
| 2-F-2# 4 | | | grey | | | | | | grey | | | | grey | |
| 2-F-2# 5 | | | grey | | | | | | | | | | grey | |
| 2-F-2# 7 | | | grey | | | | | | grey | | | | grey | |
| 20656-2-1 | | | grey | | | | | | | | | | grey | |
| 20793-3 | | | grey | | | | | | grey | | | | black | |
| 3-F-2 #6 | | | grey | | | | | grey | | | | | | |
| 320_BFRA | | | grey | | | black | | | | | | | grey | |
| 321_BFRA | | | grey | | | black | | | | | | | grey | |
| 322_BFRA | | | grey | | | black | | | | | | | grey | |
| 3397 N2 | | | grey | | | | | | | | | | grey | |
| 3397 N3 | | | grey | | | | | | | | | | grey | |
| 3397 T10 | | | grey | | | | | | | | | | grey | |
| 3397 T14 | | | grey | | | | | | | | | | grey | |
| 34-F-2 #13 | | | grey | | | | | | | | | | grey | |
| 3719 A10 | | | grey | | | | | | | | | | grey | |
| 3719 T6 | | | grey | | | | | | grey | | | | black | |
| 3725 D9 ii | | | grey | | | | | | | | | | grey | |
| 3725 D9(v) | | | grey | | | | | | | | | | black | |

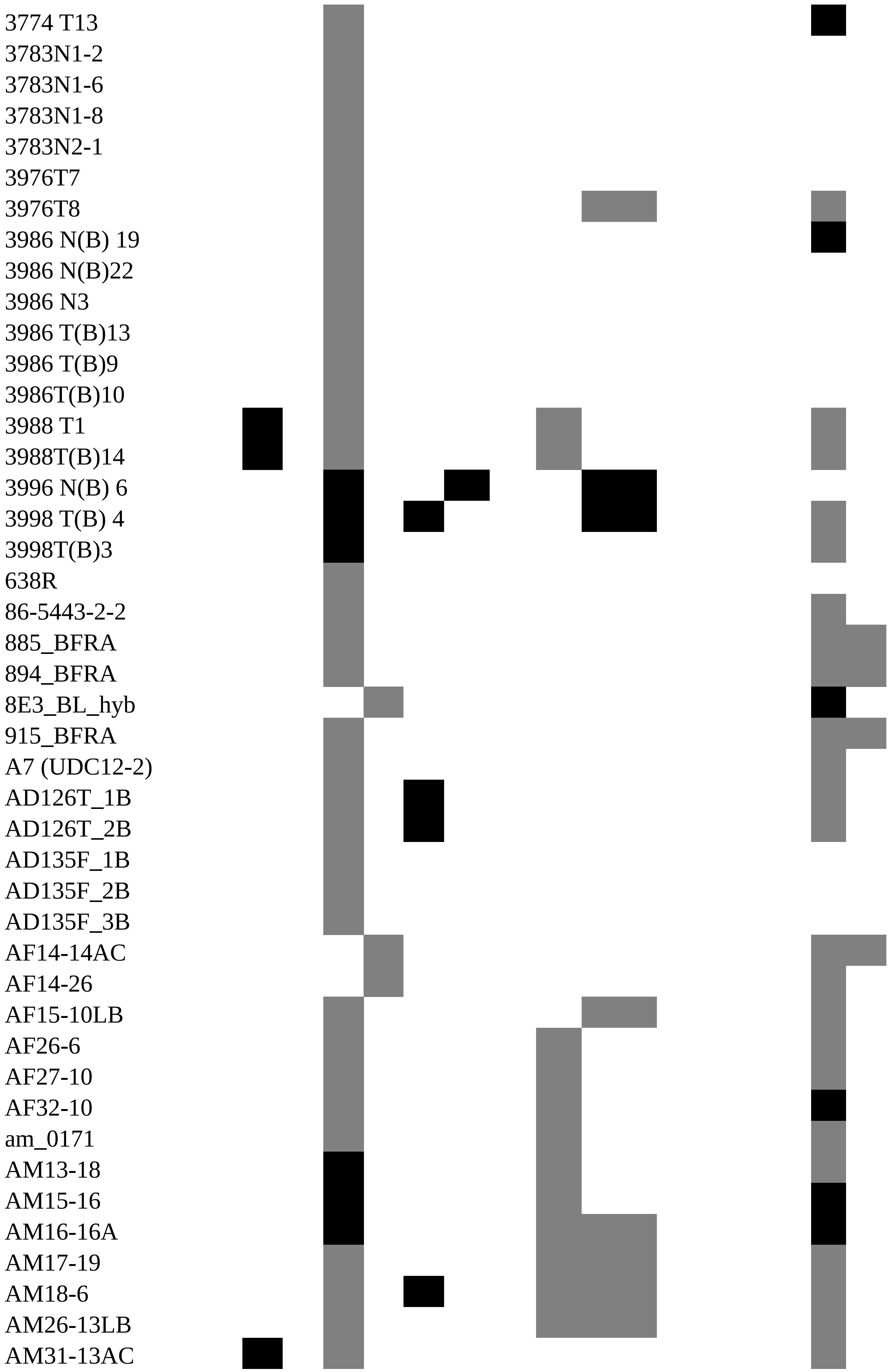
3774 T13
3783N1-2
3783N1-6
3783N1-8
3783N2-1
3976T7
3976T8
3986 N(B) 19
3986 N(B)22
3986 N3
3986 T(B)13
3986 T(B)9
3986T(B)10
3988 T1
3988T(B)14
3996 N(B) 6
3998 T(B) 4
3998T(B)3
638R
86-5443-2-2
885_BFRA
894_BFRA
8E3_BL_hyb
915_BFRA
A7 (UDC12-2)
AD126T_1B
AD126T_2B
AD135F_1B
AD135F_2B
AD135F_3B
AF14-14AC
AF14-26
AF15-10LB
AF26-6
AF27-10
AF32-10
am_0171
AM13-18
AM15-16
AM16-16A
AM17-19
AM18-6
AM26-13LB
AM31-13AC

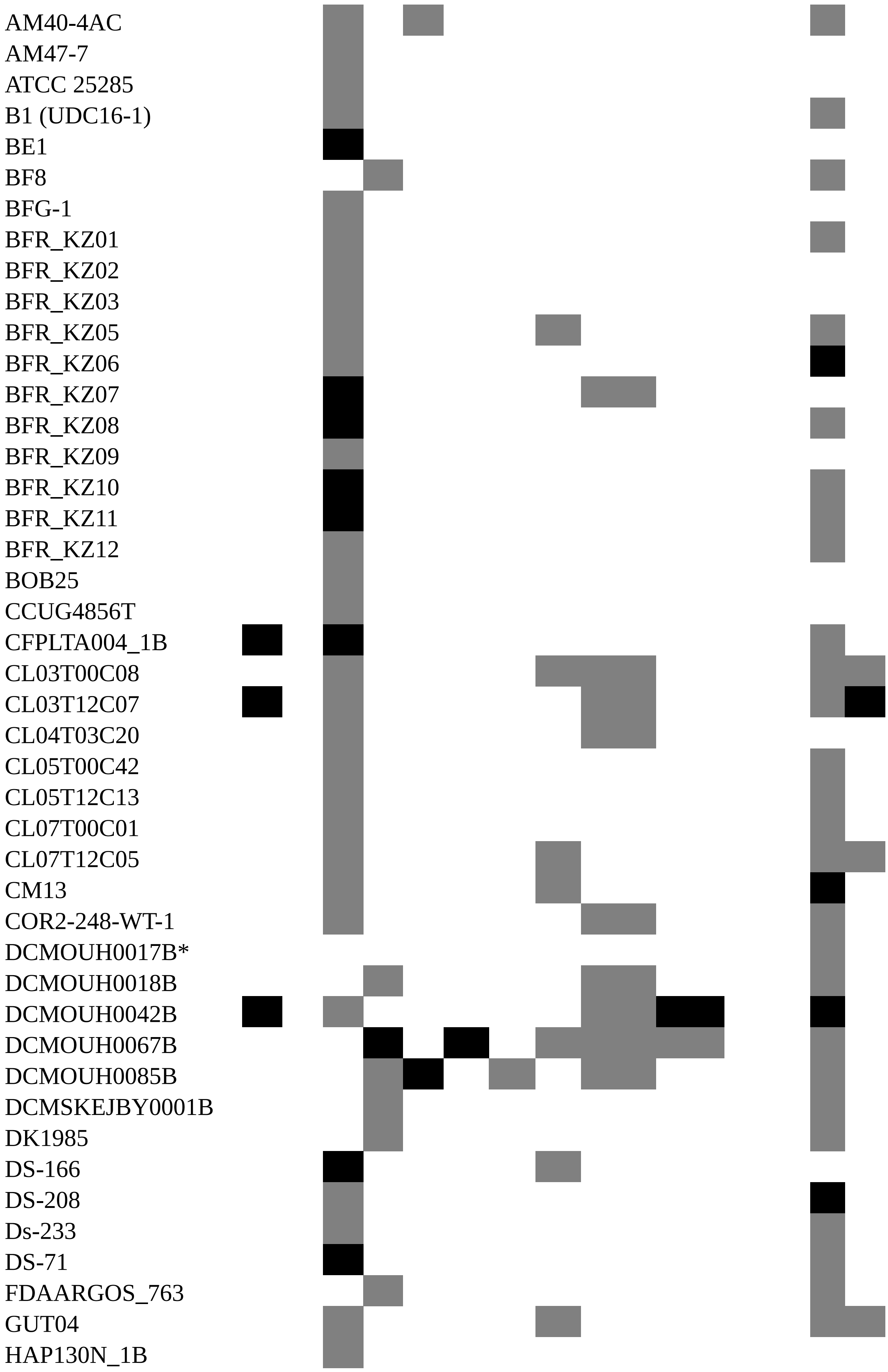
AM40-4AC
AM47-7
ATCC 25285
B1 (UDC16-1)
BE1
BF8
BFG-1
BFR_KZ01
BFR_KZ02
BFR_KZ03
BFR_KZ05
BFR_KZ06
BFR_KZ07
BFR_KZ08
BFR_KZ09
BFR_KZ10
BFR_KZ11
BFR_KZ12
BOB25
CCUG4856T
CFPLTA004_1B
CL03T00C08
CL03T12C07
CL04T03C20
CL05T00C42
CL05T12C13
CL07T00C01
CL07T12C05
CM13
COR2-248-WT-1
DCMOUH0017B*
DCMOUH0018B
DCMOUH0042B
DCMOUH0067B
DCMOUH0085B
DCMSKEJBY0001B
DK1985
DS-166
DS-208
Ds-233
DS-71
FDAARGOS_763
GUT04
HAP130N_1B

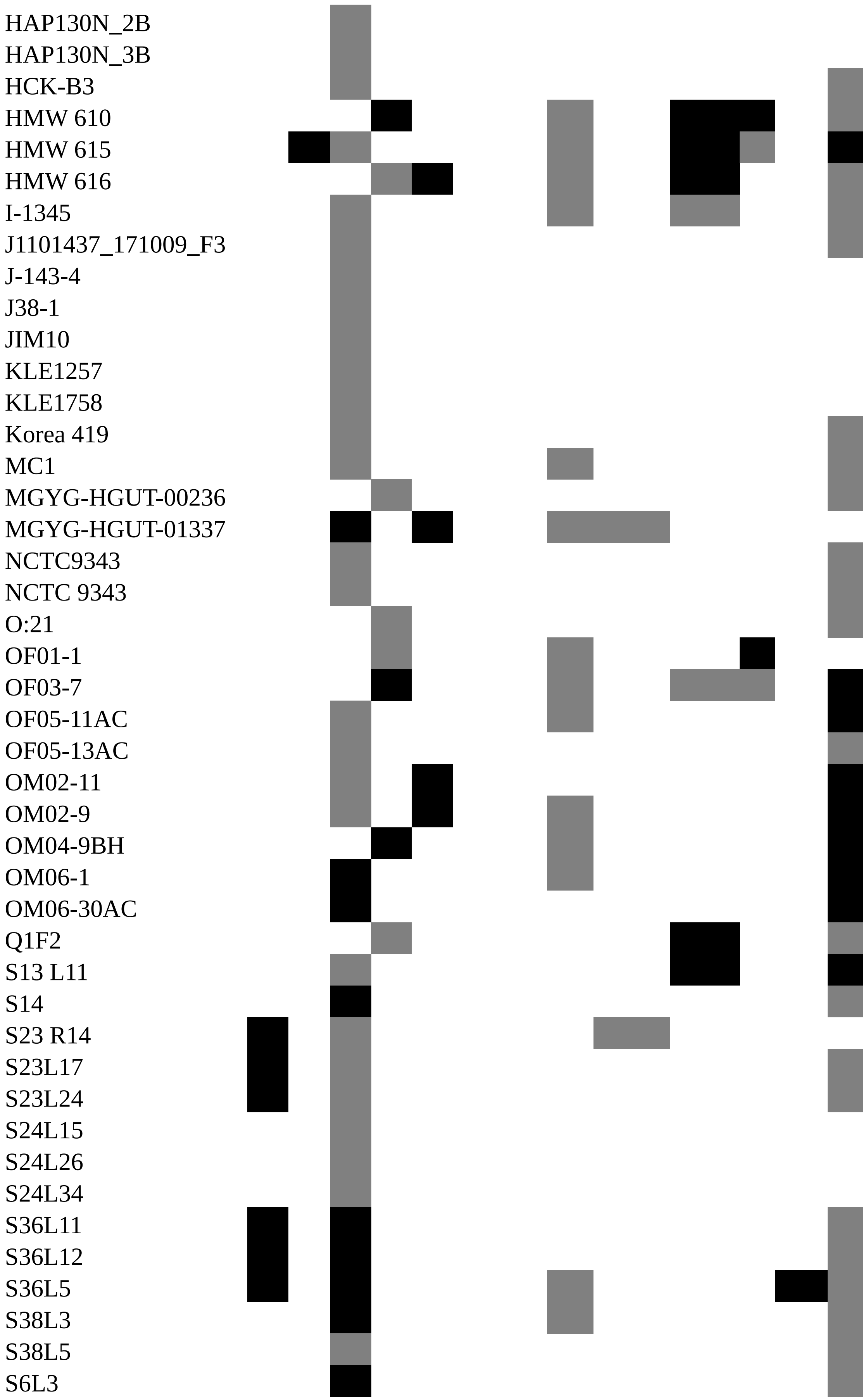
HAP130N_2B
HAP130N_3B
HCK-B3
HMW 610
HMW 615
HMW 616
I-1345
J1101437_171009_F3
J-143-4
J38-1
JIM10
KLE1257
KLE1758
Korea 419
MC1
MGYG-HGUT-00236
MGYG-HGUT-01337
NCTC9343
NCTC 9343
O:21
OF01-1
OF03-7
OF05-11AC
OF05-13AC
OM02-11
OM02-9
OM04-9BH
OM06-1
OM06-30AC
Q1F2
S13 L11
S14
S23 R14
S23L17
S23L24
S24L15
S24L26
S24L34
S36L11
S36L12
S36L5
S38L3
S38L5
S6L3

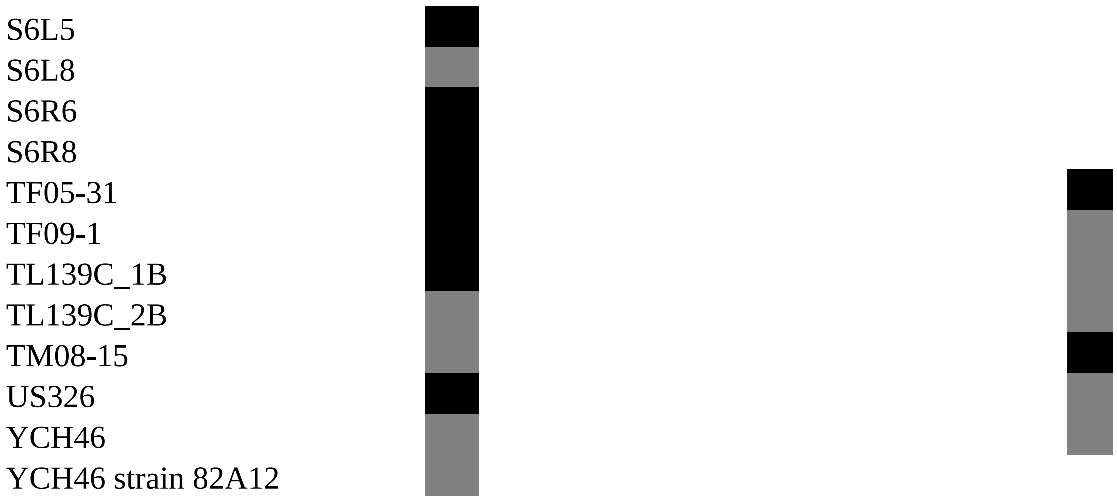
S6L5
S6L8
S6R6
S6R8
TF05-31
TF09-1
TL139C_1B
TL139C_2B
TM08-15
US326
YCH46
YCH46 strain 82A12
