## Supplementary material for "Comprehensive Comparative Genomic revels: *Bacteroides fragilis* is a reservoir of antibiotic resistance genes in the gut microbiota": S4

**Table S4. Results obtained by PhiSpy compared to the VFDB database.** pp and NS correspond to prophage, and non-significant scores, respectively.

| Strain | Start | Stop | Fuction annotation | VFDB-BLAST [blastp/protein sequences from VFDB core dataset (SetB)] | Scores (VFDB) ID % | Bit-Score | E-value |
|---|---|---|---|---|---|---|---|
| **638R** | | | | | | | |
| **pp1** | 213467 | 219708 | | | **NS** | **NS** | **NS** |
| **pp2** | 1364144 | 1432045 | SDR family oxidoreductase | VFG005766 (cylG) [Beta-hemolysin/cytolysin] [*Streptococcus agalactiae* NEM316] | 30 | 110 | 8.00E-25 |
| **pp3** | 2143106 | 2146601 | | | **NS** | **NS** | **NS** |
| **pp4** | 2331998 | 2398444 | ATP-dependent Clp protease | VFG049191 (A79E_1186) ClpB protein [*Klebsiella pneumoniae subsp. pneumoniae* 1084] | 34 | 471 | e-133 |
| **pp5** | 4290022 | 4324198 | Pyridoxal phosphate-dependent aminotransferase | VFG026073 (wcbT) 8-amino-7-oxononanoate synthase [*Burkholderia thailandensis* E264] | 32 | 183 | 2.00E-46 |
| **pp6** | 4758155 | 4779201 | Elongation factor Tu | VFG046475 (F7308_0636) translation elongation factor Tu [*Francisella sp.* TX077308] | 64 | 540 | e-154 |
| **pp7** | 4904224 | 4939422 | | | **NS** | **NS** | **NS** |
| **pp8** | 5191175 | 5238039 | | | **NS** | **NS** | **NS** |
| **BE1** | | | | | | | |
| **pp1** | 479627 | 500672 | ABC transporter ATP-binding protein | VFG030679 (sugC) ABC transporter ATP-binding protein [*Mycobacterium smegmatis* str. MC2 155] | 33 | 127 | 5.00E-30 |
| **pp2** | 975350 | 1008304 | | | **NS** | **NS** | **NS** |
| **pp3** | 2841029 | 2855674 | | | **NS** | **NS** | **NS** |
| **pp4** | 2975840 | 3016078 | Excinuclease ABC subunit UvrA | VFG013248 (msbA) lipid transporter ATP-binding/permease [LOS] [*Haemophilus influenzae* Rd KW20] | 34 | 59 | 0.0000000 2 |
| | | | ATP-binding cassette domain-containing protein | VFG008825 (ddrA) daunorubicin-DIM-transport [*M. liflandii* 128FXT] | 36 | 159 | 5.00E-39 |
| | | | Carbamoyl-phosphate synthase | VFG047708 (carB) carbamoyl phosphate synthase large subunit [*F. Tularensis subsp. tularensis* SCHU S4] | 37 | 715 | 0 |
| **BFG-1** | | | | | | | |
| **pp1** | 2150138 | 2165099 | ATP-dependent Clp protease | VFG000079 ATP-binding chain C [ClpC] [Listeria monocytogenes EGD-e] | 30 | 373 | e-103 |
| **pp2** | 2173185 | 2237377 | Bacteriodales T6SS TssH (ClpV) protein | VFG049191 (A79E_1186) ClpB protein [T6SS-II] [*Klebsiella pneumoniae subsp. pneumoniae* 1084] | 34 | 461 | e-129 |
| **pp3** | 2602065 | 2621841 | | | **NS** | **NS** | **NS** |
| **pp4** | 3834160 | 3848976 | | | **NS** | **NS** | **NS** |

| | | | | | | | |
|---|---|---|---|---|---|---|---|
| **pp5** | 4600404 | 4621446 | Elongation factor Tu | VFG046475 (F7308_0636) translation elongation factor Tu [EF-Tu] [Francisella sp. TX077308] | 64 | 540 | e-154 |
| **BOB25** | | | | | | | |
| **pp1** | 580208 | 601253 | AraC family transcriptional regulator | VFG016125 (pchR) transcriptional regulator PchR [Pyochelin] [*Pseudomonas aeruginosa* PA7] | 37 | 92 | 4.00E-19 |
| **pp2** | 3123325 | 3150969 | Amino acid adenylation domain-containing | VFG050041 (dhbF) DhbF [Bacillibactin] [*Bacillus licheniformis* ATCC 14580] | 34 | 261 | 1.00E-69 |
| **CCUG4856T** | | | | | | | |
| **pp1** | 261413 | 275158 | Sigma-54 dependent transcriptional regulator | VFG013992 (pilR) two component, sigma54 specific, transcriptional regulator [*P. mendocina ymp*] | 37 | 276 | 2.00E-74 |
| **pp2** | 438472 | 459516 | Elongation factor Tu | VFG046475 (F7308_0636) translation elongation factor Tu [*Francisella sp.* TX077308] | 64 | 540 | e-154 |
| **pp3** | 570417 | 582348 | | | **NS** | **NS** | **NS** |
| **pp4** | 2978493 | 3016553 | | | **NS** | **NS** | **NS** |
| **pp5** | 3285120 | 3315959 | | | **NS** | **NS** | **NS** |
| **pp6** | 3627370 | 3660950 | | | **NS** | **NS** | **NS** |
| **pp7** | 3733521 | 3743116 | | | **NS** | **NS** | **NS** |
| **pp8** | 4799815 | 4810122 | | | **NS** | **NS** | **NS** |
| **CL03T12C07** | | | | | | | |
| **pp1** | 1155800 | 1175320 | Bacteriodales T6SS TssH (ClpV) protein | VFG049191 (A79E_1186) ClpB protein [T6SS-II] [*Klebsiella pneumoniae subsp. pneumoniae* 1084] | 34 | 460 | e-129 |
| | | | 8-amino-7-oxononanoate synthase | VFG026073 (wcbT) 8-amino-7-oxononanoate synthase [*B.thailandensis* E264] | 32 | 183 | 2.00E-46 |
| **pp2** | 2203442 | 2238771 | | | **NS** | **NS** | **NS** |
| **pp3** | 2425181 | 2471640 | | | **NS** | **NS** | **NS** |
| **pp4** | 2775806 | 2792171 | peroxiredoxin | VFG009680 (ahpC) alkylhydroperoxide reductase [*M. smegmatis str.* MC2 155] | 35 | 123 | 7.00E-29 |
| **pp5** | 3649049 | 3670094 | | | **NS** | **NS** | **NS** |
| **pp6** | 3717918 | 3740432 | | | **NS** | **NS** | **NS** |
| **DCMOUH0017B** | | | | | | | |
| **pp1** | 401069 | 419313 | | | **NS** | **NS** | **NS** |
| **pp2** | 479699 | 500762 | Elongation factor Tu | VFG046475 (F7308_0636) translation elongation factor Tu [*Francisella sp.* TX077308] | 64 | 540 | e-154 |
| **pp3** | 781148 | 785485 | | | | | |
| **pp4** | 2528549 | 2552148 | Pyridoxal phosphate-dependent aminotransferase | VFG026073 (wcbT) 8-amino-7-oxononanoate synthase [*B. thailandensis* E264] | 32 | 193 | 2.00E-49 |
| | | | ATP-dependent Clp protease | VFG049191 (A79E_1186) ClpB protein [T6SS-II] [*K. pneumoniae subsp.* | 34 | 456 | e-128 |

| | | | | | | | |
|---|---|---|---|---|---|---|---|
| | | | | *pneumoniae* 1084] | | | |
| **pp5** | 2636624 | 2656957 | | | **NS** | **NS** | **NS** |
| **pp6** | 3412629 | 3415909 | | | **NS** | **NS** | **NS** |
| **pp7** | 3704641 | 3729617 | | | **NS** | **NS** | **NS** |
| **pp8** | 3854643 | 3900707 | | | **NS** | **NS** | **NS** |
| **pp9** | 4469287 | 4484136 | | | **NS** | **NS** | **NS** |
| **DCMOUH0018B** | | | | | | | |
| **pp1** | 676769 | 719997 | | | **NS** | **NS** | **NS** |
| **pp2** | 2366649 | 2422672 | | | **NS** | **NS** | **NS** |
| **pp3** | 3604199 | 3624453 | | | **NS** | **NS** | **NS** |
| **pp4** | 3951247 | 3963855 | Histidine kinase | VFG014983 (algZ) alginate biosynthesis protein AlgZ/FimS [*P. stutzeri* A1501] | 30 | 82 | 4.00E-16 |
| **pp5** | 4375125 | 4378366 | | | **NS** | **NS** | **NS** |
| **DCMOUH0042B** | | | | | | | |
| **pp1** | 401915 | 422957 | Elongation factor Tu | VFG046475 (F7308_0636) translation elongation factor Tu [*Francisella sp.* TX077308] | 64 | 540 | e-154 |
| **pp2** | 473969 | 485466 | | | **NS** | **NS** | **NS** |
| **pp3** | 1445469 | 1453182 | | | **NS** | **NS** | **NS** |
| **pp4** | 2215945 | 2241758 | HipA domain-containing protein | VFG045567 (lpg2370) Dot/Icm type IV secretion system effector [*Legionella pneumophila subsp. Philadelphia* 1] | 40 | 175 | 4.00E-45 |
| **pp5** | 2783821 | 2790656 | | | **NS** | **NS** | **NS** |
| **pp6** | 3095387 | 3130079 | ATP-dependent Clp protease | VFG049191 (A79E_1186) ClpB protein [T6SS-II] [*K. pneumoniae subsp. pneumoniae* 1084] | 34 | 460 | e-129 |
| | | | pyridoxal phosphate-dependent aminotransferase | VFG026073 (wcbT) 8-amino-7-oxononanoate synthase [*B.thailandensis* E264] | 32 | 184 | 7.00E-47 |
| **pp7** | 3144978 | 3164803 | | | **NS** | **NS** | **NS** |
| **pp8** | 3184963 | 3195847 | | | **NS** | **NS** | **NS** |
| **pp9** | 3633636 | 3669156 | | | **NS** | **NS** | **NS** |
| **DCMOUH0067B** | | | | | | | |
| **pp1** | 397075 | 418137 | Elongation factor Tu | VFG046475 (F7308_0636) translation elongation factor Tu [*Francisella sp.* TX077308] | 64 | 540 | e-154 |
| **pp2** | 2172900 | 2193349 | Iron ABC transporter permease | VFG044280 (PMI0229) ABC transporter permease [Proteobactin] [*Proteus mirabilis* HI4320] | 31 | 141 | 7.00E-34 |
| **pp3** | 2291800 | 2353062 | Sigma-54 dependent transcriptional regulator | VFG019760 (pilR) putative two-component system, response regulator [*P. fluorescens* SBW25] | 34 | 256 | 2.00E-68 |
| **pp4** | 2888874 | 2892114 | | | **NS** | **NS** | **NS** |

| | | | | | | | |
|---|---|---|---|---|---|---|---|
| **pp5** | 2995991 | 3022368 | | | **NS** | **NS** | **NS** |
| **pp6** | 3056030 | 3067067 | | | **NS** | **NS** | **NS** |
| **pp7** | 3105191 | 3137078 | | | **NS** | **NS** | **NS** |
| **pp8** | 3262258 | 3295323 | | | **NS** | **NS** | **NS** |
| **pp9** | 3557730 | 3561317 | | | **NS** | **NS** | **NS** |
| **pp10** | 4306791 | 4345962 | | | **NS** | **NS** | **NS** |
| **pp11** | 4527352 | 4555968 | | | **NS** | **NS** | **NS** |
| **pp12** | 4572239 | 4588407 | | | **NS** | **NS** | **NS** |
| **pp13** | 4995194 | 5011161 | | | **NS** | **NS** | **NS** |
| **pp14** | 5057869 | 5079922 | | | **NS** | **NS** | **NS** |
| **pp15** | 5403122 | 5417839 | | | **NS** | **NS** | **NS** |
| **DCMOUH0085B** | | | | | | | |
| **pp1** | 406695 | 410214 | | | **NS** | **NS** | **NS** |
| **pp2** | 451520 | 472584 | Elongation factor Tu | VFG046475 (F7308_0636) translation elongation factor Tu [*Francisella sp.* TX077308] | 64 | 540 | e-154 |
| **pp3** | 1572546 | 1599976 | AraC family transcriptional regulator | VFG016125 (pchR) transcriptional regulator PchR [Pyochelin] [*P. aeruginosa* PA7] | 36 | 88 | 9.00E-18 |
| **pp4** | 2142035 | 2145359 | | | | | |
| **pp5** | 2453075 | 2483612 | Sugar O-acetyltransferase | VFG001306 capsular polysaccharide synthesis enzyme Cap8J [*Staphylococcus aureus subsp. Aureus* MW2] | 52 | 86 | 2.00E-18 |
| | | | SDR family oxidoreductase | VFG005766 (cylG) [Beta-hemolysin/cytolysin] [*Streptococcus agalactiae* NEM316] | 34 | 133 | 1.00E-31 |
| **pp6** | 2602881 | 2631872 | Pyridoxal phosphate-dependent aminotransferase | VFG026073 (wcbT) 8-amino-7-oxononanoate synthase [*B. thailandensis* E264] | 32 | 193 | 2.00E-49 |
| | | | ATP-dependent Clp protease | VFG049191 (A79E_1186) ClpB protein [T6SS-II] [*K. pneumoniae subsp. pneumoniae* 1084] | 34 | 471 | e-133 |
| **pp7** | 2977359 | 2990520 | | | **NS** | **NS** | **NS** |
| **pp8** | 3030082 | 3059202 | | | **NS** | **NS** | **NS** |
| **pp9** | 4632999 | 4659737 | | | **NS** | **NS** | **NS** |
| **pp10** | 4951741 | 4955812 | | | **NS** | **NS** | **NS** |
| **pp11** | 5027839 | 5053959 | | | **NS** | **NS** | **NS** |
| **pp12** | 5444772 | 5483500 | | | **NS** | **NS** | **NS** |
| **DCMSKEJBY0001B** | | | | | | | |
| **pp1** | 219812 | 269103 | | | **NS** | **NS** | **NS** |
| **pp2** | 471777 | 475296 | | | **NS** | **NS** | **NS** |

| | | | | | | | |
|---|---|---|---|---|---|---|---|
| **pp3** | 516614 | 537676 | Elongation factor Tu | VFG046475 (F7308_0636) translation elongation factor Tu [*Francisella sp.* TX077308] | 64 | 540 | e-154 |
| **pp4** | 759437 | 766427 | AMP-binding protein | VFG030400 (fadD13) fatty-acid--CoA ligase [*Mycobacterium sp.* JDM601] | 37 | 235 | 5.00E-62 |
| **pp5** | 1537663 | 1540904 | | | **NS** | **NS** | **NS** |
| **pp6** | 1782984 | 1789227 | | | **NS** | **NS** | **NS** |
| **pp7** | 2523982 | 2536423 | | | **NS** | **NS** | **NS** |
| **pp8** | 3205133 | 3224008 | Sugar O-acetyltransferase | VFG005951 (eps9) exopolysaccharide biosynthesis protein [*S. thermophilus* LMG 18311] | 40 | 90 | 1.00E-18 |
| **pp9** | 3503717 | 3521970 | | | **NS** | **NS** | **NS** |
| **pp10** | 3671979 | 3692511 | | | **NS** | **NS** | **NS** |
| **pp11** | 3710587 | 3714109 | | | **NS** | **NS** | **NS** |
| **pp12** | 4891730 | 4902194 | | | **NS** | **NS** | **NS** |
| **DK1985** | | | | | | | |
| **pp1** | 416195 | 437257 | Elongation factor Tu | VFG046475 (F7308_0636) translation elongation factor Tu [*Francisella sp.* TX077308] | 64 | 540 | e-154 |
| **pp2** | 2163127 | 2202954 | Iron ABC transporter permease | VFG044280 (PMI0229) ABC transporter permease [Proteobactin] [*Proteus mirabilis* HI4320] | 30 | 136 | 2.00E-32 |
| **pp3** | 2370473 | 2386555 | | | **NS** | **NS** | **NS** |
| **pp4** | 2405645 | 2426162 | | | **NS** | **NS** | **NS** |
| **pp5** | 2471952 | 2489502 | | | **NS** | **NS** | **NS** |
| **pp6** | 3077073 | 3150518 | LuxR C-terminal-related transcriptional | VFG042095 (ECs3712) hypothetical protein [ETT2] [*Escherichia coli* O157:H7 str. Sakai] | 51 | 56 | 2.00E-08 |
| | | | DUF1911 domain-F7887_RS12570 | VFG038374 (AHA_1829) Type VI secretion system protein-T6SS [*Aeromonas hydrophila subsp. hydrophila* ATCC 7966] | 30 | 70 | 2E-12 |
| **pp7** | 3567449 | 3593784 | | | **NS** | **NS** | **NS** |
| **pp8** | 4248688 | 4265823 | | | **NS** | **NS** | **NS** |
| **pp9** | 4349082 | 4357085 | | | **NS** | **NS** | **NS** |
| **pp10** | 4608624 | 4641060 | | | **NS** | **NS** | **NS** |
| **pp11** | 4939816 | 4950333 | | | **NS** | **NS** | **NS** |
| **FDAARGOS_763** | | | | | **NS** | **NS** | **NS** |
| **pp1** | 913365 | 941945 | | | **NS** | **NS** | **NS** |
| **pp2** | 1052013 | 1083456 | | | **NS** | **NS** | **NS** |
| **pp3** | 1140292 | 1159655 | | | **NS** | **NS** | **NS** |
| **pp4** | 1168994 | 1176673 | | | **NS** | **NS** | **NS** |
| **pp5** | 1193865 | 1224243 | | | **NS** | **NS** | **NS** |
| **pp6** | 1348969 | 1355481 | | | **NS** | **NS** | **NS** |

| | | | | | | | |
|---|---|---|---|---|---|---|---|
| **pp7** | 2757828 | 2768324 | | | **NS** | **NS** | **NS** |
| **pp8** | 2861582 | 2888685 | | | **NS** | **NS** | **NS** |
| **pp9** | 2968643 | 2976855 | | | **NS** | **NS** | **NS** |
| **pp10** | 3318234 | 3330492 | | | **NS** | **NS** | **NS** |
| **pp11** | 3904632 | 3925746 | | | **NS** | **NS** | **NS** |
| **pp12** | 3966844 | 3987906 | Elongation factor Tu | VFG046475 (F7308_0636) translation elongation factor Tu [*Francisella sp.* TX077308] | 64 | 540 | e-154 |
| **pp13** | 4254840 | 4259532 | | | | | |
| **pp14** | 4770855 | 4835904 | PAS domain-containing hybrid sensor histidine | VFG011124 (bvgS) virulence sensor protein [Two-component system] [*Bordetella avium* 197N] | 31 | 1.59E+02 | 8.00E-39 |
| | | | Efflux RND transporter permease subunit | VFG037724 (adeG) Cation/multidrug efflux pump [*Acinetobacter baumannii* BJAB0715] | 31 | 566 | e-161 |
| | | | SDR family oxidoreductase | VFG038840 (flmH) flagellar-related 3-oxoacyl-ACP reductase [Polar flagella] [*A. hydrophila* ML09-119] | 34 | 124 | 6.00E-29 |
| **pp15** | 5214229 | 5244231 | | | **NS** | **NS** | **NS** |
| **GUT04** | | | | | | | |
| **pp1** | 1487092 | 1505540 | | | **NS** | **NS** | **NS** |
| **pp2** | 1487092 | 1505540 | ATP-dependent Clp protease | VFG049191 (A79E_1186) ClpB protein [T6SS-II] [*K. pneumoniae subsp. pneumoniae* 1084] | 34 | 460 | e-129 |
| | | | Pyridoxal phosphate-dependent aminotransferase | VFG026073 (wcbT) 8-amino-7-oxononanoate synthase [*B. thailandensis* E264] | 32 | 185 | 6.00E-47 |
| **pp3** | 2478257 | 2486588 | | | **NS** | **NS** | **NS** |
| **pp4** | 3056373 | 3084295 | Sugar O-acetyltransferase | VFG005951 (eps9) exopolysaccharide biosynthesis protein [*S. thermophilus* LMG 18311] | 38 | 89 | 2.00E-18 |
| **pp5** | 3462101 | 3495449 | | | **NS** | **NS** | **NS** |
| **pp6** | 3948266 | 3969309 | Elongation factor Tu | VFG046475 (F7308_0636) translation elongation factor Tu [*Francisella sp.* TX077308] | 64 | 540 | e-154 |
| **pp7** | 4034791 | 4056128 | | | **NS** | **NS** | **NS** |
| **pp8** | 4247250 | 4256540 | | | **NS** | **NS** | **NS** |
| **pp9** | 4342071 | 4346408 | | | **NS** | **NS** | **NS** |
| **pp10** | 4612082 | 4658511 | | | **NS** | **NS** | **NS** |
| **NCTC 9343** | | | | | | | |
| **pp1** | 2019195 | 2053561 | Tyrosine-type recombinase/integrase | VFG048223 (KOX_22330) integrase family protein [*K. oxytoca* KCTC 1686*]* | 31 | 85 | 9.00E-17 |
| **pp2** | 2348911 | 2381628 | ATP-dependent Clp protease | VFG049191 (A79E_1186) ClpB protein [T6SS-II] [*K. pneumoniae subsp. pneumoniae* 1084] | 34 | 460 | e-129 |
| | | | pyridoxal phosphate- | VFG026073 (wcbT) 8-amino-7-oxononanoate synthase [*B. thailandensis* E264] | 32 | 182 | 4.00E-46 |

| | | | | | | | |
|---|---|---|---|---|---|---|---|
| | | | dependent aminotransferase | | | | |
| **pp3** | 3863607 | 3873914 | | | | | |
| **pp4** | 4707403 | 4728447 | Elongation factor Tu | VFG046475 (F7308_0636) translation elongation factor Tu [*Francisella sp.* TX077308] | 64 | 540 | e-154 |
| **Q1F2** | | | | | | | |
| **pp1** | 1737 | 12253 | | | **NS** | **NS** | **NS** |
| **pp2** | 1733831 | 1755209 | | | **NS** | **NS** | **NS** |
| **pp3** | 2743845 | 2747091 | | | **NS** | **NS** | **NS** |
| **pp4** | 2954006 | 2957247 | | | **NS** | **NS** | **NS** |
| **pp5** | 3224591 | 3244202 | | | **NS** | **NS** | **NS** |
| **pp6** | 4049748 | 4070809 | Elongation factor Tu | VFG046475 (F7308_0636) translation elongation factor Tu [*Francisella sp.* TX077308]] | 64 | 540 | e-154 |
| **pp7** | 5007463 | 5032170 | | | **NS** | **NS** | **NS** |
| **pp8** | 5150670 | 5161165 | | | **NS** | **NS** | **NS** |
| **S14** | | | | | | | |
| **pp1** | 1337957 | 1368532 | | | **NS** | **NS** | **NS** |
| **pp2** | 1852018 | 1875265 | | | **NS** | **NS** | **NS** |
| **pp3** | 2998563 | 3008870 | | | **NS** | **NS** | **NS** |
| **pp4** | 3804777 | 3825821 | Elongation factor Tu | VFG046475 (F7308_0636) translation elongation factor Tu [*Francisella sp.* TX077308] | 64 | 540 | e-154 |
| **pp5** | 3884946 | 3896876 | | | **NS** | **NS** | **NS** |
| **YCH46** | | | | | | | |
| **pp1** | 160252 | 163493 | | | **NS** | **NS** | **NS** |
| **pp2** | 763378 | 769947 | Sugar O-acetyltransferase | VFG005951 (eps9) exopolysaccharide biosynthesis protein [*S. thermophilus* LMG 18311] | 33 | 91 | 6.00E-19 |
| **pp3** | 2279158 | 2329554 | ATP-dependent Clp protease | VFG049191 (A79E_1186) ClpB protein [T6SS-II] [*K. pneumoniae subsp. pneumoniae* 1084] | 34 | 461 | e-129 |
| | | | Pyridoxal phosphate-dependent aminotransferase | VFG026073 (wcbT) 8-amino-7-oxononanoate synthase [*B. thailandensis* E264] | 32 | 183 | 2.00E-46 |
| **pp4** | 2685951 | 2705412 | | | **NS** | **NS** | **NS** |
| **pp5** | 2823776 | 2858268 | | | **NS** | **NS** | **NS** |
| **pp6** | 3201025 | 3217152 | | | **NS** | **NS** | **NS** |
| **pp7** | 3408488 | 3418012 | | | **NS** | **NS** | **NS** |
| **pp8** | 3956548 | 3973535 | | | **NS** | **NS** | **NS** |
| **pp9** | 4775764 | 4796809 | | | **NS** | **NS** | **NS** |
| **pp10** | 4905881 | 4917802 | | | **NS** | **NS** | **NS** |
