## Supplementary material for "Comprehensive Comparative Genomic revels: *Bacteroides fragilis* is a reservoir of antibiotic resistance genes in the gut microbiota": S5

**Table S5. Results obtained by ICEFinder compared to NCBI BLAST and CARD databases.**

| | | | | NCBI BLAST Database | | | | | | Antibiotic Resistance Gene Results | | | |
|---|---|---|---|---|---|---|---|---|---|---|---|---|---|
| **Strains** | **Region** | **Location in Query** | **Type** | **BLAST Description** | **Location in Subject** | **Query Covarage %** | **Bit-Score** | **E-value** | **ID %** | **CARD Description** | **Bit-Score** | **E-value** | **ID %** |
| **638R** | | | | | | | | | | | | | |
| | 1 | 94878-207753 | Putative ICE with T4SS | GUT04 | 4695315 - 4785706 | 100 | 1.649e+05 | 0.0 | 99.6 | | | | |
| | 2 | 2901862-2977865 | Putative ICE with T4SS | DCMOUH0042B | 3771074-3809796 | 96 | 70112 | 0.0 | 99.3 | | | | |
| | 3 | 3723277-3768753 | Putative ICE with T4SS | CCUG4856T | 4613995-4646762 | 98 | 59489 | 0.0 | 99.4 | | | | |
| | 4 | 3942134-3966444 | Putative IME | BOB25 | 3848452-3872761 | 100 | 44859 | 0.0 | 99.9 | | | | |
| **BE1** | | | | | | | | | | | | | |
| | 1 | 1336630-1381569 | Putative IME | S14 | 643455-668985 | 58 | 46416 | 0.0 | 99.5 | | | | |
| | 2 | 2814811-2835620 | Putative IME | BOB25 | 2658485-2679294 | 100 | 38086 | 0.0 | 99.7 | | | | |
| | 3 | 3708073-3749605 | Putative IME | YCH46 | 3875108- 3916650 | 100 | 75582 | 0.0 | 99.5 | | | | |
| **BFG-1** | | | | | | | | | | | | | |
| | 1 | 1383979-1390739 | Putative IME without identified DR | B. thetaiotaomicron strain FDAARGOS_935 | 4389677-4396437 | 100 | 12486 | 0.0 | 100 | | | | |
| | 2 | 3793574-3854553 | Putative IME | BOB25 | 3831755-3872157 | 92 | 73532 | 0.0 | 99 | | | | |
| **BOB25** | | | | | | | | | | | | | |
| | 1 | 442212- | Putative | CL03T12C07 | 4610001- 4626069 | 100 | 29362 | 0.0 | 100 | | | | |

| | | | | | | | | | |
|---|---|---|---|---|---|---|---|---|---|
| | | 458279 | IME | | | | | | |
| | 2 | 710043-723195 | Putative IME without identified DR | GUT04 | 8766-21919 | 100 | 23946 | 0.0 | 99.5 |
| | 3 | 1464254-1476689 | Putative IME | 638R | 1587946-1600381 | 100 | 22916 | 0.0 | 99.9 |
| | 4 | 1736733-1769949 | Putative ICE with T4SS | | 1906125-1939343 | 100 | 60761 | 0.0 | 99.7 |
| | 5 | 3107685-3116369 | Putative IME without identified DR | BE1 | 3186923-3195609 | 100 | 15825 | 0.0 | 99.5 |
| | 6 | 3864050-3870829 | Putative IME without identified DR | 638R | 3957732-3964512 | 100 | 12504 | 0.0 | 99.9 |
| | 7 | 4641279-4682413 | Putative ICE with T4SS | CL03T12C07 | 3611961-3650482 | 99 | 70315 | 0.0 | 99.6 |
| **CL03T12C07** | | | | | | | | | |
| | 1 | 234584-281526 | Putative ICE with T4SS | *Parabacteroides distasonis* CL03T12C09 | 669863-681657 | 48 | 21782 | 0.0 | 100 |
| | 2 | 338652-344158 | Putative IME without identified DR | *B.uniformis JCM13288* | 1989959-1995462 | 100 | 8427 | 0.0 | 94.3 |

| | | | | | | | | | | | | | |
|---|---|---|---|---|---|---|---|---|---|---|---|---|---|
| | 3 | 415608-458082 | Putative ICE with T4SS | *B. uniformis* CL03T12C37 | 2589808-2632282 | 100 | 78437 | 0.0 | 100 | | | | |
| | 4 | 522065-543592 | Putative IME | DCMOUH0042B | 2441255-2456251 | 97 | 24131 | 0.0 | 95.7 | | | | |
| | 5 | 2186363-2255887 | Putative ICE with T4SS | *P.distasonis* CL03T12C09 | 2809432-2865775 | 81 | 1.040e+05 | 1.077 e+05 | 99.9 | | | | |
| | 6 | 2685338-2783348 | Putative ICE with T4SS | *B. vulgatus* ATCC 8482 | 2041154-2076327 | 60 | 60525 | 1.548 e+05 | 97.7 | | | | |
| | 7 | 3145989-3219056 | Putative ICE with T4SS | *P. merdae* BFG-280 | 1898788-1931046 | 64 | 59516 | 0.0 | 99.9 | | | | |
| | | 3161472-3163397 | | | | | | | | ARO:3000191 [*TETQ*] [*Bacteroides fragilis*] | 1287 | 0.0 | 97 |
| | | 3177393-3178193 | | | | | | | | ARO:3000498 [*ERMF*] [[*Bacteroides fragilis*] pseudogene | 1419 | 0.0 | 98 |
| | | 3178499-3179638 | | | | | | | | ARO:3005166 [*TETX1)*] [*Bacteroides thetaiotaomicron*] | 737 | 0.0 | 99 |
| | | 3179807-3180973 | | | | | | | | ARO:3000205 [*TETX*] [*Bacteroides fragilis*] | 798 | 0.0 | 99 |
| | | 3181149-3182012 | | | | | | | | ARO:3004683 [*AADS*] Transposon (Tn4551) | 583 | 0.0 | 100 |
| | 8 | 3722159-3742410 | Putative IME | FDAARGOS_1225 | | 64 | 21228 | 0.0 | 97.8 | | | | |
| **DCMOUH0017B** | | | | | | | | | | | | | |
| | 1 | 559182-572069 | Putative IME | DCMOUH0067B | 476550-489459 | 100 | 21859 | 0.0 | 97.3 | | | | |
| | 2 | 718104-822380 | Putative ICE with T4SS | DK1985 | 640536-703245 | 62 | 1.147e+05 | 0.0 | 99.7 | | | | |
| | | 801744-803690 | | | | | | | | ARO:3000191 [*TETQ*] [*Bacteroides fragilis*] | 1301 | 0.0 | 98 |

| | | | | | | | | | | | | | |
|---|---|---|---|---|---|---|---|---|---|---|---|---|---|
| | 3 | 2408617-2492949 | Putative ICE with T4SS | DK1985 | 2284239-2326198 | 58 | 76219 | 0.0 | 99.5 | | | | |
| | 4 | 3296126-3320708 | Putative ICE without identified DR | *Myoviridae sp.* isolate ctNMP7 (Viruses, Caudovirales) | 14189-18020 | 23 | 1834 | 0.0 | 75.6 | | | | |
| | 5 | 3972537-4011739 | Putative ICE with T4SS | *Bacteroides sp.* ZJ-18 | 3535222-355202 | 52 | 23305 | 0.0 | 91.8 | | | | |
| | 6 | 5024366-5073661 | Putative IME | DK1985 | 4944516-4967919 | 85 | 43148 | 0.0 | 99.9 | | | | |
| **DCMOUH0018B** | | | | | | | | | | | | | |
| | 1 | 459588-466424 | Putative IME without identified DR | *Odoribacter splanchnicus* NCTC10825 (Bacteroidales) | 1186437-1192718 | 91 | 9888 | 0.0 | 95.1 | | | | |
| | 2 | 3497905-3538007 | Putative IME | DK1985 | 3507173-3512864 | 17 | 10384 | 0.0 | 99.6 | | | | |
| | 3 | 3950960-4008026 | Putative ICE with T4SS | *B.thetaiotaomicron* DSM 2079 | 3657201-3673083 | 45 | 39337 | 0.0 | 96.7 | | | | |
| | 4 | 4718681-4728986 | Putative IME | *Phocaeicola vulgatus* VIC01 (Bacteroidales) | 4021919-4030332 | 95 | 14619 | 0.0 | 98.0 | | | | |
| | | 4726906-4727418 | | | | | | | | ARO:3002835 [LNUA] [*Staphylococcus haemolyticus*] | 175 | 1.65E-56 | 52 |
| | | 4727443-4728648 | | | | | | | | ARO:3004659 [*MEF(EN2)*] [*Bacteroides fragilis*] | 792 | 0.0 | 99 |
| **DCMOUH0042B** | | | | | | | | | | | | | |
| | 1 | 654998- | Putative | *Phocaeicola dorei* | 2829186-2832719 | 87 | 4985 | 0.0 | 92.2 | | | | |

| | | | | | | | | | |
|---|---|---|---|---|---|---|---|---|---|
| | | 660771 | IME without identified DR | JR01 (Bacteroidales) | | | | | |
| | 2 | 2703903-2771046 | Putative ICE with T4SS | YCH46 | 1917483-1946664 | 92 | 53404 | 0.0 | 99.7 |
| | 3 | 3124460-3196107 | Putative ICE with T4SS | *Caudovirales sp.* isolate ctodZ3 (Viruses, Caudovirales) | 40164-77124 | 51 | 68116 | 0.0 | 99.9 |
| **DCMOUH0067B** | | | | | | | | | |
| | 1 | 476551-489459 | Putative IME | DCMOUH0017B | 559183-572069 | 100 | 21858 | 0.0 | 97.5 |
| | 2 | 1334428-1359277 | Putative IME | *Butyricimonas faecalis* strain H184 (Bacteroidales) | 41182-49645 | 41 | 14423 | 0.0 | 97.4 |
| | | 1342273-1343559 | | *B.fragilis* H3 metallo-beta-lactamase (cfiA) gene, partial cds and insertion sequence ISBf11 | 161-1445 | 99 | 2302 | 0.0 | 98.9 |
| | 3 | 2177817-2228671 | Putative IME | DK1985 | 2241863-2268818 | 87 | 47611 | 0.0 | 98.5 |
| | 4 | 2293365-2383506 | Putative ICE with T4SS | DCMOUH0017B | 2454710-2480098 | 64 | 44442 | 0.0 | 98.3 |
| | 5 | 2860251-2925828 | Putative ICE with T4SS | *Ornithobacterium rhinotracheale* ORT-UMN 88 (Flavobacteriales) | 2117090-2154373 | 56 | 68696 | 0.0 | 99.9 |

| | | | | | | | | | | | | | | |
|---|---|---|---|---|---|---|---|---|---|---|---|---|---|---|
| | | 2902927-2904852 | | | | | | | | ARO:3000191 [*Bacteroides fragilis*] | [*TETQ*] | 1287 | 0.0 | 97 |
| | 6 | 2985236-3047486 | Putative IME | *B.eggerthii* FDAARGOS_1221 | 3737969-3785581 | 76 | 89765 | 0.0 | 99.9 | | | | | |
| | 7 | 3261885-3312982 | Putative ICE with T4SS | BOB25 | 608614-623860 | 67 | 21182 | 0.0 | 91.7 | | | | | |
| | 8 | 3739795-3746756 | Putative IME | DCMOUH0085B | 3682195-3685832 | 57 | 6362 | 0.0 | 98.2 | | | | | |
| | 9 | 4488310-4547926 | Putative ICE with T4SS | *B. uniformis* CL03T12C37 | 4051458-4094477 | 73 | 78930 | 0.0 | 99.8 | | | | | |
| | 10 | 4994931-5009696 | Putative IME | DCMSKEJBY0001B | 4893965-4900970 | 74 | 11062 | 0.0 | 95.1 | | | | | |
| | 11 | 5403122-5437411 | Putative IME without identified DR | *B.xylanisolvens* CL11T00C03 | 5103641-5122467 | 95 | 31316 | 0.0 | 96.7 | | | | | |
| **DCMOUH0085B** | | | | | | | | | | | | | | |
| | 1 | 1582421-1623592 | Putative ICE with T4SS | *Phocaeicola dorei* JR04 (Bacteroidales) | 2093147-2102402 | 22 | 15073 | 0.0 | 96.1 | | | | | |
| | 2 | 2115666-2145359 | Putative IME without identified DR | *B. xylanisolvens* XB1A | 4256519-4276628 | 87 | 30184 | 0.0 | 93.8 | | | | | |
| | | 2127830-2129755 | | | | | | | | ARO:3000191 [*Bacteroides fragilis*] | [*TETQ*] | 1186 | 0.0 | 89 |
| | 3 | 2597413- | Putative | *B.thetaiotaomicron* | 4389677-4396437 | 100 | 11273 | 0.0 | 96.8 | | | | | |

| | | | | | | | | | | | | | |
|---|---|---|---|---|---|---|---|---|---|---|---|---|---|
| | | 2604173 | IME without identified DR | FDAARGOS_935 | | | | | | | | | |
| | 4 | 2993358-3044318 | Putative ICE with T4SS | *Butyricimonas faecalis* H184 (Bacteroidales) | 1297236-1335207 | 98 | 70088 | 0.0 | 99.9 | | | | |
| | | 3029059-3029925 | | | | | | | | ARO:3002629 [*ANT(6)-IB* [*Campylobacter fetus subsp.* Fetus] | 440 | 5.32E-157 | 72 |
| | | 3032264-3033025 | | | | | | | | ARO:3000375 [*ERMB*] [*Enterococcus faecium*] | 482 | 1.81E-1741 | 98 |
| | | 3036822-3037598 | | | | | | | | ARO:3002630 ANT(9)-Ia [*Staphylococcus aureus*] | 248 | 2.40E-82 | 47 |
| | 5 | 4993673-5067015 | Putative ICE with T4SS | DCMSKEJBY0001B | 4918590-4936412 | 49 | 31076 | 0.0 | 98.1 | | | | |
| | 6 | 5200112-5246628 | Putative IME | DCMSKEJBY0001B | 5118907-5143906 | 55 | 44577 | 0.0 | 98.8 | | | | |
| | | 5210244-5211209 | | | | | | | | ARO:3003002 *CFXA2* [*Prevotella intermedia*] | 661 | 0.0 | 100 |
| | 7 | 5406706-5484510 | Putative ICE with T4SS | *Butyricimonas faecalis* H184 (Bacteroidales) | 1233504-1275145 | 53 | 76585 | 0.0 | 99.9 | | | | |
| | | 5432028-5433953 | | | | | | | | ARO:3000191 [*TETQ*] [*Bacteroides fragilis*] | 128 | 0.0 | 96 |
| **DCMSKEJBY0001B** | | | | | | | | | | | | | |
| | 1 | 216517-289653 | Putative ICE with T4SS | *B.thetaiotaomicron* F9-2 | 4838710-4903521 | 94 | 1.185e+05 | 0.0 | 99.7 | | | | |

| | | | | | | | | | | | | | |
|---|---|---|---|---|---|---|---|---|---|---|---|---|---|
| | 2 | 1509223-1540904 | Putative IME without identified DR | *Bacteroides sp.* PHL 2737 | 1589526-1621207 | 100 | 58506 | 0.0 | 100 | | | | |
| | | 1524919-1526844 | | | | | | | | ARO:3000191 [*TETQ*] [*Bacteroides fragilis*] | 1281 | 0.0 | 96 |
| | 3 | 2351125-2373640 | Putative IME | *Bacteroides sp.* PHL 2737 | 2426646-244916 | 100 | 41580 | 0.0 | 100 | | | | |
| | 4 | 2404377-2423319 | Putative IME | *Phocaeicola vulgatus* FDAARGOS_1098 (Bacteroidales) | 4007266-4026190 | 99 | 33833 | 0.0 | 98.9 | | | | |
| | 5 | 2491582-2551726 | Putative ICE with T4SS | *B.xylanisolvens strain* funn3 | 798744-819804 | 53 | 37589 | 0.0 | 98.9 | | | | |
| | 6 | 2952506-2984524 | Putative IME | *Bacteroides sp.* PHL 2737 | 3053431-3070700 | 72 | 31892 | 0.0 | 100 | | | | |
| | 7 | 3189962-3219977 | Putative ICE with T4SS | *Bacteroides sp.* PHL 2737 | 3276151-3306166 | 100 | 55430 | 0.0 | 100 | | | | |
| | 8 | 3481900-3533006 | Putative IME | *Bacteroides sp.* PHL 2737 | 3566669-3614021 | 100 | 87380 | 0.0 | 99.9 | | | | |
| | 9 | 4865357-4904826 | Putative IME | *Bacteroides sp.* PHL 2737 | 5087499-5100867 | 55 | 24683 | 0.0 | 99.9 | | | | |
| **DK1985** | | | | | | | | | | | | | |
| | 1 | 2145791-2228778 | Putative ICE with T4SS | DCMSKEJBY0001B | 2322774-2352883 | 40 | 54408 | 0.0 | 99.3 | | | | |
| | 2 | 2275790-2289426 | Putative IME | BOB25 | 1409246-1418356 | 87 | 16183 | 0.0 | 98.7 | | | | |

| | | | | | | | | | | | | | | |
|---|---|---|---|---|---|---|---|---|---|---|---|---|---|---|
| | 3 | 2359292-2401194 | Putative IME | *B.thetaiotaomicron* FDAARGOS_935 | 4389499-4402131 | 35 | 22077 | 0.0 | 98.2 | | | | | |
| | 4 | 3067457-3147352 | Putative ICE with T4SS | *Parabacteroides distasonis* FDAARGOS_759 | 1889831-1901585 | 33 | 18842 | 0.0 | 95.6 | | | | | |
| | 5 | 3553239-3652219 | Putative ICE with T4SS | DCMOUH0017B | 4003634-4036773 | 94 | 61152 | 0.0 | 99.9 | | | | | |
| **FDAARGOS_763** | | | | | | | | | | | | | | |
| | 1 | 329883-379844 | Putative IME | DK1985 | 3875836-3902450 | 73 | 48041 | 0.0 | 99.2 | | | | | |
| | 2 | 804610-831187 | Putative ICE with T4SS | *B. uniformis* JCM13288 | 1720461-1735534 | 100 | 27750 | 0.0 | 99.9 | | | | | |
| | 3 | 899212-976022 | Putative IME | DCMOUH0085B | 3465709-3490456 | 62 | 44996 | 0.0 | 99.5 | | | | | |
| | 4 | 1110377-1170387 | Putative ICE with T4SS | CL03T12C07 | 4800925-4820937 | 99 | 36128 | 0.0 | 99.3 | | | | | |
| | 5 | 2761749-2774618 | Putative IME | Q1F2 | 5144376-5157244 | 100 | 23721 | 0.0 | 99.9 | | | | | |
| | 6 | 3468471-3506361 | Putative IME | Q1F2 | 4519271-4546922 | 74 | 48484 | 0.0 | 98.3 | | | | | |
| | 7 | 4227272-4291896 | Putative ICE with T4SS | *Phocaeicola dorei* JR04 (Bacteroidales) | 169071-204650 | 57 | 65704 | 0.0 | 100 | | | | | |
| | | 4270351-4272276 | | | | | | | | ARO:3000191 [*Bacteroides fragilis*] | [*TETQ*] | 1281 | 0.0 | 96 |
| **GUT04** | | | | | | | | | | | | | | |

| | | | | | | | | | | | | | |
|---|---|---|---|---|---|---|---|---|---|---|---|---|---|
| | 1 | 683264-690043 | Putative IME without identified DR | BOB25 | 1411404-1418183 | 100 | 12499 | 0.0 | 99.9 | | | | |
| | 2 | 2484166-2534271 | Putative ICE without identified DR | CCUG4856T | 2685747-2724774 | 99 | 70727 | 0.0 | 99.4 | | | | |
| | 3 | 4322995-4374014 | Putative IME | Q1F2 | 2955695-2978839 | 84 | 37166 | 0.0 | 95.7 | | | | |
| **NCTC 9343** | | | | | | | | | | | | | |
| | 1 | 1993262-2186284 | Putative ICE with T4SS | CCUG4856T | 3029915-3122493 | 100 | 1.707e+05 | 0.0 | 99.9 | | | | |
| **Q1F2** | | | | | | | | | | | | | |
| | 1 | 2743845-2771693 | Putative IME without identified DR | *B. xylanisolvens* XB1A | 4265908-4279136 | 85 | 19643 | 0.0 | 93.5 | | | | |
| | | 2757939-2759864 | | | | | | | | ARO:3000191 [*TETQ*] [*Bacteroides fragilis*] | 282 | 0.0 | 96 |
| | | 2765896-2766696 | | | | | | | | ARO:3000498 [*ERMF*] [*Bacteroides fragilis*] | 353 | 2.59E-125 | 63 |
| | 2 | 2937644-2990379 | Putative IME | *B. xylanisolvens* funn3 | 3098641-3122737 | 76 | 44453 | 0.0 | 99.9 | | | | |
| | | 2968093-2970039 | | | | | | | | ARO:3000191 [*TETQ*] [*Bacteroides fragilis*] | 1283 | 0.0 | 97 |
| | 3 | 5144376-5157244 | Putative | FDAARGOS_763 | 2761749-2774618 | 100 | 23721 | 0.0 | 99.9 | | | | |

| | | | | | | | | | |
|---|---|---|---|---|---|---|---|---|---|
| | | | IME | | | | | | |
| **S14** | | | | | | | | | |
| | 1 | 3846386-3869709 | Putative IME | CCUG4856T | 533594-545480 | 72 | 20943 | 0.0 | 98.5 |
| **YCH46** | | | | | | | | | |
| | 1 | 3649673-3692378 | Putative ICE with T4SS | CL03T12C07 | 2319892-2362627 | 100 | 77362 | 0.0 | 99.3 |
| | 2 | 3754864-3761625 | Putative IME without identified DR | GUT04 | 2856364-2863125 | 100 | 12272 | 0.0 | 99.4 |
| | 3 | 4507328-4515706 | Putative IME without identified DR | BE1 | 4413326-4421704 | 100 | 15368 | 0.0 | 99.8 |
| | 4 | 5108133-5156366 | Putative ICE with T4SS | CCUG4856T | 772106-820339 | 100 | 88169 | 0.0 | 99.7 |
