## Supplementary material for "Comprehensive Comparative Genomic revels: *Bacteroides fragilis* is a reservoir of antibiotic resistance genes in the gut microbiota": S6

### Table S6. Results obtained by MobileElementFinder.

| Strain | Accession | IS Type | Family | Accession no. | Position in Contigs | Coverage % | ID % |
|---|---|---|---|---|---|---|---|
| **143** | SRKB01000004.1 | IS4351 | IS30 | M17124 | 492641-493796 | 99.9 | 99.8 |
| | SRKB01000010.1 | ISBf5 | IS1182 | NC_006347 | 106027-107856 | 100 | 99.9 |
| | SRKB01000020.1 | IS612 | IS1380 | AB042548 | 1-1598 | 99.6 | 99.3 |
| **12905** | PDCW01000080.1 | IS613 | IS1380 | AB042549 | 1-1594 | 99.7 | 93.2 |
| **1001175st1_C3** | SPGY01000045.1 | repUS2 | | U30316.1 | 1-655 | 96.0 | 98.5 |
| | SPGY01000018.1 | IS66 | ISBf10 | NC_006347 | 88912-91850 | 100 | 100 |
| | | ISBthe1 | IS4 | NC_004663 | 93521-94836 | 99.9 | 98.9 |
| **1001285H_161024_ D4** | NZ_JADNGU010000038.1 | repUS2 | | U30316.1 | 1186-1865 | 99 | 98.5 |
| **14-106904-1** | MKXL01000001 | ISBf5 | IS1182 | NC_006347 | | 100 | 99.9 |
| | MKXL01000067 | ISBf6 | IS5 | AM042593 | 1-950 | 100 | 99.9 |
| **2_1_16** | GG705215.1 | ISBthe1- nimB | IS4 | NC_004663 | 39206-40521 | 99.9 | 98.9 |
| | GG705220.1 | repUS2 | | BFU30316 | 1733-2412 | 99.7 | 98.5 |
| **2_1_56FAA** | GL945044.1 | IS4351 | IS30 | X58717 | 951061-952986 | 100 | 96.8 |
| **2-F-2 #4** | JGDM01000124 | IS613 | IS1380 | AB042549 | 1-1594 | 96.6 | 99.9 |
| | JGDM01000140 | IS613 | IS1380 | AB042549 | 1-1594 | 96.6 | 99.9 |
| | JGDM01000143 | IS613 | IS1380 | AB042549 | 5885-7478 | 96.6 | 99.9 |
| **2-F-2 #5** | JGCN01000047 | IS613 | IS1380 | AB042549 | 29375-30968 | 96.6 | 99.9 |
| | JGCN01000127 | IS613 | IS1380 | AB042549 | 16497-18089 | 96.5 | 99.8 |
| | JGCN01000167 | IS613 | IS1380 | AB042549 | 1-1594 | 96.6 | 99.9 |
| | JGCN01000236 | IS613 | IS1380 | AB042549 | 1-1594 | 96.6 | 99.9 |
| **2-F-2 #7** | JGCO01000059 | IS613 | IS1380 | AB042549 | 1-1594 | 96.6 | 99.9 |
| | JGCO01000205 | IS613 | IS1380 | AB042549 | 1-1594 | 96.6 | 99.9 |
| | JGCO01000278 | IS613 | IS1380 | AB042549 | 29524-31117 | 96.6 | 99.9 |
| **3-F-2 #6** | JGDT01000013 | IS4351 | IS30 | M17124 | 78617-79772 | 99.9 | 99.9 |
| | JGDT01000156 | repUS2 | | BFU30316 | 1061-382 | 99.7 | 98.5 |
| **320_BFRA** | JVLR01000059 | ISBf8 | IS4 | U75371 | 1-1663 | 100 | 99.7 |
| | JVLR01000108 | ISBf5 | IS1182 | NC_006347 | 103993-105822 | 100 | 100 |
| | JVLR01000059 | ISBf8 (Tn4555)-cfxA | IS4 | U75371 | 1-1663 | 100 | 99.7 |
| | JVLR01000108 | ISBf5 | IS1182 | NC_006347 | 103993-105822 | 100 | 100 |
| **321_BFRA** | JVLQ01000062 | ISBf8 | IS4 | U75371 | 4-1666 | 100 | 99.7 |
| | JVLQ01000142 | ISBf5 | IS1182 | NC_006347 | 21028-22857 | 100 | 100 |
| **322_BFRA** | JVLP01000003 | ISBf5 | IS1182 | NC_006347 | 39663-41492 | 100 | 100 |
| **3397 N2** | JGDN01000087 | ISBf5 | IS1182 | NC_006347 | 328-2157 | 100 | 99.5 |
| **3397 N3** | JGEG01000095 | ISBf5 | IS1182 | NC_006347 | 258-2087 | 100 | 99.4 |
| **3397 T10** | JGCP01001284.1 | ISBf5 | IS1182 | NC_006347 | 394-2223 | 100 | 99.5 |
| **3397 T14** | JGDO01000057.1 | ISBf5 | IS1182 | NC_006347 | 249-2078 | 100 | 99.5 |
| **3719 T6** | JGEH01000035 | ISBf1 | IS21 | U05888 | 1267-4052 | 99.9 | 99.9 |
| **3725 D9 ii** | JNHH01000006.1 | ISBthe4 | ISAs1 | NC_004663 | 158862-160211 | 99.8 | 97.3 |
| | JNHH01000013.1 | IS4351 | IS30 | M17124 | 506767-507921 | 100 | 100 |
| | JNHH01000021.1 | IS4351 | IS30 | M17124 | 441579-442733 | 100 | 100 |
| | JNHH01000027.1 | IS4351 | IS30 | M17124 | 61744-62898 | 100 | 100 |
| | JNHH01000031.1 | IS4351 | IS30 | M17124 | 1-1155 | 100 | 100 |

| | | | | | | | |
|---|---|---|---|---|---|---|---|
| | JNHH01000037.1 | ISBthe4 | ISAs1 | NC_004663 | 2310-3659 | 99.8 | 97.3 |
| **3783N2-1** | JGCT01000308 | repUS2 | | BFU30316 | 232-911 | 99.7 | 98.7 |
| **3976T8** | JGDS01000062 | ISBthe5 | IS66 | NC_004663 | 113747-116278 | 98.7 | 93.2 |
| **3986 N(B)19** | JGCW01000128 | ISBf1 | IS21 | U05888 | 1810-4595 | 99.9 | 99.9 |
| | JGCW01000656 | IS613 | IS1380 | AB042549 | 10-1603 | 99.7 | 93.1 |
| **3986T(B)10** | JGCV01000312 | repUS2 | | BFU30316 | 2013-2692 | 99.7 | 98.5 |
| **3988 T1** | JGCZ01000286 | repUS2 | | BFU30316 | 169-848 | 99.7 | 98.5 |
| **3996 N(B) 6** | JGDA01000135 | ISBaov1 | IS1380 | AJ557257 | 44685-46278 | 100 | 99.9 |
| **3998 T(B) 4** | JGDC01000202 | ISBf8 | IS4 | U75371 | 1-1663 | 100 | 99.7 |
| | JGDC01000032 | ISBf8 | IS4 | U75371 | 4-1666 | 100 | 99.7 |
| | JGDC01000133 | ISBf8 | IS4 | U75371 | 14986-16648 | 100 | 99.7 |
| | JGDC01000134 | ISBf8 | IS4 | U75371 | 4-1666 | 100 | 99.7 |
| | JGDC01000274 | ISBf8 | IS4 | U75371 | 22635-24297 | 100 | 99.7 |
| | JGDC01000343 | ISBf8 | IS4 | U75371 | 4-1666 | 100 | 99.7 |
| **638R** | NC_016776.1 | ISBf5 | IS1182 | NC_006347 | 4817558-4819387 | 100 | 99.5 |
| | NC_016776.1 | ISBthe2 | IS1182 | NC_004663 | 4926119-4927960 | 99.6 | 99.0 |
| **86-5443-2-2** | LIDS01000020 | ISBaov1 | IS1380 | AJ557257 | 18167-19760 | 100 | 100 |
| **885_BFRA** | JUPJ01000031 | IS4351 (Tn4351) –tetX | IS30 | M37699.1 | 2258-3424 | 100 | 99.9 |
| | JUPJ01000044 | ISBf1 | IS21 | U05888 | 1-2786 | 99.9 | 99.9 |
| **894_BFRA** | JUOZ01000029 | IS4351 (Tn4351) –tetX | IS30 | M37699.1 | 2258-3424 | 100 | 99.9 |
| | JUOZ01000239 | ISBf1 | IS21 | U05888 | 7127-9912 | 99.9 | 99.9 |
| **915_BFRA** | JUOA01000729 | IS4351 (Tn4351) –tetX | IS30 | M17808 | 3147-3947 | 100 | 100 |
| **A7 (UDC12-2)** | JGEK01000081 | ISBf5 | IS1182 | NC_006347 | 2190-4019 | 100 | 99.8 |
| **AD126T_1B** | VOHZ01000031.1 | repUS2 | | BFU30316 | 1479-2158 | 99.7 | 99.3 |
| | VOIJ01000030.1 | repUS2 | | BFU30316 | 66-745 | 99.7 | 99.3 |
| **AD135F_1B** | VOHV01000015.1 | ISBf1 | IS21 | U05888 | 61445-64230 | 99.9 | 99.9 |
| **AD135F_2B** | WCHZ01000016.1 | ISBf1 | IS21 | U05888 | 1-2786 | 99.9 | 99.9 |
| **AD135F_3B** | VOHT01000015.1 | ISBf1 | IS21 | U05888 | 61445-64230 | 99.9 | 99.9 |
| **AF14-14AC** | QRZO01000083.1 | IS613 | IS1380 | AB042549 | 1-1595 | 96.7 | 100 |
| **AF14-26** | QRZH01000050.1 | IS613 | IS1380 | AB042549 | 1-1595 | 96.7 | 100 |
| **AF32-10** | QRQH01000036.1 | repUS2 | | BFU30316 | 1063-384 | 99.7 | 98.5 |
| | QRQH01000001.1 | ISBf1 | IS21 | U05888 | 85120-87905 | 99.9 | 99.7 |
| **AM13-18** | QRLH01000012.1 | ISBf5 | IS1182 | NC_006347 | 97505-99334 | 100 | 99.9 |
| **AM15-16** | QRKP01000005.1 | ISBf5 | IS1182 | NC_006347 | 105280-107109 | 100 | 100 |
| **AM16-16A** | QRKL01000003.1 | ISBf5 | IS1182 | NC_006347 | 105162-106991 | 100 | 100 |
| **AM17-19** | QRJX01000009.1 | IS612 | IS1380 | AB042548 | 63078-64674 | 99.7 | 93.1 |
| **AM18-6** | QRJE01000005.1 | IS4351 | IS30 | M17124 | 136151-137306 | 99.9 | 99.91 |
| | QRJE01000011.1 | ISBf10 | IS66 | NC_006347 | 1-2939 | 99.9 | 99.7 |
| **AM40-4AC** | QSGL01000025.1 | IS612 | IS1380 | AB042548 | 23061-24658 | 99.7 | 93.1 |
| | QSGL01000011.1 | ISBbi1 | IS1595 | DQ093580 | 188409-189389 | 100 | 99.9 |
| | QSGL01000059.1 | IS616 | IS1380 | AB042560 | 1-1691 | 100 | 100 |
| | QSGL01000060.1 | IS613 | IS1380 | AB042549 | 1-1595 | 96.7 | 100 |

| | | | | | | | |
|---|---|---|---|---|---|---|---|
| **AM47-7** | QSER01000002.1 | ISBf1 | IS21 | U05888 | 253904-256689 | 99.7 | 99.9 |
| **ATCC 25285** | MTGH01000014.1 | ISBf1 | IS21 | U05888 | 154506-157291 | 99.7 | 99.9 |
| **B1 (UDC16-1)** | JGDU01000297 | ISBf5 | IS1182 | NC_006347 | 1042-2871 | 100 | 99.8 |
| **BE1.1** | NZ_LN877293.1 | ISBf5 | IS1182 | NC_006347 | 4744685-4746514 | 100 | 99.4 |
| **BF8 Consensus_4** | LGTH01000004 | IS4351 (Tn4351) – tetX | IS30 | M37699 | 924279-925445 | 100 | 99.9 |
| | LGTH01000004 | IS4351 (Tn4351) – ermF | IS30 | M17124 | 922085-923239 | 100 | 100 |
| | LGTH01000004 | IS4351 | IS30 | M17124 | 83916-85070 | 100 | 100 |
| **BF8 Consensus_1** | LGTH01000001 | IS1168 -nimB | | X71443 | 612426-612920 | 100 | 100 |
| | LGTH01000001 | IS4351 | IS30 | M17124 | 1413422-1414576 | 100 | 100 |
| | LGTH01000001 | IS4351 | IS30 | M17124 | 986563-987717 | 100 | 100 |
| **BF8 Consensus_2** | LGTH01000002 | IS942 -cfiA | IS4 | FM200789 | 1220392-1221141 | 100 | 99.1 |
| | LGTH01000002 | IS4351 | IS30 | M17124 | 320552-321706 | 100 | 100 |
| | LGTH01000002 | IS4351 | IS30 | M17124 | 905526-906680 | 100 | 100 |
| **BF8 Consensus_3** | LGTH01000003 | IS4351 | IS30 | M17124 | 592529-593683 | 100 | 100 |
| **BFG-1** | NZ_CP081922.1 | ISBf1 | IS21 | U05888 | 1288909-1291694 | 99.9 | 99.9 |
| **BFR_KZ03** | SSKJ01000060.1 | ISBf5 | IS1182 | NC_006347 | 1366-3196 | 99.9 | 99.8 |
| **BFR_KZ05** | NZ_JACENG010000047.1 | repUS2 | | BFU30316 | 67-746 | 99.7 | 98.5 |
| | NZ_JACENG010000022.1 | ISBf5 | IS1182 | NC_006347 | 1929-3758 | 100 | 99.8 |
| **BFR_KZ08** | NZ_JACFSU010000010.1 | ISBf1 | IS21 | U05888 | 243181-245966 | 99.9 | 99.9 |
| | NZ_JACFSU010000014.1 | ISBf5 | IS1182 | NC_006347 | 13765-15594 | 100 | 99.6 |
| **BFR_KZ10** | NZ_JACFSW010000006.1 | ISBf5 | IS1182 | NC_006347 | 103488-105317 | 100 | 99.4 |
| **BFR_KZ12** | NZ_JACFSY010000040.1 | repUS2 | | BFU30316 | 1019-1340 | 99.7 | 98.5 |
| | NZ_JACFSY010000037.1 | ISBf5 | IS1182 | NC_006347 | 16163-17992 | 100 | 99.8 |
| **BOB25** | NZ_CP011073.1 | ISBf5 | IS1182 | NC_006347 | 4740323-4742152 | 100 | 100 |
| **CCUG4856T** | NZ_CP036555.1 | ISBf1 | IS21 | U05888 | 2731376-2734161 | 99.9 | 99.9 |
| | | ISBf5 | IS1182 | NC_006347 | 494880-496709 | 100 | 99.8 |
| **CFPLTA004_1B** | VOHY01000054.1 | repUS2 | | BFU30316 | 1741-2420 | 99.7 | 98.5 |
| | VOHY01000019.1 | IS4351 | IS30 | M17124 | 55107-56261 | 100 | 100 |
| | VOHY01000044.1 | ISBthe1 | IS4 | NC_004663 | 5943-7258 | 99.9 | 98.9 |
| **CL04T03C20** | PDCU01000001.1 | ISBf1 | IS21 | U05888 | 374756-377541 | 99.9 | 99.9 |
| | PDCU01000006.1 | ISBf5 | IS1182 | NC_006347 | 312864-314693 | 100 | 99.8 |
| **CL05T00C42** | AKBY01000001 | ISBf4 | IS1182 | NC_006347 | 211444-213294 | 99.3 | 95.6 |
| | AKBY01000002 | ISBf4 | IS1182 | NC_006347 | 898994-900844 | 99.3 | 95.6 |
| | AKBY01000007 | ISBf4 | IS1182 | NC_006347 | 42442-44292 | 99.3 | 95.6 |
| **CL05T12C13** | AGXP01000020 | ISBf4 | IS1182 | NC_006347 | 2473-4323 | 99.3 | 95.6 |
| | AGXP01000022 | ISBf4 | IS1182 | NC_006347 | 163844-165694 | 99.3 | 95.6 |
| | AGXP01000043 | ISBf4 | IS1182 | NC_006347 | 8011-9861 | 99.3 | 95.6 |
| **CL07T00C01** | AGXM01000001 | IS4351 (Tn4351) – ermF | IS30 | M17808.1 | 582395-583195 | 100 | 100 |
| | AGXM01000006 | IS4351 | IS30 | M17124 | 138196-139349 | 99.9 | 99.8 |
| | AGXM01000002 | IS4351 | IS30 | M17124 | 41415-42570 | 99.9 | 99.9 |
| | | IS4351 | IS30 | M17124 | 395702-396856 | 100 | 100 |
| | | IS4351 | IS30 | M17124 | 481839-482993 | 100 | 100 |

| | | | | | | | |
|---|---|---|---|---|---|---|---|
| | AGXM01000004 | IS4351 | IS30 | M17124 | 125743-126897 | 100 | 100 |
| | AGXM01000016 | IS4351 | IS30 | M17124 | 277828-278982 | 100 | 100 |
| **CL07T12C05** | AGXN01000012 | IS4351 | IS30 | M17124 | 480566-481720 | 100 | 100 |
| | | IS4351 | IS30 | M17124 | 85078-86232 | 100 | 100 |
| | AGXN01000005 | IS4351 | IS30 | M17124 | 959008-960162 | 100 | 100 |
| | | IS4351 | IS30 | M17124 | 845586-846740 | 100 | 100 |
| | AGXN01000007 | IS4351 | IS30 | M17124 | 51266-52420 | 100 | 100 |
| | AGXN01000022 | IS4351 | IS30 | M17124 | 140193-141347 | 100 | 100 |
| **COR2-248-WT-1** | NZ_JABAGK010000001.1 | ISBf1 | IS21 | U05888 | 444473-447258 | 99.9 | 99.9 |
| **DCMOUH0017B** | JMZY02000010 | ISBf10 | IS66 | NC_006347 | 1-2939 | 99.9 | 96.4 |
| **DCMOUH0018B** | JMZZ02000154 | IS4351 | IS30 | M17124 | 88724-89879 | 99.9 | 99.8 |
| | JMZZ02000029 | ISBbi1 | IS1595 | DQ093580 | 9943-10923 | 100 | 99.9 |
| **DCMOUH0042B** | JPGQ01000091 | ISBbi1 | | DQ093580 | 52955-53935 | 100 | 99.0 |
| **DCMOUH0067B** | JPHS01000166 | IS4351 | IS30 | M17124 | 1-1155 | 100 | 100 |
| **DCMOUH0085B** | JPHP01000036 | IS614 | IS1380 | AB042559 | 187690-189285 | 100 | 100 |
| **DK1985** | NZ_CP044428.1 | ISBf3 | IS1182 | NC_006297 | 4342106-4343946 | 99.8 | 99.8 |
| | NZ_CP044428.1 | ISBf3 | IS1182 | NC_006297 | 1389635-1391479 | 100 | 99.9 |
| | NZ_CP044428.1 | ISBf3 | IS1182 | NC_006297 | 5209506-5211350 | 100 | 100 |
| | NZ_CP044428.1 | ISBf3 | IS1182 | NC_006297 | 3692513-3694357 | 100 | 100 |
| | NZ_CP044428.1 | ISBf3 | IS1182 | NC_006297 | 3472411-3474255 | 100 | 100 |
| | NZ_CP044428.1 | ISBf3 | IS1182 | NC_006297 | 1965583-1967427 | 100 | 100 |
| | NZ_CP044428.1 | ISBf3 | IS1182 | NC_006297 | 1766997-1768841 | 100 | 100 |
| | NZ_CP044428.1 | ISBf3 | IS1182 | NC_006297 | 1588979-1590823 | 100 | 100 |
| | NZ_CP044428.1 | ISBf3 | IS1182 | NC_006297 | 1453002-1454846 | 100 | 100 |
| | NZ_CP044428.1 | ISBf3 | IS1182 | NC_006297 | 978372-980216 | 100 | 100 |
| | NZ_CP044428.1 | ISBf3 | | NC_006297 | 625825-627669 | 100 | 100 |
| **DS-166** | JGDD01000090 | ISBf5 | IS1182 | NC_006347 | 286324-288153 | 100 | 99.7 |
| **Ds-233** | JGDG01000279 | ISBf1 | IS21 | U05888 | 18451-21236 | 99.9 | 99.9 |
| **DS-71** | JGDF01000150 | ISBf5 | IS1182 | NC_006347 | 4426-6255 | 100 | 99.9 |
| **GUT04** | NZ_CP043610.1 | ISBf1 | IS21 | U05888 | 2483317-2486102 | 99.9 | 99.9 |
| | NZ_CP043610.1 | ISBf5 | IS1182 | NC_006347 | 4007623-4009452 | 100 | 99.9 |
| **HAP130N_2B** | VOIS01000009.1 | ISBf1 | IS21 | U05888 | 36572-39357 | 99.9 | 99.9 |
| | WCIE01000037.1 | repUS2 | | BFU30316 | 1-439 | 64.5 | 99.8 |
| **HCK-B3** | QRBX01000010.1 | ISBf1 | IS21 | U05888 | 11267-14052 | 99.9 | 99.9 |
| **HMW 610** | AGXQ01000032 | IS613 | IS1380 | AB042549 | 1929-3523 | 96.7 | 100 |
| | AGXQ01000023 | ISBthe6 | IS66 | NC_004663 | 46030-48566 | 99.7 | 100 |
| | AGXQ01000023 | ISBthe6 | IS66 | NC_004663 | 41826-44362 | 99.7 | 100 |
| | AGXQ01000023 | IS613 | IS1380 | AB042549 | 119136-120730 | 96.68 | 100 |
| | AGXQ01000008 | ISBbi1 | IS1595 | DQ093580 | 39433-40413 | 100 | 99.9 |
| | AGXQ01000012 | IS613 | IS1380 | NC_004663 | 57461-59055 | 96.7 | 100 |
| | AGXQ01000012 | IS613 | IS1380 | AB042549 | 232461-234055 | 96.7 | 100 |
| | AGXQ01000039 | ISBf6 | IS5 | AM042593 | 6917-7866 | 100 | 99.9 |

| | | | | | | | |
|---|---|---|---|---|---|---|---|
| | AGXQ01000002 | ISBthe6 | IS66 | NC_004663 | 1-2544 | 100 | 100 |
| | AGXQ01000002 | IS613 | IS1380 | AB042549 | 317011-318605 | 96.7 | 100 |
| | AGXQ01000003 | IS613 | IS1380 | AB042549 | 37955-39549 | 96.7 | 100 |
| | AGXQ01000026 | ISBf5 | IS1182 | NC_006347 | 237410-239239 | 100 | 99.9 |
| **HMW 615** | AGXR01000023 | IS612 | IS1380 | AB042548 | 42095-43692 | 99.7 | 98.2 |
| | AGXR01000022 | ISBbi1 | IS1595 | DQ093580 | 178822-179802 | 100 | 99.9 |
| | AGXR01000022 | IS613 | IS1380 | AB042549 | 58531-60125 | 96.7 | 99.9 |
| | AGXR01000005 | IS612 | IS1380 | AB042548 | 728199-729795 | 99.7 | 97.9 |
| | AGXR01000016 | IS613 | IS1380 | AB042549 | 240865-242459 | 96.7 | 99.9 |
| | AGXR01000017 | IS613 | IS1380 | AB042549 | 110593-112187 | 96.7 | 100 |
| | AGXR01000019 | IS612 | IS1380 | AB042548 | 353764-355361 | 99.6 | 99.5 |
| | AGXR01000019 | IS612 | IS1380 | AB042548 | 83002-84599 | 99.6 | 99.5 |
| | AGXR01000021 | IS612 | IS1380 | AB042548 | 57583-59180 | 99.6 | 99.5 |
| | AGXR01000026 | IS612 | IS1380 | AB042548 | 157308-158905 | 99.6 | 99.4 |
| | AGXR01000031 | IS612 | IS1380 | AB042548 | 154896-156493 | 99.6 | 99.5 |
| | AGXR01000031 | IS613 | IS1380 | AB042549 | 151680-153274 | 96.7 | 100 |
| **HMW 616** | ALOB01000009 | IS613 | IS1380 | AB042549 | 248610-250204 | 96.7 | 100 |
| | ALOB01000013 | ISBf5 | IS1182 | NC_006347 | 126222-128051 | 100 | 99.8 |
| **I1345** | JGEW01000032 | ISBf1 | IS21 | U05888 | 70210-72995 | 99.9 | 99.9 |
| **J1101437_171009_F3** | JADNMB010000013.1 | ISBf1 | IS21 | U05888 | 67234-70019 | 99.9 | 99.9 |
| **KLE1257** | KQ970992.1 | ISBf1 | IS21 | U05888 | 202490-205275 | 99.9 | 99.9 |
| | KQ971004.1 | ISBf5 | IS1182 | NC_006347 | 102229-104058 | 100 | 99.6 |
| **Korea 419** | JGDW01000109 | ISBf5 | IS1182 | NC_006347 | 4345-6174 | 100 | 99.8 |
| **MC1** | CAAKNW010000064.1 | ISBf5 | IS1182 | NC_006347 | 13944-15773 | 100 | 99.9 |
| **MGYG-HGUT-00236** | CABJEQ010000009.1 | IS612 | IS1380 | AB042548 | 63078-64674 | 99.7 | 97.8 |
| **NCTC 9343 strain ATCC 25285** | NC_003228.3 | ISBf1 | IS21 | U05888 | 1795167-1797952 | 99.9 | 99.9 |
| | NC_003228.3 | ISBf5 | IS1182 | NC_006347 | 4763811-4765640 | 100 | 99.8 |
| **O:21** | KV751178.1 | IS4351 | IS30 | M17124 | 477666-478820 | 100 | 100 |
| | KV751174.1 | IS4351 | IS30 | M17124 | 320642-321795 | 99.91 | 100 |
| **OF01-1** | QSDG01000002.1 | ISBf5 | IS1182 | NC_006347 | 430864-432693 | 100 | 99.8 |
| | QSDG01000050.1 | IS613 | IS1380 | AB042549 | 1-1595 | 96.7 | 100 |
| **OF03-7** | QSCG01000179.1 | repUS2 | | BFU30316 | 1647-2326 | 99.7 | 98.5 |
| | QSCG01000021.1 | ISBf5 | IS1182 | NC_006347 | 21398-23227 | 100 | 99.8 |
| | QSCG01000184.1 | ISBthe6 | IS66 | NC_004663 | 1-2544 | 100 | 100 |
| **OF05-11AC** | QSWE01000001.1 | ISBaov1 | IS1380 | AJ557257 | 732507-734100 | 100 | 100 |
| **OF05-13AC** | QSWB01000001.1 | ISBaov1 | IS1380 | AJ557257 | 732487-734080 | 100 | 100 |
| **OM02-11** | QSVT01000020.1 | IS612 | IS1380 | AB042548 | 1-1574 | 98.0 | 97.9 |
| **OM02-9** | QSVD01000016.1 | IS612 | IS1380 | AB042548 | 1-1597 | 99.4 | 97.9 |
| **OM04-9BH** | QSUS01000002.1 | ISBf5 | IS1182 | NC_006347 | 764419-766248 | 100 | 99.8 |
| | QSUS01000033.1 | repUS2 | | BFU30316 | 1778-2457 | 99.7 | 98.5 |
| | QSUS01000034.1 | ISBthe6 | IS66 | NC_004663 | 1-2544 | 100 | 100 |
| **OM06-1** | QSUD01000017.1 | repUS2 | | BFU30316 | 1063-384 | 99.71 | 98.5 |
| | QSUD01000005.1 | ISBf5 | IS1182 | NC_006347 | 328846-330675 | 100 | 99.7 |
| **OM06-30AC** | QSTV01000022.1 | repUS2 | | BFU30316 | 2014-2693 | 99.7 | 98.5 |
| **OM06-30AC** | QSTV01000011.1 | ISBf5 | IS1182 | NC_006347 | 93700-95529 | 100 | 99.7 |
| **Q1F2** | NZ_CP018937.1 | IS613 | IS1380 | AB042549 | 5173821- | 96.7 | 100 |

| | | | | | | | |
|---|---|---|---|---|---|---|---|
| | | | | | 5175415 | | |
| | NZ_CP018937.1 | IS613 | IS1380 | AB042549 | 28633-30227 | 96.7 | 100 |
| | NZ_CP018937.1 | IS613 | IS1380 | AB042549 | 512603-514197 | 96.7 | 100 |
| **S13 L11** | JGDI01000124 | IS613 | IS1380 | AB042549 | 22608-24201 | 99.7 | 93.1 |
| | JGDI01000131 | IS613 | IS1380 | AB042549 | 1-1594 | 99.7 | 93.1 |
| | JGDI01000322 | IS613 | IS1380 | AB042549 | 1-1594 | 99.7 | 93.1 |
| | JGDI01000414 | IS613 | IS1380 | AB042549 | 1-1594 | 93.1 | 100 |
| | JGDI01000564 | IS613 | IS1380 | AB042549 | 4721-6314 | 93.1 | 100 |
| **S24L15** | JGEM01000086 | ISBf5 | IS1182 | NC_006347 | 105273-107102 | 100 | 99.9 |
| | JGEM01000086 | ISBf5 | IS1182 | NC_006347 | 105273-107102 | 100 | 99.9 |
| **S24L26** | JGEN01000067 | ISBf5 | IS1182 | NC_006347 | 105520-107349 | 100 | 100 |
| **S24L34** | JGEO01000061 | ISBf5 | IS1182 | NC_006347 | 105522-107351 | 100 | 99.9 |
| **S36L11** | JGDJ01000031 | ISBthe1 | IS4 | NC_004663 | 4422-5737 | 99.9 | 98.9 |
| | JGDJ01000062 | ISBthe1 | IS4 | NC_004663 | 2-1317 | 99.9 | 98.9 |
| | JGDJ01000076 | ISBthe1 | IS4 | NC_004663 | 5-1320 | 99.9 | 98.9 |
| | JGDJ01000289 | ISBthe1 | IS4 | NC_004663 | 47913-49228 | 99.9 | 98.9 |
| **S36L12** | JGEP01000004 | ISBthe1 | IS4 | NC_004663 | 5-1320 | 99.9 | 98.9 |
| | JGEP01000035 | ISBthe1 | IS4 | NC_004663 | 159408-160723 | 99.9 | 98.9 |
| | JGEP01000089 | ISBthe1 | IS4 | NC_004663 | 49502-50817 | 99.9 | 98.9 |
| | JGEP01000090 | ISBthe1 | IS4 | NC_004663 | 2-1317 | 99.9 | 98.9 |
| **S36L5** | JGEQ01000027 | ISBthe1 | IS4 | NC_004663 | 5-1320 | 99.9 | 98.9 |
| | JGEQ01000050 | ISBthe1 | IS4 | NC_004663 | 143858-145173 | 99.9 | 98.9 |
| | JGEQ01000103 | ISBthe1 | IS4 | NC_004663 | 47913-49228 | 99.9 | 98.9 |
| | JGEQ01000104 | ISBthe1 | IS4 | NC_004663 | 2017-3332 | 99.9 | 98.9 |
| **TF05-31** | QSSJ01000009.1 | ISBf5 | IS1182 | NC_006347 | 118116-119945 | 100 | 99.9 |
| **TF09-1** | QSRG01000016.1 | ISBf5 | IS1182 | NC_006347 | 55857-57686 | 100 | 99.9 |
| **TM08-15** | QSOR01000002.1 | ISBf1 | IS21 | U05888 | 85158-87943 | 99.9 | 99.8 |
| | QSOR01000010.1 | ISBf5 | IS1182 | NC_006347 | 103118-104947 | 100 | 99.6 |
| **US326** | PDCS01000003.1 | ISBf5 | IS1182 | NC_006347 | 82252-84081 | 100 | 99.7 |
| **YCH46 strain 82A12** | NZ_UYXF00000000.1 | IS616 | IS1380 | AB042560 | 1-1691 | 100 | 100 |
