## Supplementary material for "Comprehensive Comparative Genomic revels: *Bacteroides fragilis* is a reservoir of antibiotic resistance genes in the gut microbiota": S7

**Table S7. Results obtained by Hyphy software.**

| Gene | NCBI Annotation | BUSTED | | FEL | | FUBAR | | MEME | | SLAC | |
|---|---|---|---|---|---|---|---|---|---|---|---|
| | | dN/dS | Proportion (%) | Site | P-value | Site | Posterior Probability | Site | P-value | Site | P-value |
| *AADK* | Aminoglycoside 6-adenylyltransferase | 5.177 | 2.161 | | | | | 91 | 0.031 | | |
| *ABFA* | Intracellular exo-alpha-(1→exo-alpha-(1->5)-L-arabinofuranosidase | 1.216 | 14.744 | 593<br>725<br>1188 | 0.046<br>0.011<br>0.010 | 210<br>224<br>593<br>725<br>1188 | 0.966<br>0.981<br>0.989<br>0.997<br>0.998 | 210<br>224<br>725<br>861<br>926<br>1188 | 0.030<br>0.032<br>0.006<br>0.021<br>0.047<br>0.016 | | |
| *ABGT* | p-aminobenzoyl-glutamate transport protein | 4.008 | 4.478 | 448<br>479 | 0.046<br>0.032 | 225<br>479 | 0.962<br>0.988 | 365<br>448<br>479 | 0.021<br>0.008<br>0.009 | | |
| *ACKA* | Acetate kinase | 40.472 | 0.376 | | | | | | | | |
| *ACM* | Lysozyme M1 | 14.567 | 0.000 | 202 | 0.015 | 96<br>202<br>278 | 0.989<br>0.996<br>0.987 | 202 | 0.024 | | |
| *ACND* | 2-methylcitrate dehydratase | 3.939 | 3.12 | | | | | 574<br>736 | 0.037<br>0.032 | | |
| *ACSA* | Acetyl-coenzyme A synthetase | 8.061 | 2.192 | 59<br>274 | 0.050<br>0.032 | 59<br>274 | 0.992<br>0.989 | 274 | 0.002 | | |
| *ADH* | Alcohol dehydrogenase | | | | | 254<br>255 | 0.966<br>0.981 | | | | |
| *AMIA* | N-acetylmuramoyl-L-alanine amidase | | | | | 280 | 0.983 | | | | |
| *AMYB* | Alpha-amylase | 3.675 | 4.011 | 541 | 0.049 | 541 | 0.963 | 346 | 0.010 | | |
| *APRX* | Serine protease | 1.288 | 8.033 | 401 | 0.005 | 401<br>426 | 0.999<br>0.979 | 293<br>401 | 0.001<br>0.008 | 401 | 0.039 |

| | | | | | | | | | | | |
|---|---|---|---|---|---|---|---|---|---|---|---|
| *APT* | Adenine phosphoribosyltransferase | 4.125 | 2.615 | | | | | | | | |
| *ARAC_1* | Arabinose operon regulatory protein | | | | | | | 139 | 0.009 | | |
| *ARBA* | Extracellular exo-alpha-(1->5)-L-arabinofuranosidase | 45.924 | 0.836 | 3 | 0.045 | 3<br>129<br>233 | 0.994<br>0.970<br>0.973 | 3<br>122 | 0.000<br>0.002 | | |
| *ARGB* | Acetylglutamate kinase | | | 81 | 0.001 | 81 | 0.9988 | 81 | 0.001 | 81 | 0.017 |
| *ARGG* | Argininosuccinate synthase | 70.446 | 0.393 | 339 | 0.032 | 339 | 0.994 | 258<br>339 | 0.044<br>0.046 | | |
| *AROF* | Phospho-2-dehydro-3-deoxyheptonate aldolase | 7.144 | 1.055 | | | 197 | 0.992 | 197 | 0 | | |
| *AROQ* | 3-dehydroquinate dehydratase | 11.192 | 1.887 | | | 32 | 0.963 | 137 | 0.023 | | |
| *ARTP* | Arginine-binding extracellular protein | 5.232 | 3.026 | 32 | 0.023 | 32 | 0.992 | 32 | 0.044 | | |
| *ASNS* | Asparagine--tRNA ligase | 1.681 | 2.935 | | | 86 | 0.957 | | | | |
| *ATPG* | ATP synthase gamma chain | 3.432 | 5.65 | 272 | 0.023 | 272 | 0.990 | 272 | 0.003 | | |
| *ATSA* | Arylsulfatase | 21.218 | 2.875 | | | 22<br>244<br>270 | 0.975<br>0.981<br>0.961 | 245 | 0.022 | | |
| *ATSA_7* | Arylsulfatase | 4.508 | 3.108 | 375<br>445 | 0.001<br>0.027 | 302<br>375<br>445 | 0.965<br>1.000<br>0.995 | 375<br>445 | 0.000<br>0.005 | 375 | 0.005 |
| *ASPT_4* | Aspartate/alanine antiporter | | | | | 465 | 0.976 | | | | |
| *AXEA1_4* | Acetylxylan esterase | 2.344 | 11.008 | 203<br>286<br>816 | 0.005<br>0.043<br>0.026 | 203<br>286<br>528<br>816 | 0.999<br>0.992<br>0.964<br>0.995 | 203<br>816 | 0.009<br>0.018 | | |
| *BAMA_3* | Outer membrane protein assembly factor | | | | | | | 187<br>390 | 0.036<br>0.040 | | |
| *BAMB* | Outer membrane protein assembly factor | 7.345 | 6.107 | | | 62<br>374<br>528<br>601 | 0.963<br>0.976<br>0.981<br>0.955 | | | | |
| *BAMD_2* | Outer membrane protein assembly factor | 2.713 | 2.787 | | | 246 | 0.979 | | | | |
| *BARA_1* | Signal transduction histidine-protein kinase BarA | | | | | 310 | 0.978 | | | | |

| | | | | | | | | | | | |
|---|---|---|---|---|---|---|---|---|---|---|---|
| *BCRC* | Hypothetical protein | | | | | | | 198 | 0.054 | | |
| *BEPA_1* | Hypothetical protein | | | 594 | 0.005 | 594 | 0.994 | 282<br>594 | 0.038<br>0.004 | 594 | 0.039 |
| *BEPC_1* | Outer membrane efflux protein | | | | | 172 | 0.9839 | 172 | 0.006 | | |
| *BEPE_2* | Efflux pump membrane transporter | 2.846 | 1.872 | 189<br>191 | 0.011<br>0.033 | 189<br>191 | 0.999<br>0.994 | 189<br>191<br>588 | 0.000<br>0.000<br>0.025 | | |
| *BEPE_3* | Efflux pump membrane transporter | 7.674 | 1.266 | 534<br>1033 | 0.031<br>0.039 | 534<br>847<br>1033 | 0.9907<br>0.9856<br>0.9869 | 188<br>482<br>534<br>756<br>847 | 0.051<br>0.042<br>0.024<br>0.051<br>0.008 | | |
| *BFMBAB* | 2-oxoisovalerate dehydrogenase subunit beta | 2.592 | 2.075 | | | | | 656 | 0.055 | | |
| *BGLB_4* | Thermostable beta-glucosidase B | | | 841 | 0.046 | 16<br>654<br>782<br>841 | 0.963<br>0.989<br>0.965<br>0.991 | | | | |
| *BIPA* | GTP-binding protein TypA/BipA | 4.745 | 1.617 | 418 | 0.013 | 418 | 0.9934 | 418 | 0.002 | | |
| *BMRU* | Putative lipid kinase | 4.666 | 1.466 | | | | | | | | |
| *BCR_1* | Bicyclomycin resistance protein | 1.342 | 20.497 | 103 | 0.021 | 103<br>210<br>299<br>387 | 0.995<br>0.980<br>0.966<br>0.966 | 72<br>103<br>210 | 0.045<br>0.032<br>0.005 | | |
| *BSAA* | Glutathione peroxidase | | | | | 86<br>179 | 0.954<br>0.981 | | | | |
| *BTR_2* | HTH-type transcriptional activator | 8.991 | 0.84 | | | 73 | 0.973 | | | | |
| *CARB_1* | Carbamoyl-phosphate synthase large chain | 1.001 | 1.822 | | | 881 | 0.954 | 881 | 0.0201 | | |
| *CBGA* | Beta-galactosidase | | | 309<br>311 | 0.019<br>0.014 | 309<br>311 | 0.998<br>0.998 | 309<br>311 | 0.029<br>0.022 | 311 | 0.039 |

| | | | | | | | | | | | |
|---|---|---|---|---|---|---|---|---|---|---|---|
| *CCSA_1* | Cytochrome c biogenesis protein | 27.082 | 1.354 | | | 237 | 0.952 | 18<br>29<br>33<br>237 | 0.022<br>0.003<br>0.005<br>0.011 | | |
| *CDD* | Cytidine deaminase | 10.904 | 1.724 | | | 91 | 0.951 | | | | |
| *CDSA* | Phosphatidate cytidylyltransferase | 1 | 8.588 | 279 | 0.033 | 279 | 0.995 | 279 | 0.048 | | |
| *CEPA* | cepA family class A extended-spectrum beta-lactamase | | | | | 88 | 0.9878 | | | | |
| *CIRA_1* | Colicin I receptor | 55.055 | 0.919 | 6 | 0.019 | 6<br>31<br>320<br>730 | 0.997<br>0.978<br>0.960<br>0.964 | 1 | 0.030 | | |
| *CIRA_3* | Colicin I receptor | 3.444 | 2.061 | 372 | 0.020 | 372 | 0.996 | 333<br>372<br>490 | 0.026<br>0.006<br>0.025 | | |
| *CIRA_4* | Colicin I receptor | | | | | 499 | 0.983 | | | | |
| *CIRA_5* | Colicin I receptor | 2.766 | 3.976 | 11 | 0.0112 | 2<br>11<br>586 | 0.973<br>0.993<br>0.976 | 11<br>29<br>320<br>451<br>518<br>586 | 0.000<br>0.000<br>0.037<br>0.051<br>0.034<br>0.042 | 11 | 0.012 |
| *CIRA_7* | Colicin I receptor | | 1.413 | 6.791 | | | | 226<br>279 | 0.045<br>0.002 | | |
| *CIRA_8* | Colicin I receptor | 530.567 | 0.096 | 516<br>731 | 0.026<br>0.036 | 252<br>378<br>391<br>516<br>672<br>731<br>750 | 0.974<br>0.959<br>0.979<br>0.990<br>0.960<br>0.991<br>0.958 | 431<br>516<br>692 | 0.011<br>0.039<br>0.002 | | |

| | | | | | | | | | | | |
|---|---|---|---|---|---|---|---|---|---|---|---|
| *CIRA_10* | Colicin I receptor | 1.087 | 6.115 | 324 | 0.044 | 324<br>452<br>674 | 0.988<br>0.972<br>0.971 | 324<br>452<br>586<br>674 | | | |
| *CLCA* | H(+)/Cl(-) exchange transporter | 1.815 | 3.927 | 16 | 0.001 | 16 | 0.999 | 16 | 0 | 16 | 0.017 |
| *CLCB* | Voltage-gated ClC-type chloride channel | 1.447 | 1.403 | | | 508 | 0.978 | 172 | 0.046 | | |
| *CLOSI* | Clostripain | 31.377 | 1.055 | | | 340<br>372 | 0.975<br>0.986 | | | | |
| *CLPB* | Chaperone protein | 4002,8 | 0.054 | | | 83<br>86<br>484<br>509 | 0.979<br>0.965<br>0.958<br>0.972 | 83<br>626 | 0.000<br>0.049 | | |
| *CLPB1* | Chaperone protein ClpB 1 | 407.747 | 0.037 | | | 796 | 0.981 | 733<br>796 | 0.001<br>0.022 | | |
| *CLSA_1* | Major cardiolipin synthase ClsA | 9.501 | 0.933 | 101 | 0.979 | 93 | 0.022 | | | | |
| *CMK_2* | Cytidylate kinase | 3.944 | 4.047 | 78 | 0.005 | 78 | 0.999 | 78 | 0.000 | 78 | 0.005 |
| *COABC* | Coenzyme A biosynthesis bifunctional protein | 25.405 | 0.676 | | | 188 | 0.957 | | | | |
| *COBD_2* | Threonine-phosphate decarboxylase | 2.025 | 9.595 | 14<br>127 | 0.0115<br>0.0161 | 14<br>127 | 0.998<br>0.996 | 14<br>127<br>321 | 0.012<br>0.002<br>0.047 | | |
| *COBP* | Bifunctional adenosylcobalamin biosynthesis protein | 6.744 | 7.697 | 21 | 0.000 | 21<br>45<br>93<br>94<br>139 | 0.999<br>0.982<br>0.962<br>0.977<br>0.953 | 21<br>45<br>93<br>98 | 0.001<br>0.038<br>0.028<br>0.038 | 21 | 0.001 |
| *COBS* | Adenosylcobinamide-GDP ribazoletransferase | 1.016 | 0.000 | | | 42<br>202 | 0.985<br>0.977 | | | | |
| *CRNA* | Creatinine amidohydrolase | | | | | 149 | 0.963 | | | | |
| *CTPB_2* | Carboxy-terminal processing protease | 3.913 | 1.987 | 9 | 0.0422 | 9<br>141<br>466 | 0.993<br>0.972<br>0.964 | 3<br>9<br>141 | 0.027<br>0.008<br>0.006 | | |

| | | | | | | | | | | | |
|---|---|---|---|---|---|---|---|---|---|---|---|
| *CUSA* | Cation efflux system protein | 3.364 | 4.245 | | | 848 | 0.977 | 41<br>295<br>304<br>667<br>668<br>848<br>862<br>1055<br>1164 | 0.032<br>0.044<br>0.031<br>0.026<br>0.040<br>0.000<br>0.024<br>0.007<br>0.043 | | |
| *CVFB* | Conserved virulence factor B | | | 223 | 0.042 | 223 | 0.984 | | | | |
| *CYSC* | Adenylyl-sulfate kinase | 9.359 | 1.224 | | | 4 | 0.964 | 4 | 0.044 | | |
| *CYSD* | Sulfate adenylyltransferase subunit 2 | 2457.895 | 0.066 | | | | | 101<br>122 | 0.019<br>0.033 | | |
| *CYSN* | Sulfate adenylyltransferase subunit 1 | 1.837 | 4.699 | 323 | 0.023 | 323 | 0.995 | 323<br>442 | 0.009<br>0.042 | | |
| *CZCA_1* | Cobalt-zinc-cadmium resistance protein | | | 1023 | 0.025 | 776<br>888<br>1023 | 0.960<br>0.985<br>0.993 | 1023 | 0.038 | 1023 | 0.031 |
| *DADD* | 5'-deoxyadenosine deaminase | 6.153 | 2.974 | 166 | 0.013 | 166 | 0.996 | 22<br>166<br>341<br>376<br>407<br>414 | 0.024<br>0.002<br>0.009<br>0.011<br>0.048<br>0.028 | | |
| *CZCA_3* | Cobalt-zinc-cadmium resistance protein CzcA | 2.66 | 4.714 | 280<br>368<br>897 | 0.001<br>0.022<br>0.012 | 280<br>368<br>897 | 0.999<br>0.992<br>0.998 | 61<br>104<br>178<br>280<br>304<br>368<br>897 | 0.027<br>0.018<br>0.025<br>0.000<br>0.022<br>0.002<br>0.001 | 280<br>897 | 0.004<br>0.034 |
| *DAP4* | Dipeptidyl aminopeptidase 4 | 3.862 | 2.28 | 382 | 0.041 | 382 | 0.997 | 110<br>382 | 0.007<br>0.021 | | |

| | | | | | | | | | | | |
|---|---|---|---|---|---|---|---|---|---|---|---|
| *DAPE_1* | Putative succinyl-diaminopimelate desuccinylase | 25.126 | 2.134 | | | 23 | 0.959 | 19<br>23<br>241<br>266 | 0.044<br>0.031<br>0.049<br>0.042 | | |
| *DAPL_2* | LL-diaminopimelate aminotransferase | 38.342 | 1.067 | 391 | 0.015 | 197<br>203<br>299<br>391 | 0.985<br>0.983<br>0.984<br>0.994 | 203<br>391 | 0.003<br>0.022 | 391 | 0.049 |
| *DCUA* | Anaerobic C4-dicarboxylate transporter | 21.3 | 0.283 | 420 | 0.031 | 420 | 0.989 | 203<br>420 | 0.024<br>0.003 | | |
| *DDL* | D-alanine--D-alanine ligase | | | 199 | 0.030 | 199 | 0.996 | 52<br>199 | 0.019<br>0.043 | | |
| *DEOB* | Phosphopentomutase | | | | | 30<br>225 | 0.965<br>0.992 | | | | |
| *DGT* | Deoxyguanosinetriphosphate triphosphohydrolase | 1.753 | 0.000 | 135 | 0.011 | 88<br>135<br>398 | 0.964<br>0.997<br>0.963 | 135 | 0.018 | | |
| *DINB_1* | DNA polymerase IV | 10.052 | 1.435 | | | 199 | 0.964 | 199<br>200 | 0.011<br>0.048 | | |
| *DNAA* | Chromosomal replication initiator protein | 6.902 | 0.86 | | | | | | | | |
| *DNAE* | DNA polymerase III subunit alpha | 2.807 | 2.42 | 461 | 0.000 | 13<br>461 | 0.986<br>1 | 461 | 0.000 | 461 | 0 |
| *DNAX_1* | DNA polymerase III subunit tau | 17.609 | 1.682 | 380<br>546 | 0.032<br>0.019 | 380<br>385<br>486<br>546 | 0.992<br>0.982<br>0.977<br>0.996 | 374<br>380<br>385<br>546 | 0.001<br>0.003<br>0.004<br>0.005 | | |
| *DNAX_2* | DNA polymerase III subunit tau | 4.256 | 4.908 | 14 | 0.037 | 14 | 0.992 | 73 | 0.048 | | |
| *DPP7_1* | Dipeptidyl-peptidase 7 | 13.499 | 2.012 | 8 | 0.004 | 8<br>383<br>469 | 0.999<br>0.987<br>0.976 | 8<br>469 | 0.000<br>0.013 | 8 | 0.017 |

| | | | | | | | | | | | |
|---|---|---|---|---|---|---|---|---|---|---|---|
| *DPP7_2* | Dipeptidyl-peptidase 7 | | | 280 | 0.002 | 280<br>379 | 0.9992<br>0.9804 | 280 | 0.001 | 280 | 0.008 |
| *DPP7_3* | Dipeptidyl-peptidase 7 | | | 524<br>565 | 0.008<br>0.034 | 32<br>413<br>524<br>565 | 0.970<br>0.976<br>0.995<br>0.991 | 524<br>565 | 0.014<br>0.049 | 524 | 0.043 |
| *DSBD* | Thiol:disulfide interchange protein | 3.25 | 4.18 | 181 | 0.016 | 6<br>181 | 0.969<br>0.993 | 6<br>147<br>181 | 0.016<br>0.047<br>0.024 | | |
| *DXS_1* | 1-deoxy-D-xylulose-5-phosphate synthase | 1.801 | 5.846 | 350<br>633 | 0.002<br>0.029 | 350<br>633 | 0.999<br>0.991 | 350<br>526<br>565<br>597<br>633 | 0.000<br>0.041<br>0.020<br>0.020<br>0.013 | 350 | 0.01 |
| *EAM* | Glutamate 2,3-aminomutase | 2.049 | 8.17 | 52<br>286<br>451<br>616 | 0.044<br>0.033<br>0.008<br>0.031 | 52<br>286<br>451<br>616 | 0.989<br>0.987<br>0.998<br>0.990 | 52<br>87<br>286<br>451<br>474<br>558<br>616<br>624<br>692 | 0.016<br>0.016<br>0.011<br>0.011<br>0.035<br>0.043<br>0.013<br>0.012<br>0.019 | | |
| *EMRB* | Multidrug export protein EmrB | 3.518 | 3.109 | 463<br>499 | 0.009<br>0.045 | 186<br>463<br>499 | 0.979<br>0.998<br>0.989 | 463<br>499 | 0.004<br>0.019 | | |
| *ENVC* | Murein hydrolase activator | | | 63 | 0.010 | 18<br>63<br>95<br>126<br>257 | 0.975<br>0.998<br>0.977<br>0.973<br>0.961 | 63 | 0.017 | | |

| | | | | | | | | | |
|---|---|---|---|---|---|---|---|---|---|
| *ERA* | GTPase | | | | | 135 | 0.962 | | |
| *EMRA_1* | Multidrug export protein | 6.631 | 3.063 | 314 | 0.029 | 314 | 0.992 | 314 | 0.042 |
| *ESIB* | Secretory immunoglobulin A-binding protein | | | | | 12<br>39<br>378<br>425 | 0.985<br>0.937<br>0.962<br>0.976 | | |
| *ETP* | Low molecular weight protein-tyrosine-phosphatase | 1.323 | 0.000 | 75<br>117 | 0.014<br>0.033 | 75<br>117 | 0.998<br>0.994 | 75<br>117 | 0.022<br>0.044 |
| *ETTA_1* | Energy-dependent translational throttle protein | 4.08 | 2.443 | 2<br>35 | 0.006<br>0.031 | 2<br>35<br>39 | 0.999<br>0.995<br>0.993 | 2<br>35<br>39 | 0.001<br>0.006<br>0.014 |
| *EVGS* | Sensor protein EvgS | | | 704 | 0.011 | 704 | 0.997 | 704 | 0.001 |
| *EXO_I_3* | Beta-hexosaminidase | 8 | 0.028 | 8<br>113<br>651 | 0.992<br>0.961<br>0.985 | 8<br>376<br>600<br>630<br>657<br>690 | 0.041<br>0.038<br>0.033<br>0.030<br>0.012<br>0.022 | | |
| *EXO_I_6* | Beta-hexosaminidase | 1.636 | 8.439 | 603 | 0.048 | 238<br>292<br>603 | 0.986<br>0.985<br>0.991 | 35<br>292<br>673 | 0.010<br>0.016<br>0.019 |
| *FABD* | Malonyl CoA-acyl carrier protein transacylase | 264.221 | 0.077 | | | | | | |
| *FABF* | 3-oxoacyl-[acyl-carrier-protein] synthase 2 | | | | | | | | |
| *FABG* | 3-oxoacyl-[acyl-carrier-protein] reductase | 1.604 | 2.157 | | | | | | |
| *FABG_1* | 3-oxoacyl-[acyl-carrier-protein] reductase | 1.602 | 2.162 | | | | | | |
| *FABH_1* | 3-oxoacyl-[acyl-carrier-protein] synthase 3 | | | | | 283 | 0.991 | 283 | 0.025 |
| *FBAB* | Fructose-bisphosphate aldolase class 1 | 13.141 | 1.218 | | | 9 | 0.973 | | |
| *FBPB* | Diacylglycerol acyltransferase/mycolyltransferase Ag85B | 1.087 | 11.252 | | | 57<br>64 | 0.982<br>0.990 | 57<br>64<br>138 | 0.031<br>0.031<br>0.022 |

| | | | | | | | | | | | |
|---|---|---|---|---|---|---|---|---|---|---|---|
| *FCL* | GDP-L-fucose synthase | 2.812 | 3.628 | 189 | 0.033 | 189<br>275<br>349 | 0.996<br>0.961<br>0.983 | 189<br>349 | 0.033<br>0.016 | | |
| *FFH* | Signal recognition particle protein | 4.651 | 0.886 | | | 337 | 0.990 | | | | |
| *FIEF* | Ferrous-iron efflux pump | 2.859 | 6.589 | 187<br>195 | 0.015<br>0.022 | 187<br>195 | 0.995<br>0.992 | 187<br>195 | 0.004<br>0.005 | | |
| *FIM1C_2* | hypothetical protein | 1.307 | 12.202 | | | 25 | 0.956 | 195 | 0.031 | | |
| *FIMB* | Type 1 fimbriae regulatory protein FimB | 283<br>411<br>455<br>493 | 0.016<br>0.046<br>0.046<br>0.008 | 82<br>83<br>88<br>207<br>283<br>374<br>411<br>455<br>462<br>493<br>528<br>540 | 0.973<br>0.981<br>0.977<br>0.960<br>0.994<br>0.961<br>0.992<br>0.992<br>0.969<br>0.998<br>0.990<br>0.991 | 283<br>493 | 0.025<br>0.000 | | | | |
| *FMT* | Methionyl-tRNA formyltransferase | 225.909 | 0.35 | | | | | | | | |
| *FOLD* | Bifunctional protein FolD protein | 84.125 | 0.324 | | | 27 | 0.981 | 27<br>67 | 0.031<br>0.040 | | |
| *FPGS* | Folylpolyglutamate synthase | | | 102<br>219<br>253 | 0.037<br>0.033<br>0.045 | 102<br>180<br>219<br>245<br>253 | 0.989<br>0.976<br>0.990<br>0.969<br>0.983 | 219 | 0.048 | | |
| *FPTA* | Fe(3+)-pyochelin receptor | 2.737 | 5.13 | 594 | 0.001 | 88<br>169<br>327<br>594 | 0.999<br>0.961<br>0.984<br>0.999 | 88<br>594 | 0.008<br>0.000 | 88<br>594 | 0.039<br>0.005 |
| *FTSX* | Cell division protein | 29.671 | 0.5 | | | | | 271 | 0.028 | | |

| | | | | | | | | | | | |
|---|---|---|---|---|---|---|---|---|---|---|---|
| *FUMB* | Fumarate hydratase class I, anaerobic | 1.778 | 5.947 | 362<br>364 | 0.005<br>0.020 | 362<br>364 | 0.999<br>0.994 | 362<br>364 | 0.009<br>0.011 | 362<br>364 | 0.049<br>0.039 |
| *FYUA_1* | Pesticin receptor | | | | | 16<br>282<br>429<br>712 | 0.992<br>0.972<br>0.958<br>0.992 | 371 | 0.028 | 16 | 0.047 |
| *GADC* | Glutamate/gamma-aminobutyrate antiporter | 1.544 | 5.943 | | | 495 | 0.962 | 505 | 0.039 | | |
| *GCDB_1* | Glutaconyl-CoA decarboxylase subunit beta | 8.53 | 1.104 | 405 | 0.000 | 405 | 0.999 | 405 | 0.000 | 405 | 0.001 |
| *GCVT* | Aminomethyltransferase | 9.88 | 2.75 | | | 42 | 0.969 | 357 | 0.042 | | |
| *GLAB_1* | Alpha-1,3-galactosidase B | | | 224<br>436 | 0.028<br>0.027 | 137<br>224<br>296<br>390<br>436 | 0.991<br>0.992<br>0.984<br>0.957<br>0.992 | 224<br>436 | 0.041<br>0.040 | | |
| *GLAB_2* | Alpha-1,3-galactosidase B | | | | | 9<br>75<br>159<br>188<br>229<br>429<br>549 | 0.984<br>0.973<br>0.959<br>0.963<br>0.972<br>0.957<br>0.981 | | | | |
| *GLGB_1* | 1,4-alpha-glucan branching enzyme | 1.828 | 4.045 | 243 | 0.030 | 243 | 0.999 | 74<br>107<br>243 | 0.023<br>0.049<br>0.043 | | |
| *GLNA_1* | Glutamine synthetase | 9.122 | 1.134 | 282 | 0 | 20<br>282 | 0.962<br>0.999 | 276<br>280<br>282 | 0.035<br>0.023 | 282 | 0.000 |

| | | | | | | | | | |
|---|---|---|---|---|---|---|---|---|---|
| *GLPC* | Anaerobic glycerol-3-phosphate dehydrogenase subunit C | | | 602<br>704<br>786 | 0.031<br>0.043<br>0.013 | 217<br>444<br>590<br>602<br>630<br>704<br>786<br>948 | 0.991<br>0.967<br>0.986<br>0.992<br>0.973<br>0.997<br>0.998<br>0.979 | 80<br>602<br>786<br>943 | 0.050<br>0.046<br>0.021<br>0.001 |
| *GNO* | Gluconate 5-dehydrogenase | | | | | 203 | 0.975 | | |
| *GROUP_11981* | Aminopeptidase S | 1.847 | 4.428 | | | 265 | 0.980 | 265 | 0.012 |
| *GROUP_14692* | Macrolide export ATP-binding/permease protein | | | 10 | 0.047 | 10 | 0.986 | | |
| *GROUP_14757* | Penicillin-binding protein 1C | | | | | 8<br>476 | 0.969<br>0.971 | | |
| *GROUP_15240* | N-carbamoyl-D-amino acid hydrolase | 1.921 | 6.699 | | | 14 | 0.975 | 4<br>37<br>224 | 0.039<br>0.048<br>0.024 |
| *GROUP_15605* | Keratan-sulfate endo-1,4-beta-galactosidase | 4.079 | 5.496 | 85<br>165 | 0.020<br>0.012 | 85<br>165 | 0.997<br>0.998 | 85<br>165 | 0.011<br>0.012 |
| *GROUP_16529* | Beta-galactosidase BoGH2A | 1 | 4.945 | 762 | 0.043 | 762 | 0.994 | 210<br>318<br>762 | 0.022<br>0.033<br>0.006 |
| *GROUP_21132* | Glycosyl hydrolase family 109 protein 1 | | | | | 20 | 0.984 | 20 | 0.040 |
| *GROUP_29929* | Lysozyme M1 | | | | | 33 | 0.966 | | |
| *GROUP_30172* | Murein hydrolase activator | | | | | 218 | 0.953 | | |
| *GROUP_3* | Esterase EstB | 1 | 14.560 | | | 528 | 0.984 | | |

| | | | | | | | | | | | |
|---|---|---|---|---|---|---|---|---|---|---|---|
| *0185* | | | | | | | | | | | |
| *GUAB_1* | Inosine-5'-monophosphate dehydrogenase | 7.159 | 1.656 | 340 | 0.042 | 340 | 0.9883 | 121<br>129<br>311<br>340 | 0.042<br>0.036<br>0.016<br>0.001 | | |
| *GUAD* | Guanine deaminase | | | | | 39 | 0.986 | 39 | 0.030 | | |
| *GYRA* | DNA gyrase subunit B | 140.626 | 0.385 | | | 82<br>418 | 0.990<br>0.955 | 418 | 0.038 | | |
| *HDC* | Histidine decarboxylase proenzyme | 2.449 | 5.123 | 258 | 0.006 | 249<br>258 | 0.963<br>0.999 | 258 | 0.000 | 258 | 0.017 |
| *HEMA* | Glutamyl-tRNA reductase | 1.538 | 10.41 | 112 | 0.019 | 83<br>112 | 0.977<br>0.990 | 12<br>112 | 0.046<br>0.012 | | |
| *HEME* | Uroporphyrinogen decarboxylase | | | | | 5 | 0.954 | | | | |
| *HISD* | Histidinol dehydrogenase | 7.98 | 2.269 | 95 | 0.026 | 95 | 0.983 | 20<br>95 | 0.007<br>0.004 | | |
| *HEME_1* | Uroporphyrinogen decarboxylase | | | | | 17<br>35<br>125 | 0.962<br>0.959<br>0.989 | 125 | 0.047 | | |
| *HLYD* | Hemolysin secretion protein D, chromosomal | 44.83 | 0.761 | | | 135 | 0.955 | | | | |
| *HTPG_1* | Chaperone protein HtpG | 198.793 | 1.637 | 122 | 0.006 | 122 | 0.986 | 122 | 0.002 | 122 | 0.005 |
| *HTPG_2* | Chaperone protein HtpG | 2.815 | 3.663 | | | | | 399<br>519 | 0.045<br>0.025 | | |
| *HYPBA1_3* | Non-reducing end beta-L-arabinofuranosidase | | | | | 337<br>481<br>551 | 0.964<br>0.969<br>0.981 | | | | |
| *ICD* | Isocitrate dehydrogenase [NADP] | 2.483 | 7.811 | 349 | 0.008 | 51<br>349 | 0.956<br>0.994 | 349 | 0.002 | | |

| | | | | | | | | | | | |
|---|---|---|---|---|---|---|---|---|---|---|---|
| *IHFA_1* | Integration host factor subunit alpha | | | 218 | 0.043 | 218<br>265<br>370<br>411<br>461 | 0.989<br>0.963<br>0.962<br>0.970<br>0.956 | | | | |
| *ILVD* | Dihydroxy-acid dehydratase | 5.694 | 1.036 | | | 292<br>403 | 0.969<br>0.987 | 235<br>292 | 0.009<br>0.048 | | |
| *KATA* | Catalase | 1.002 | 3.861 | | | 474 | 0.952 | | | | |
| *KBL* | 2-amino-3-ketobutyrate coenzyme A ligase | 6.003 | 1.172 | | | 158 | 0.969 | | | | |
| *KDPA* | Potassium-transporting ATPase potassium-binding subunit | 2.277 | 5.516 | 313<br>315 | 0.018<br>0.051 | 313<br>315 | 0.994<br>0.963 | 315<br>392 | 0.009<br>0.034 | | |
| *LOIP* | Metalloprotease | 7.669 | 2.920 | 52 | 0.043 | 52 | 0.981 | | | | |
| *LON* | Lon protease | 1.002 | 1.994 | | | 820 | 0.969 | 242 | 0.024 | | |
| *LPTB* | Lipopolysaccharide export system ATP-binding protein | 7.733 | 0.463 | | | 2 | 0.981 | 2 | 0.028 | | |
| *LPTD_1* | LPS-assembly protein | | | 759 | 0.0074 | 759 | 0.998 | 759 | 0.012 | 759 | 0.039 |
| *LDHA* | D-lactate dehydrogenase | 117.034 | 0.35 | | | 91 | 0.960 | 29<br>68<br>326 | 0.008<br>0.011<br>0.016 | | |
| *LEPA* | Elongation factor 4 | 4.926 | 0.998 | | | 215 | 0.957 | 215 | 0.024 | | |
| *LIPA* | Lipoyl synthase | 1.723 | 8.743 | 88<br>150 | 0.988<br>0.954 | 74<br>150 | 0.050<br>0.012 | | | | |
| *LYSJ* | [LysW]-aminoadipate semialdehyde transaminase | 4.297 | 6.435 | 332 | 0.024 | 306<br>332 | 0.976<br>0.996 | 169<br>332 | 0.032<br>0.004 | | |
| *LYSU* | Lysine--tRNA ligase, heat inducible | | | 518 | 0.041 | 403<br>518 | 0.952<br>0.992 | 518 | 0.004 | | |
| *LPXC* | UDP-3-O-acyl-N-acetylglucosamine deacetylase | | | 249 | 0.005 | 58<br>249 | 0.984<br>0.995 | 249 | 0.001 | 249 | 0.012 |
| *LPXD* | UDP-3-O-acylglucosamine N-acyltransferase | | | | | 103 | 0.953 | | | | |
| *LMRA* | Putative multidrug export ATP-binding/permease protein | | | 450 | 0.004 | 450 | 0.997 | 450 | 0.007 | | |

| | | | | | | | | | |
|---|---|---|---|---|---|---|---|---|---|
| *INFB* | Translation initiation factor IF-2 | 1.43 | 3.064 | | | 126 | 0.964 | 126<br>128<br>179 | 0.044<br>0.034<br>0.044 |
| *IOLX* | Scyllo-inositol 2-dehydrogenase (NAD(+)) | 7.345 | 1.603 | | | 52 | 0.988 | 52<br>54 | 0.012<br>0.037 |
| *LPXL* | Lipid A biosynthesis lauroyltransferase | | | 258 | 0.044 | 258 | 0.978 | 258 | 0.017 |
| *LTXD* | Leukotoxin export protein | | | | | 222 | 0.959 | | |
| *MACA_1* | Macrolide export protein | 1 | 5.339 | | | 365 | 0.980 | 38 | 0.005 |
| *MACB_3* | Macrolide export ATP-binding/permease protein | | | | | 154 | 0.968 | | |
| *MACB_4* | Macrolide export ATP-binding/permease protein | | | 340 | 0.021 | 340<br>342<br>363 | 0.995<br>0.960<br>0.978 | | |
| *MACB_5* | Macrolide export ATP-binding/permease protein | | | 368 | 0.038 | 103<br>271<br>295<br>368 | 0.961<br>0.988<br>0.977<br>0.989 | 271<br>368 | 0.072<br>0.038 |
| *MACB_6* | Macrolide export ATP-binding/permease protein | 146.998 | 0.397 | 233 | 0.044 | 233 | 0.991 | 117<br>233 | 0.045<br>0.012 |
| *MACB_7* | Macrolide export ATP-binding/permease protein | 1.276 | 8.472 | | | | | 354 | 0.028 |
| *MACB_8* | Macrolide export ATP-binding/permease protein | 14.538 | 0.692 | 181 | 0.019 | 181 | 0.997 | 181 | 0.030 |
| *MACB_10* | Macrolide export ATP-binding/permease protein | | | 219 | 0.039 | 219<br>234 | 0.990<br>0.966 | | |
| *MACB_11* | Macrolide export ATP-binding/permease protein | | | 513 | 0.033 | 470<br>513<br>585 | 0.976<br>0.991<br>0.980 | 513 | 0.048 |

| | | | | | | | | | | | |
|---|---|---|---|---|---|---|---|---|---|---|---|
| *MACB_12* | Macrolide export ATP-binding/permease protein | | | 383<br>620 | 0.045<br>0.017 | 115<br>117<br>222<br>234<br>236<br>315<br>354<br>383<br>620<br>659 | 0.984<br>0.984<br>0.981<br>0.984<br>0.971<br>0.988<br>0.977<br>0.988<br>0.994<br>0.986 | 620 | 0.026 | | |
| *MACB_13* | Macrolide export ATP-binding/permease protein | | | 16<br>42<br>618<br>779 | 0.021<br>0.026<br>0.033<br>0.022 | 16<br>42<br>101<br>618<br>640<br>779 | 0.997<br>0.996<br>0.975<br>0.996<br>0.976<br>0.995 | 16<br>42<br>618<br>779 | 0.032<br>0.039<br>0.047<br>0.033 | | |
| *MACB_14* | Macrolide export ATP-binding/permease protein | 143.103 | 0.898 | | | 3<br>153<br>159<br>712 | 0.954<br>0.959<br>0.984<br>0.978 | 3 | 0.014 | | |
| *MALQ* | 4-alpha-glucanotransferase | 3.329 | 7.107 | 370<br>496<br>858 | 0.013<br>0.003<br>0.002 | 83<br>370<br>496<br>821<br>858 | 0.966<br>0.995<br>0.999<br>0.978<br>0.999 | 82<br>370<br>496<br>576<br>821<br>858 | 0.041<br>0.004<br>0.006<br>0.045<br>0.033<br>0.004 | 496<br>858 | 0.019<br>0.009 |

| | | | | | | | | | | | |
|---|---|---|---|---|---|---|---|---|---|---|---|
| *MAUG* | Methylamine utilization protein | | | 118<br>150<br>201<br>255<br>321 | 0.037<br>0.048<br>0.026<br>0.046<br>0.043 | 4<br>118<br>150<br>201<br>255<br>262<br>321 | 0.976<br>0.991<br>0.991<br>0.993<br>0.978<br>0.971<br>0.986 | 4<br>118<br>201<br>262<br>321 | 0.042<br>0.013<br>0.038<br>0.010<br>0.014 | | |
| *MDTA_1* | Multidrug resistance protein | 41.944 | 0.299 | | | 97 | 0.031 | | | | |
| *MDTC_2* | Multidrug resistance protein | | | 250<br>788<br>825 | 0.007<br>0.019<br>0.009 | 204<br>250<br>743<br>788<br>825<br>885 | 0.975<br>0.999<br>0.975<br>0.995<br>0.998<br>0.975 | 250<br>514<br>788<br>825 | 0.012<br>0.049<br>0.029<br>0.015 | 250<br>788<br>825 | 0.047<br>0.039<br>0.045 |
| *MDTE_2* | Multidrug resistance protein | | | | | 341 | 0.963 | | | | |
| *MDTE_3* | Multidrug resistance protein | 50.21 | 0.959 | 318 | 0.042 | | | 2<br>9<br>127<br>130<br>167<br>269<br>318<br>356 | 0.048<br>0.037<br>0.034<br>0.071<br>0.031<br>0.074<br>0.004<br>0.014 | | |
| *MDTE_4* | Multidrug resistance protein | 1.203 | 3.461 | | | | | | | | |
| *MDTK_1* | Multidrug resistance protein | 1.201 | 7.767 | | | 121 | 0.960 | 249 | 0.040 | | |
| *MDTN* | Hypothetical protein | 138.574 | 0.661 | | | 226<br>242 | 0.9842<br>0.972 | | | | |
| *MECI* | Methicillin resistance regulatory protein | 1.709 | 7.006 | | | | | | | | |

| | | | | | | | | | | | |
|---|---|---|---|---|---|---|---|---|---|---|---|
| *MENE* | 2-succinylbenzoate--CoA ligase | 108.978 | 2.266 | 158<br>204 | 0.044<br>0.012 | 153<br>158<br>204<br>252<br>364 | 0.979<br>0.991<br>0.996<br>0.975<br>0.979 | 204 | 0.019 | 204 | 0.034 |
| *MEPA_2* | Multidrug export protein | 11.628 | 2.201 | 41 | 0.002 | 41 | 0.998 | 41<br>404 | 0.001<br>0.048 | 41 | 0.011 |
| *MEPA_5* | Multidrug export protein | 4.176 | 1.845 | | | 435<br>459 | 0.957<br>0.993 | | | | |
| *METH_1* | Methionine synthase | 37.306 | 0.963 | 84 | 0.021 | 84 | 0.995 | 29<br>84 | 0.045<br>0.005 | | |
| *METH_4* | Methionine synthase | 1.228 | 5.9 | 133<br>323 | 0.0453<br>0.021 | 133<br>265<br>323 | 0.979<br>0.967<br>0.990 | 323<br>654<br>902 | 0.0001<br>0.045<br>0.022 | | |
| *METK* | S-adenosylmethionine synthase | 10.136 | 1.344 | | | 200 | 0.984 | 331 | 0.022 | | |
| *MEXA* | Multidrug resistance protein | 2.142 | 12.049 | 150 | 0.050 | 20<br>76<br>150 | 0.973<br>0.966<br>0.989 | 33<br>339 | 0.042<br>0.017 | | |
| *MEXB* | Multidrug resistance protein | 3.116 | 2.201 | 271 | 0.042 | 30<br>31<br>271<br>937 | 0.951<br>0.964<br>0.992<br>0.971 | 18<br>271<br>647<br>870 | 0.045<br>0.003<br>0.006<br>0.002 | | |
| *MFA2_1* | Minor fimbrium anchoring subunit | 1.082 | 0 | 102<br>129 | 0.005<br>0.041 | 102<br>129 | 0.997<br>0.981 | 102<br>129 | 0.008<br>0.047 | 102 | 0.008 |
| *MFD* | Transcription-repair-coupling factor | 2.427 | 3.036 | 265 | 0.042 | 265<br>371 | 0.990<br>0.985 | 371<br>806 | 0.029<br>0.034 | | |
| *MGGB* | Mannosylglucosyl-3-phosphoglycerate phosphatase | 1.711 | 5.7 | 46 | 0.002 | 46 | 0.999 | 46<br>105<br>461<br>462 | 0.000<br>0.041<br>0.028<br>0.020 | 46 | 0.017 |

| | | | | | | | | | | | |
|---|---|---|---|---|---|---|---|---|---|---|---|
| *MIAA* | tRNA dimethylallyltransferase | 9.127 | 5.31 | 216 | 0.049 | 3<br>216 | 0.958<br>0.992 | 3<br>201 | 0.017<br>0.031 | | |
| *MIAA_2* | tRNA dimethylallyltransferase | 31.193 | 1.943 | | | 287 | 0.957 | 248 | 0.049 | | |
| *MIP_2* | Outer membrane protein | 10.335 | 1.741 | | | 366 | 0.988 | 366 | 0.020 | | |
| *MLEN* | Malate-2H(+)/Na(+)-lactate antiporter | 3.577 | 2.665 | 66 | 0.034 | 66 | 0.990 | 66<br>268 | 0.010<br>0.037 | | |
| *MMGC* | Acyl-CoA dehydrogenase | 9.073 | 0.493 | | | 366 | 0.988 | 366 | 0.019 | | |
| *MNMA_3* | tRNA-specific 2-thiouridylase | 4.05 | 5.868 | 171<br>268<br>339 | 0.037<br>0.018<br>0.024 | 69<br>171<br>268<br>287<br>313<br>339 | 0.988<br>0.992<br>0.995<br>0.979<br>0.981<br>0.992 | 69<br>171<br>268<br>313<br>339 | 0.028<br>0.011<br>0.028<br>0.019<br>0.001 | | |
| *MNMG* | tRNA uridine 5-carboxymethylaminomethyl modification enzyme | 1.279 | 7.507 | 46 | 0.002 | 46 | 0.999 | 46<br>105<br>461<br>462 | 0.000<br>0.041<br>0.028<br>0.020 | 46 | 0.017 |
| *MRCA* | Penicillin-binding protein 1A | 5.521 | 1.679 | 286<br>660<br>746 | 0.001<br>0.011<br>0.030 | 286<br>660<br>746 | 0.999<br>0.995<br>0.993 | 286<br>398<br>637<br>660<br>710<br>712<br>718<br>746 | 0.001<br>0.007<br>0.015<br>0.000<br>0.000<br>0.027<br>0.024<br>0.007 | 286 | 0.008 |
| *MRDA* | Peptidoglycan D,D-transpeptidase | 6.144 | 1.051 | | | 147 | 0.974 | | | | |
| *MSCM* | Miniconductance mechanosensitive channel | | | 573 | 0.0200 | 573 | 0.993 | 107<br>502<br>573 | 0.038<br>0.042<br>0.008 | | |
| *MTAB* | Threonylcarbamoyladenosine tRNA methylthiotransferase | 4.455 | 3.974 | 73 | 0.0018 | 73 | 0.9991 | 73<br>395 | 0.000<br>0.036 | 73 | 0.017 |

| | | | | | | | | | | | |
|---|---|---|---|---|---|---|---|---|---|---|---|
| *MUTS_3* | DNA mismatch repair protein | 10.517 | 0.503 | | | 612 | 0.978 | 340 | 0.043 | | |
| *NAGB_2* | Glucosamine-6-phosphate deaminase 1 | 9.33 | 0.362 | | | | | 255 | 0.027 | | |
| *NANM_2* | N-acetylneuraminate epimerase | 5.572 | 4.445 | 21<br>208 | 0.0473<br>0.0415 | 25<br>208<br>321 | 0.9963<br>0.9991<br>0.9772 | 21<br>25<br>208<br>321 | 0.013<br>0.007<br>0.013<br>0.009 | | |
| *NEDA_2* | Sialidase | | | | | 119 | 0.959 | 119 | 0.044 | | |
| *NEUA* | N-acylneuraminate cytidylyltransferase | | | 38<br>47<br>66<br>122 | 0.004<br>0.033<br>0.046<br>0.041 | 24<br>38<br>47<br>66<br>69<br>93<br>122 | 0.954<br>0.999<br>0.989<br>0.984<br>0.955<br>0.956<br>0.9836 | 38<br>47 | 0.007<br>0.048 | | |
| *NFO* | Endonuclease 4 | 42.325 | 1.631 | 214 | 0.003 | 214 | 0.999 | 214 | 0 | 214 | 0.026 |
| *NHAA* | Na(+)/H(+) antiporter NhaA | 1.43 | 9.108 | 337<br>353 | 0.000<br>0.008 | 337<br>353<br>435 | 1.000<br>0.999<br>0.978 | 222<br>337<br>353 | 0.044<br>0.000<br>0.003 | 337<br>353 | 0.000<br>0.026 |
| *NLPE* | Lipoprotein | 10.867 | 3.902 | 27<br>39 | 0.962<br>0.956 | 27 | 0.050 | | | | |
| *NNR* | Bifunctional NAD(P)H-hydrate repair enzyme | 1.852 | 4.492 | 117 | 0.030 | 117 | 0.993 | 117 | 0.044 | | |
| *NORM_2* | Multidrug resistance | 31.56 | 0.693 | | | | | 17 | 0.026 | | |
| *NQO6* | NADH-quinone oxidoreductase subunit 6 | | | | | 89 | 0.9684 | | | | |
| *NRDD* | Anaerobic ribonucleoside-triphosphate reductase | 10.173 | 0.969 | 743 | 0.0005 | 608<br>743 | 0.985<br>0.999 | 543<br>743<br>797 | 0.005<br>0.000<br>0.007 | 743 | 0.002 |
| *NUDC_2* | NADH pyrophosphatase | 4.614 | 4.108 | | | 151 | 0.9566 | | | | |
| *NUOL* | NADH-quinone oxidoreductase subunit L | 2.098 | 3.844 | 463 | 0.005 | 463 | 0.999 | 166<br>307<br>463<br>525 | 0.013<br>0.011<br>0.001<br>0.042 | | |

| | | | | | | | | | | | |
|---|---|---|---|---|---|---|---|---|---|---|---|
| *NUON* | NADH-quinone oxidoreductase subunit N | | | 28<br>455 | 0.008<br>0.002 | 28<br>37<br>42<br>65<br>455 | 0.998<br>0.981<br>0.958<br>0.958<br>0.999 | 28<br>65<br>455 | 0.014<br>0.020<br>0.003 | 28<br>455 | 0.050<br>0.008 |
| *OPRM_1* | Outer membrane protein | 2.484 | 5.441 | 107 | 0.007 | 107<br>297 | 0.997<br>0.966 | 107<br>297 | 0.000<br>0.011 | | |
| *OPRM_2* | Outer membrane protein | 9.084 | 1.27 | | | 16 | 0.9596 | | | | |
| *OXYR* | Hydrogen peroxide-inducible genes activator | 1.533 | 4.233 | | | | | 59 | 0.042 | | |
| *PAAK* | Phenylacetate-coenzyme A ligase | 31.510 | 0.842 | | | 44<br>312 | 0.971<br>0.970 | | | | |
| *PABA* | Aminodeoxychorismate synthase component 2 | | | | | 38 | 0.958 | | | | |
| *PANC* | Pantothenate synthetase | 2.748 | 7.012 | | | 171 | 0.964 | 104<br>171 | 0.009<br>0.032 | | |
| *PABB* | Aminodeoxychorismate synthase component 1 | | | | | 62<br>64 | 0.970<br>0.963 | | | | |
| *PATA_2* | Peptidoglycan O-acetyltransferase | 3.324 | 4.136 | 83<br>435 | 0.047<br>0.000 | 83<br>435 | 0.984<br>0.999 | 47<br>83<br>435 | 0.024<br>0.006<br>0.000 | 435 | 0.005 |
| *PBPC* | Penicillin-binding protein 1C | 13.678 | 2.188 | 435<br>486 | 0.0055<br>0.0054 | 19<br>55<br>435<br>486 | 0.953<br>0.960<br>0.999<br>0.999 | 435<br>486 | 0.009<br>0.009 | 435<br>486 | 0.026<br>0.040 |
| *PCM* | Protein-L-isoaspartate O-methyltransferase | | | | | 240<br>337 | 0.953<br>0.963 | | | | |
| *PCRA* | ATP-dependent DNA helicase | 12.638 | 1.373 | | | 703 | 0.980 | 2<br>296<br>449 | 0.039<br>0.024<br>0.034 | | |
| *PEPD_2* | Dipeptidase | 6.159 | 1.280 | | | | | 454<br>462 | 0.009<br>0.061 | | |
| *PEPD_3* | Cytosol non-specific dipeptidase | | | | | 359 | 0.961 | | | | |

| | | | | | | | | | | | |
|---|---|---|---|---|---|---|---|---|---|---|---|
| *PEPE_2* | Aminopeptidase E | 3.594 | 1.671 | | | 19 | 0.978 | | | | |
| *PEPN* | Aminopeptidase N | | | 124<br>262<br>616<br>628 | 0.022<br>0.031<br>0.049<br>0.034 | 62<br>124<br>262<br>421<br>437<br>453<br>494<br>616<br>628<br>800 | 0.999<br>0.996<br>0.995<br>0.974<br>0.986<br>0.989<br>0.989<br>0.988<br>0.995<br>0.989 | 62<br>124<br>262<br>628 | 0.001<br>0.033<br>0.020<br>0.049 | 62 | 0.012 |
| *PEPO* | Neutral endopeptidase | 1.397 | 8.811 | 551 | 0.009 | 446<br>551 | 0.965<br>0.996 | 216<br>233<br>446<br>551 | 0.042<br>0.04<br>0.025<br>0.000 | 551 | 0.039 |
| *PEPT* | Peptidase T | 7.458 | 3.216 | 61<br>106 | 0.000<br>0.011 | 61<br>106 | 0.999<br>0.995 | 61<br>69<br>106 | 0.000<br>0.029<br>0.003 | 61 | 0.000 |
| *PER1* | Extended-spectrum beta-lactamase PER-1 | 16.857 | 0.62 | 181 | 0.019 | 181 | 0.997 | 181 | 0.030 | | |
| *PGCA* | Phosphoglucomutase | 6.297 | 1.561 | 201<br>520 | 0.973<br>0.960 | 520 | 0.0030 | | | | |
| *PGK* | Phosphoglycerate kinase | 14.017 | 2.038 | 271 | 0.016 | 271<br>272 | 0.999<br>0.972 | 271<br>272 | 0.001<br>0.001 | 271 | 0.039 |
| *PGPH* | Cyclic-di-AMP phosphodiesterase | 1.667 | 3.981 | | | 194 | 0.984 | 146 | 0.009 | | |
| *PHAB* | Acetoacetyl-CoA reductase | | | 28 | 0.011 | 28 | 0.997 | 28 | 0.012 | 28 | 0.037 |
| *PHOB* | Alkaline phosphatase 3 | 19.105 | 0.555 | | | | | | | | |
| *PHOR_1* | Phosphate regulon sensor protein | 1.187 | 0 | 534 | 0.049 | 534<br>575<br>871<br>1154<br>1418 | 0.987<br>0.972<br>0.985<br>0.974<br>0.956 | | | | |

| | | | | | | | | | | | |
|---|---|---|---|---|---|---|---|---|---|---|---|
| *PHOR_3* | Alkaline phosphatase synthesis sensor protein | | | | | 528<br>540<br>542 | 0.951<br>0.967<br>0.982 | | | | |
| *PINR* | Serine recombinase | | | | | 159 | 0.960 | | | | |
| *PNCC* | Nicotinamide-nucleotide amidohydrolase | | | | | 23<br>177 | 0.955<br>0.965 | | | | |
| *PNP* | Polyribonucleotide nucleotidyltransferase | 49.993 | 0.151 | | | | | | | | |
| *POLA* | DNA polymerase I | 1.001 | 8.763 | 324 | 0.0404 | 324<br>410<br>726 | 0.994<br>0.950<br>0.974 | 410 | 0.004 | | |
| *POTA* | Spermidine/putrescine import ATP-binding protein | 45.888 | 0.401 | | | 380 | 0.963 | | | | |
| *POTD* | Spermidine/putrescine-binding periplasmic protein | 133.775 | 0.451 | | | 167 | 0.981 | 263 | 0.023 | | |
| *PSAA* | Manganese ABC transporter substrate-binding lipoprotein | | | | | 38 | 0.958 | | | | |
| *PSAA_2* | Manganese ABC transporter substrate-binding lipoprotein | | | | | 175 | 0.959 | | | | |
| *PURC* | Phosphoribosylaminoimidazole-succinocarboxamide synthase | 2.077 | 6.238 | 218 | 0.022 | 218 | 0.994 | 218<br>298 | 0.008<br>0.048 | 218 | 0.027 |
| *PURF_1* | Amidophosphoribosyltransferase | | | | | 139 | 0.950 | | | | |
| *PURF_2* | Amidophosphoribosyltransferase | 62.222 | 0.284 | 60 | 0.023 | 60<br>99 | 0.979<br>0.984 | 60<br>148 | 0.002<br>0.011 | | |
| *PURM* | Phosphoribosylformylglycinamidine cyclo-ligase | 30.686 | 0.644 | 354<br>357 | 0.953<br>0.953 | 357 | 0.041 | | | | |
| *PUTP* | High-affinity proline transporter | 6.782 | 1.957 | | | | | 21<br>476<br>499 | 0.027<br>0.031<br>0.019 | | |
| *PYRC* | Dihydroorotase | 21.571 | 1.048 | 18<br>48 | 0.012<br>0.044 | 18<br>48 | 0.994<br>0.995 | 18<br>71<br>317 | 0.020<br>0.038<br>0.045 | | |

| | | | | | | | | | |
|---|---|---|---|---|---|---|---|---|---|
| *PYRD_1* | Dihydroorotate dehydrogenase (quinone) | | | 37<br>188<br>257 | 0.040<br>0.039<br>0.048 | 37<br>42<br>228<br>257 | 0.992<br>0.968<br>0.983<br>0.990 | | |
| *PYRD_2* | Dihydroorotate dehydrogenase B (NAD(+)), catalytic subunit | 3.206 | 3.179 | | | 274 | 0.978 | 274 | 0.018 |
| *PYRF* | Orotidine 5'-phosphate decarboxylase | 6.931 | 2.555 | | | 161 | 0.976 | 161 | 0.037 |
| *PYRK* | Dihydroorotate dehydrogenase B (NAD(+)), electron transfer subunit | | | | | 33 | 0.957 | | |
| *QUEA_2* | S-adenosylmethionine:tRNA ribosyltransferase-isomerase | 33.059 | 0.957 | 273 | 0.039 | 273 | 0.993 | 273 | 0.012 |
| *QUEE_1* | 7-carboxy-7-deazaguanine synthase | 5.652 | 2.877 | | | 43 | 0.970 | | |
| *RBR* | Rubrerythrin | 8.692 | 1.569 | | | 253<br>259 | 0.994<br>0.992 | 253<br>259 | 0.019<br>0.006 |
| *RECD2_1* | ATP-dependent RecD-like DNA helicase | 5.926 | 3.103 | | | 158 | 0.955 | 158 | 0.019 |
| *RECG* | ATP-dependent DNA helicase | 2.369 | 4.218 | 183<br>400 | 0.034<br>0.009 | 183<br>365<br>400 | 0.981<br>0.976<br>0.999 | 183<br>365<br>400<br>646 | 0.045<br>0.014<br>0.000<br>0.035 |
| *RECJ* | Single-stranded-DNA-specific exonuclease | 1.943 | 1.739 | 175 | 0.033 | 175 | 0.990 | 175 | 0.015 |
| *RECN* | DNA repair protein RecN | 5.836 | 4.408 | 169<br>250<br>265 | 0.044<br>0.051<br>0.006 | 169<br>250<br>265 | 0.990<br>0.973<br>0.997 | 169<br>250<br>265 | 0.019<br>0.006<br>0.008 |
| *RESA_3* | Thiol-disulfide oxidoreductase | 4.502 | 0.000 | 24<br>33 | 0.009<br>0.022 | 24<br>33 | 0.998<br>0.997 | 24<br>33 | 0.004<br>0.034 |
| *RFBA* | Glucose-1-phosphate thymidylyltransferase 2 | 18.797 | 1.5 | 38<br>98<br>212 | 0.973<br>0.980<br>0.966 | 98 | 0.034 | | |
| *RFBB* | dTDP-glucose 4,6-dehydratase | 8.692 | 1.569 | | | 253<br>259 | 0.994<br>0.992 | 253<br>259 | 0.019<br>0.006 |

| | | | | | | | | | | | |
|---|---|---|---|---|---|---|---|---|---|---|---|
| *RFFH_1* | Glucose-1-phosphate thymidylyltransferase 2 | 4.252 | 0.774 | | | | | 234<br>266 | 0.018<br>0.006 | | |
| *RFFH_2* | Glucose-1-phosphate thymidylyltransferase 2 | 2.074 | 3.903 | | | 76 | 0.031 | 76 | 0.031 | | |
| *RFFH_3* | Glucose-1-phosphate thymidylyltransferase 2 | 4.034 | 3.25 | 12<br>78 | 0.016<br>0.020 | 12<br>78 | 0.992<br>0.995 | 12<br>78 | 0.004<br>0.030 | | |
| *RFFH_4* | Glucose-1-phosphate thymidylyltransferase 2 | 50.867 | 0.327 | | | | | 66<br>95<br>99 | 0.033<br>0.044<br>0.014 | | |
| *RHO* | Transcription termination factor | | | 640 | 0.009 | 182<br>640 | 0.964<br>0.989 | 297<br>640 | 0.043<br>0.000 | 640 | 0.024 |
| *RIBBA* | Riboflavin biosynthesis protein | 1.463 | 3.036 | | | 120 | 0.983 | 120 | 0.026 | | |
| *RIBF* | Riboflavin biosynthesis protein | 251.64 | 1.353 | | | 81 | 0.9609 | | | | |
| *RLMB* | Putative TrmH family tRNA/rRNA methyltransferase | 136.542 | 0.443 | | | 148 | 0.971 | 148 | 0.024 | | |
| *RNC* | Ribonuclease 3 | | | | | 276 | 0.952 | | | | |
| *RPOC* | DNA-directed RNA polymerase subunit beta' | 4.958 | 0.456 | | | 672 | 0.992 | 265<br>672 | 0.035<br>0.044 | | |
| *SACC_1* | Levanase | 36.499 | 0.761 | 221<br>297<br>539 | 0.036<br>0.049<br>0.027 | 221<br>297<br>539 | 0.992<br>0.987<br>0.978 | 19<br>487<br>539 | 0.043<br>0.048<br>0.005 | | |
| *SCPA* | Methylmalonyl-CoA mutase | | | | | 253 | 0.951 | 253<br>313 | 0.045<br>0.044 | | |
| *SDCS_1* | Sodium-dependent dicarboxylate transporter | 4.824 | 2.307 | | | 438<br>455 | 0.955<br>0.969 | 163<br>352<br>438 | 0.034<br>0.036<br>0.028 | | |
| *SDCS_2* | Sodium-dependent dicarboxylate transporter | 3.324 | 4.218 | 131 | 0.000 | 130<br>131 | 0.956<br>0.999 | 131 | 0.000 | 131 | 0.000 |

| | | | | | | | | | | | |
|---|---|---|---|---|---|---|---|---|---|---|---|
| *SENX3* | Signal-transduction histidine kinase senX3 | 1.029 | 26.813 | 214 | 0.026 | 84<br>120<br>177<br>195<br>214<br>261 | 0.956<br>0.963<br>0.991<br>0.989<br>0.993<br>0.990 | 214 | 0.038 | | |
| *SERB* | Phosphoserine phosphatase | 6.134 | 2.531 | 118 | 0.0013 | 118 | 0.9934 | 118<br>168 | 0.0027<br>0.0183 | 118 | 0.039 |
| *RLMN* | Dual-specificity RNA methyltransferase | 7.111 | 4.128 | | | 303 | 0.973 | 303 | 0.023 | | |
| *RMPM* | Outer membrane protein class 4 | 17.944 | 0.938 | 572 | 0.0467 | 157<br>572 | 0.953<br>0.988 | 596 | 0.031 | | |
| *SIGE* | ECF RNA polymerase sigma factor SigE- 1 | 1.155 | 8.83 | | | 142 | 0.9523 | | | | |
| *SIGW_6* | ECF RNA polymerase sigma factor SigW | 2.380 | 7.970 | | | 29 | 0.961 | 49 | 0.030 | | |
| *SPOIIIE* | DNA translocase | 1.476 | 4.746 | 99<br>262<br>325 | 0.990<br>0.956<br>0.956 | 99 | 0.039 | | | | |
| *SRPC* | putative chromate transport protein | 1.848 | 4.65 | | | | | | | | |
| *SUFS* | Cysteine desulfurase | 36.364 | 0.459 | | | 301 | 0.965 | 301 | 0.032 | | |
| *SURA_1* | Chaperone SurA | | | | | | | 596<br>609 | 0.012<br>0.025 | | |
| *SURA_2* | Chaperone SurA | 6.989 | 1.192 | 139 | 0.0289 | 139 | 0.994 | 119<br>139 | 0.027<br>0.001 | | |
| *SUSC* | TonB-dependent receptor | 1.844 | 4.652 | | | | | | | | |
| *SUSC_1* | TonB-dependent receptor | 6.284 | 1.093 | | | 45 | 0.9613 | 45<br>443 | 0.025<br>0.044 | | |
| *SUSC_2* | TonB-dependent receptor | 1.194 | 3.586 | | | | | 411<br>951 | 0.0065<br>0.0003 | | |
| *SUSC_3* | TonB-dependent receptor | 8.317 | 1.212 | 479 | 0.0039 | 479<br>978<br>1054 | 0.999<br>0.960<br>0.994 | 479<br>1054 | 0.000<br>0.028 | 479 | 0.026 |

| | | | | | | | | | | | |
|---|---|---|---|---|---|---|---|---|---|---|---|
| *SUSC_4* | TonB-dependent receptor | 1.208 | 3.996 | | | 8 | 0.954 | 8<br>15 | 0.015<br>0.003 | | |
| *SUSC_5* | TonB-dependent receptor | 3.743 | 3.797 | 244 | 0.0003 | 18<br>89<br>244 | 0.979<br>0.950<br>0.999 | 5<br>18<br>89<br>244 | 0.960<br>0.979<br>0.950<br>0.999 | 244 | 0.002 |
| *SUSC_6* | TonB-dependent receptor | | | 23<br>434<br>435<br>438 | 0.005<br>0.003<br>0.000<br>0.000 | 23<br>434<br>435<br>438<br>814<br>954<br>1085 | 0.999<br>0.999<br>0.999<br>0.999<br>0.9728<br>0.958<br>0.9830 | 23<br>434<br>435<br>438 | 0.008<br>0.003<br>0.000<br>0.000 | 23<br>434<br>435<br>438 | 0.039<br>0.026<br>0.000<br>0.000 |
| *SUSC_7* | TonB-dependent receptor | 4.778 | 4.413 | 579<br>585<br>1006 | 0.017<br>0.009<br>0.027 | 83<br>155<br>579<br>585<br>1006<br>1050 | 0.976<br>0.993<br>0.991<br>0.999<br>0.986<br>0.990 | 155<br>276<br>579<br>585<br>852<br>1006<br>1050 | 0.034<br>0.041<br>0.031<br>0.000<br>0.013<br>0.002<br>0.022 | 585 | 0.039 |
| *SUSC_8* | TonB-dependent receptor | 1.555 | 3.131 | | | 17<br>825<br>948 | 0.977<br>0.971<br>0.981 | 742<br>825 | 0.011<br>0.012 | | |
| *SUSC_9* | TonB-dependent receptor | 1 | 3.872 | 394 | 0.025 | 394 | 0.990 | 10<br>80<br>93<br>394<br>660<br>741<br>927 | 0.034<br>0.032<br>0.033<br>0.037<br>0.015<br>0.042<br>0.015 | | |

| | | | | | | | | | |
|---|---|---|---|---|---|---|---|---|---|
| *SUSC_10* | TonB-dependent receptor | 13.936 | 0.803 | | | 249 | 0.981 | 73<br>249<br>379<br>670<br>840<br>951 | 0.049<br>0.044<br>0.028<br>0.037<br>0.031<br>0.042 |
| *SUSC_17* | TonB-dependent receptor | 7.122 | 0.734 | | | 523 | 0.995 | 523<br>982 | 0.016<br>0.008 |
| *SUSC_28* | TonB-dependent receptor | 1 | 5.925 | 996 | 0.045 | 996 | 0.984 | 87 | 0.044 |
| *SUSC_52* | TonB-dependent receptor | 1.373 | 11.945 | 84 | 0.0424 | 84<br>225 | 0.983<br>0.985 | 84 | 0.012 |
| *SUSC_62* | TonB-dependent receptor | 103.508 | 0.307 | 710<br>833 | 0.977<br>0.973 | 249 | 0.007 | | |
| *SUSD_2* | TonB-dependent receptor | 1.357 | 2.935 | 299 | 0.045 | 299 | 0.998 | | |
| *SUSD_3* | TonB-dependent receptor | 78.036 | 0.344 | | | 264 | 0.967 | 362 | 0.019 |
| *TABA* | hypothetical protein | 4.803 | 6.173 | | | | | | |
| *TAM* | Trans-aconitate 2-methyltransferase | 2.165 | 0.000 | 12<br>80<br>186 | 0.018<br>0.016<br>0.008 | 12<br>26<br>29<br>31<br>80<br>175<br>186 | 0.997<br>0.991<br>0.954<br>0.985<br>0.998<br>0.967<br>0.998 | 12<br>80<br>186 | 0.015<br>0.025<br>0.013 |
| *TCDA* | tRNA threonylcarbamoyladenosine dehydratase | 1.515 | 3.586 | | | 204 | 0.9515 | 120<br>194<br>204 | 0.0387<br>0.0171<br>0.0035 |

| | | | | | | | | | | | |
|---|---|---|---|---|---|---|---|---|---|---|---|
| *TETQ* | Tetracycline resistance ribosomal protection protein tet(Q) | | | | | 35<br>237<br>426 | 0.9676<br>0.9669<br>0.9623 | 354<br>550 | 0.003<br>0.0464 | | |
| *TETM* | Tetracycline resistance protein TetM from transposon TnFO1 | | | | | 35 | 0.964 | | | | |
| *THRS* | Threonine--tRNA ligase | 5.359 | 1.028 | 40 | 0.010 | 40 | 0.996 | 40 | 0.001 | | |
| *TIG* | Trigger factor | 1.231 | 6.139 | | | 81 | 0.983 | | | | |
| *TMOS_1* | Sensor histidine kinase | | | 111<br>1206<br>1215 | 0.005<br>0.010<br>0.017 | 111<br>684<br>781<br>1206<br>1215 | 0.999<br>0.983<br>0.965<br>0.998<br>0.998 | 111<br>1206<br>1215 | 0.009<br>0.017<br>0.026 | 1206 | 0.039 |
| *TMOS_2* | Sensor histidine kinase | 28.293 | 0.546 | | | 472 | 0.983 | 456<br>472<br>473<br>789 | 0.022<br>0.017<br>0.003<br>0.016 | | |
| *TMOS_4* | Sensor histidine kinase | | | 1252 | 0.026 | 555<br>1252 | 0.988<br>0.998 | 113<br>525<br>937<br>1252<br>1324 | 0.020<br>0.018<br>0.044<br>0.038<br>0.025 | | |
| *TMOS_5* | Sensor histidine kinase | | | | | 158<br>677<br>1120 | 0.983<br>0.974<br>0.985 | | | | |
| *TMOS_6* | Sensor histidine kinase | 1.219 | 24.151 | | | 292 | 0.961 | | | | |
| *TODS_1* | Sensor histidine kinase | 46.454 | 0.611 | | | 1244 | 0.972 | 792<br>820<br>1340 | 0.028<br>0.042<br>0.024 | | |
| *TODS_2* | Sensor histidine kinase | 30.002 | 0.530 | | | 472 | 0.9832 | 456<br>472<br>473<br>789 | 0.022<br>0.017<br>0.003<br>0.016 | | |

| | | | | | | | | | | | |
|---|---|---|---|---|---|---|---|---|---|---|---|
| *TODS_3* | Sensor histidine kinase | 11.914 | 2.390 | 16<br>729 | 0.025<br>0.009 | 16<br>141<br>146<br>543<br>547<br>627<br>729 | 0.995<br>0.951<br>0.952<br>0.956<br>0.982<br>0.960<br>0.998 | 16<br>500<br>729 | 0.037<br>0.000<br>0.001 | 729 | 0.026 |
| *TODS_4* | Sensor histidine kinase | 9.224 | 0.982 | 1252 | 0.026 | 555<br>1252 | 0.988<br>0.998 | 113<br>525<br>937<br>1252<br>1324 | 0.020<br>0.018<br>0.044<br>0.038<br>0.025 | | |
| *TODS_5* | Sensor histidine kinase | | | | | 158<br>677<br>1120 | 0.983<br>0.9734<br>0.985 | | | | |
| *TOLC_1* | Outer membrane protein | 4.695 | 2.724 | 436 | 0.043 | 436 | 0.979 | 8<br>199<br>421<br>436 | 0.022<br>0.009<br>0.046<br>0.0097 | | |
| *TOPA* | DNA topoisomerase 1 | | | 65 | 0.0224 | 65<br>735 | 0.996<br>0.951 | 65 | 0.009 | | |
| *TQSA* | AI-2 transport protein | 2.8 | 4.772 | | | 87 | 0.960 | 87<br>385 | 0.029<br>0.041 | | |
| *TRPB_1* | Tryptophan synthase beta chain | 10.301 | 0.654 | | | | | | | | |
| *TRPB_2* | Tryptophan synthase beta chain | 2.9 | 3.546 | | | 8 | 0.973 | 8<br>64<br>66 | 0.019<br>0.020<br>0.044 | | |
| *TRPD2* | Anthranilate phosphoribosyltransferase 2 | | | 267 | 0.021 | 267 | 0.993 | 267 | 0.004 | | |
| *TRPE* | Anthranilate synthase component 1 | 2.805 | 4.761 | | | | | 71 | 0.028 | | |
| *TRPG* | Anthranilate synthase component 2 | 24.276 | 2.354 | | | | | | | | |
| *TRUA* | tRNA pseudouridine synthase A | 5.081 | 1.775 | 216 | 0.974 | 135<br>216 | 0.049<br>0.041 | | | | |

| | | | | | | | | | | | |
|---|---|---|---|---|---|---|---|---|---|---|---|
| *TSAB* | tRNA threonylcarbamoyladenosine biosynthesis protein | 18.71 | 1.016 | | | | | | | | |
| *TUAA* | Putative undecaprenyl-phosphate N-acetylgalactosaminyl 1-phosphate transferase | 1 | 13.743 | | | | | | | | |
| *TUAB_1* | Teichuronic acid biosynthesis protein | | | | | 310 | 0.977 | | | | |
| *TUAB_3* | Teichuronic acid biosynthesis protein | | | | | | | | | | |
| *TUAD_1* | UDP-glucose 6-dehydrogenase | 2.039 | 5.038 | | | 43 | 0.971 | 43 | 0.036 | | |
| *UBIG* | Ubiquinone biosynthesis O-methyltransferase | 4.704 | 3.504 | 276 | 0.0228 | 276 | 0.993 | 128<br>276 | 0.035<br>0.002 | | |
| *UVRC* | UvrABC system protein C | | | 23 | 0.024 | 23 | 0.995 | 23<br>126 | 0.003<br>0.045 | | |
| *VALS* | Valine--tRNA ligase | 81.796 | 0.259 | 188 | 0.049 | 188 | 0.981 | 142<br>188<br>327 | 0.014<br>0.000<br>0.018 | | |
| *VIAA* | Protein ViaA | | | | | 280 | 0.962 | | | | |
| *WAAA* | 3-deoxy-D-manno-octulosonic acid transferase | 1.944 | 3.885 | | | | | | | | |
| *YBDG_2* | Miniconductance mechanosensitive channel | 46.264 | 0.124 | | | 19<br>338 | 0.985<br>0.971 | 19 | 0.002 | | |
| *YBHR_1* | Inner membrane transport permease | 2.616 | 10.098 | 83<br>95<br>304 | 0.000<br>0.018<br>0.020 | 83<br>95<br>304<br>399 | 0.999<br>0.993<br>0.995<br>0.984 | 83<br>95<br>292<br>304<br>353<br>399 | 0.000<br>0.027<br>0.021<br>0.005<br>0.038<br>0.028 | 83<br>95 | 0.000<br>0.010 |
| *YDCP_2* | Putative protease | | | 371 | 0.044 | 52<br>371 | 0.977<br>0.996 | 52<br>371 | 0.017<br>0.014 | | |
| *YDHD* | Putative sporulation-specific glycosylase | 4.680 | 5.147 | 415 | 0.013 | 150<br>222<br>415 | 0.973<br>0.971<br>0.994 | 84<br>213<br>222<br>415<br>419 | 0.007<br>0.031<br>0.027<br>0.026<br>0.029 | | |

| | | | | | | | | | | | |
|---|---|---|---|---|---|---|---|---|---|---|---|
| *YHES* | putative ABC transporter ATP-binding protein | 8.691 | 1.312 | 549<br>553 | 0.044<br>0.041 | 549<br>553 | 0.9546<br>0.9933 | 549<br>560 | 0.009<br>0.030 | | |
| *YHIM* | Inner membrane protein | 2.372 | 6.295 | | | 155 | 0.984 | 89<br>101<br>155 | 0.018<br>0.046<br>0.007 | | |
| *YIAD_2* | Putative lipoprotein | 4.89 | 7.861 | 1<br>23<br>281 | 0.014<br>0.0167<br>0.049 | 1<br>19<br>63<br>281 | 0.996<br>0.971<br>0.983<br>0.991 | 1<br>73 | 0.023<br>0.029 | 1 | 0.040 |
| *YIDC* | Membrane protein insertase | | | 75 | 0.005 | 75<br>277 | 0.999<br>0.961 | 75 | 0.001 | 75 | 0.045 |
| *YIGZ* | IMPACT family member YigZ | 6.537 | 5.209 | 198 | 0.011 | 198 | 0.996 | 35<br>198 | 0.029<br>0.017 | | |
| *YJAB* | putative N-acetyltransferase | | | | | 134<br>160<br>172<br>248<br>278 | 0.975<br>0.979<br>0.961<br>0.986<br>0.984 | | | | |
| *YJJP_2* | Inner membrane protein | 2.359 | 6.301 | | | 155 | 0.984 | 89<br>101<br>155 | 0.018<br>0.046<br>0.007 | | |
| *YKFC* | Gamma-D-glutamyl-L-lysine endopeptidase | 2.906 | 7.173 | 195 | 0.029 | 173<br>195 | 0.958<br>0.991 | 5<br>173<br>195 | 0.043<br>0.042<br>0.042 | | |
| *YKNY_1* | putative ABC transporter ATP-binding protein | 4.782 | 1.694 | | | | | | | | |
| *YKNY_2* | putative ABC transporter ATP-binding protein | 17.105 | 1.258 | | | 119 | 0.960 | | | | |
| *YKNY_4* | putative ABC transporter ATP-binding protein | 1.715 | 3.822 | | | 132 | 0.979 | | | | |
| *YKNY_5* | putative ABC transporter ATP-binding protein | | | 126 | 0.018 | 53<br>126 | 0.977<br>0.994 | 126 | 0.028 | | |

| | | | | | | | | | | | |
|---|---|---|---|---|---|---|---|---|---|---|---|
| *YKNY_6* | putative ABC transporter ATP-binding protein | 1.001 | 1.186 | | | 10<br>147 | 0.031<br>0.027 | 10<br>147 | 0.031<br>0.027 | | |
| *YKNY_8* | putative ABC transporter ATP-binding protein | 9.311 | 0.279 | | | | | | | | |
| *YKNZ_1* | putative ABC transporter permease | 5.173 | 3.419 | 76 | 0.04 | 67<br>76<br>161 | 0.962<br>0.990<br>0.975 | 67<br>76<br>161<br>389 | 0.041<br>0.033<br>0.011<br>0.033 | | |
| *YKNZ_2* | putative ABC transporter permease | | | 217<br>340 | 0.046<br>0.042 | 70<br>217<br>340 | 0.994<br>0.993<br>0.989 | | | | |
| *YKNZ_3* | putative ABC transporter permease | | | 19 | 0.024 | 19<br>630<br>706 | 0.991<br>0.981<br>0.952 | 19<br>722 | 0.036<br>0.029 | | |
| *YLMA* | putative ABC transporter ATP-binding protein | | | 253<br>430 | 0.027<br>0.027 | 163<br>253<br>397<br>430 | 0.954<br>0.991<br>0.981<br>0.992 | 253<br>430 | 0.039<br>0.004 | | |
| *YOAB* | Calcium-transporting ATPase 1 | | | 106<br>582 | 0.022<br>0.006 | 106<br>582 | 0.997<br>0.998 | 75<br>106<br>224<br>366<br>582 | 0.039<br>0.009<br>0.046<br>0.034<br>0.001 | | |
| *YPDA_1* | Sensor histidine kinase | | | 33 | 0.040 | 33<br>108 | 0.991<br>0.985 | | | | |
| *YPDA_2* | Sensor histidine kinase | | | | | | | 242 | 0.049 | | |
| *YQHD* | Alcohol dehydrogenase | 10.805 | 2.366 | 135 | 0.003 | 135 | 0.999 | 135<br>291 | 0.000<br>0.010 | 135 | 0.027 |
| *YRAA* | Putative cysteine protease | | | 69 | 0.0452 | 69 | 0.974 | | | | |
| *YRRB* | TPR repeat-containing protein | | | | | | | 566 | 0.009 | | |
| *ZWF* | Glucose-6-phosphate 1-dehydrogenase | 4.356 | 1.902 | | | 75<br>138 | 0.952<br>0.950 | 297 | 0.008 | | |
